# No-take marine reserves promote oligotrophic reef bacterioplankton communities across the Great Barrier Reef

**DOI:** 10.64898/2026.07.23.740257

**Authors:** Marko Terzin, Steven J. Robbins, Kim-Anh Lê Cao, Sara C. Bell, Katherine E. Dougan, Julian Zaugg, Renee K. Gruber, Michael J. Emslie, Daniela M. Ceccarelli, Samuel Chaffron, Philip Hugenholtz, Nicole S. Webster, David G. Bourne, Yun Kit Yeoh, Patrick W. Laffy

**Affiliations:** Australian Institute of Marine Science, PMB no3 Townsville MC, Townsville QLD 4810; College of Science and Engineering, James Cook University, Townsville, 4811; AIMS@JCU, James Cook University, Townsville QLD 4811; Australian Centre for Ecogenomics, School of Chemistry and Molecular Biosciences, The University of Queensland, St Lucia, QLD 4072; Melbourne Integrative Genomics and School of Mathematics and Statistics, University of Melbourne, Melbourne, Parkville VIC 3052; Nantes Université, École Centrale Nantes, CNRS, LS2N, UMR 6004, F-44000 Nantes, France; Research Federation for the Study of Global Ocean Systems Ecology and Evolution, FR2022/Tara Oceans GOSEE, F-75016 Paris, France; The University of Tasmania, Hobart 7005

**Keywords:** Marine Protected Areas, No-Take Marine Reserves, Ecosystem Resilience, Coral Reef Management, Microbial Communities, Network Analysis, Fishing Impact, Bacterial Metagenomes, Reef Health Indicators

## Abstract

Australia’s Great Barrier Reef is a biodiversity hotspot critical to ocean health, yet it faces increasing threats from climate change and localised impacts requiring effective conservation and management action. Rezoning of the Great Barrier Reef Marine Park in 2004 expanded No-Take Marine Reserves (NTMRs) to restrict extractive activities like fishing and collecting, creating one of the largest networks of marine reserves globally. Benefits like increased biomass of fisheries-targeted species and improved coral community health metrics have been reported, though the effects of zoning on water chemistry and seawater microbiology remain unexplored. Using data from the Great Barrier Reef Microbial Genomics Database, we investigated the structure of seawater microbiomes on 48 offshore reefs within NTMRs and fished reefs. A supervised classification method (MINT sPLS-DA) identified 350 indicator species that predict zoning with ∼71% accuracy (range 58–85%). Microbial communities broadly reflected reef states, with NTMR zones enriched in streamlined microbial oligotrophs (*Pelagibacter* and SAR86) correlating with higher cover of hard coral, crustose coralline algae, and herbivore fish abundance under lower nutrient conditions. By contrast, fished reefs harbored opportunists (Flavobacteriales, especially UA16, and Pseudomonadales) associating with elevated nutrients and turf algae cover. Co-occurrence networks revealed stronger competitive interactions in fished reefs, where nutrient-responsive taxa may outcompete other microbes, underscoring the need to investigate how these shifts influence reef nutrient cycling and function. Our findings reveal ecosystem-wide effects of marine zoning beyond fish protection, with distinct seawater microbiomes between fished reefs and NTMRs, which will help build decision tools for more targeted reef health monitoring assessments.

## Introduction

Marine Protected Areas, particularly No-Take Marine Reserves (NTMRs), represent conservation tools aimed at protecting exploited species and ecosystems through restricting extractive activities such as fishing and mining^1–4^. Recent estimates indicate that coral reefs within these marine reserves support significant recovery and maintenance of exploited fish biomass, accounting for an estimated 10% of the existing fish biomass in reefs globally^5^. Fisheries management success was also documented for Australia’s Great Barrier Reef (GBR) Marine Park which was rezoned in 2004, resulting in NTMRs increasing from ∼5% to ∼33% of the entire area^6^ and thereby becoming the world’s largest network of marine reserves at the time^7,8^. Benefits of rezoning accrued rapidly^7,9^, with coral trout (*Plectropomus* spp., *Variola* spp.) density increasing by 57–75% within two years^10^ and biomass by up to 89% within four years of rezoning^11^. NTMRs now support half of grouper biomass and 47% of fishery yield within the GBR Marine Park through spillover into adjacent fished areas^12^. In addition to supporting fish biomass recovery, NTMRs in the GBR also by displayed lower impacts and faster recovery from disturbances such as bleaching and cyclones^11,13^, in addition to showing more spatially heterogeneous and biodiverse benthic communities^14^, with fewer crown-of-thorns starfish outbreaks^15^ and reduced coral disease prevalence^16^. These GBR-specific trends align with global evidence demonstrating that NTMRs support healthier reef benthic communities, with higher cover of hard coral and crustose coralline algae (CCA), increased fish diversity and biomass, and reduced macroalgal cover, all of which may contribute to faster recovery rates^17,18^ and enhanced reef resilience against climate change and other disturbances^19,20^.

The observed reef health benefits within NTMRs including enhanced fish biomass across trophic levels and shifts in coral versus algal dominance, may also alter nutrient dynamics as benthic primary producers (e.g. corals and algae) differ in the type and quantity of dissolved organic matter they release^21–23^, and fish communities at different trophic levels excrete and egest nutrients with distinct N:P stoichiometries^24,25^. This can collectively modify the quantity, composition, and stoichiometry of dissolved and particulate nutrients entering the surrounding reef water. Because seawater microbial communities respond rapidly to changes in water chemistry, nutrient fluxes, and organic matter inputs^26–29^, protection-driven shifts in reef ecosystems may also restructure microbial community composition and function, with potential consequences for microbially mediated ecosystem services like nutrient/biogeochemical cycling and productivity^30^ yet these processes remain poorly understood in the Great Barrier Reef and coral reefs globally. Currently, only a few meta-omics studies have explored how reef protection affects microbial dynamics of the surrounding bacterioplankton, but emerging evidence suggests: (1) NTMRs in Brazil’s Abrolhos Bank supported distinct seawater microbial communities alongside higher fish biomass and lower macroalgal cover^31^ and (2) in Kuwait’s Sulaibikhat Bay, protected sites showed higher microbial diversity correlated with elevated nitrogen, phosphorus and salinity levels compared to fished areas^32^. However, in both studies, protected and unprotected sites were located far apart and thus the effects of protection status remain challenging to isolate from inherent environmental differences, and may be partially confounded by the offshore placement of protected reefs.

Reef protection measures may influence seawater microbial communities by reducing human impacts and promoting more diverse bacterioplankton communities, however, more targeted research is needed to clarify the relationship between reef zoning and the health and functioning of seawater microbes. To address this gap, we examined the distribution of 5,283 prokaryotic metagenome-assembled genomes (pMAGs) assembled from 48 offshore reef-associated seawater metagenomes from the Great Barrier Reef Microbial Genomics Database (GBR-MGD)^33^ (**Fig. 1**). Here, we compared co-located fished and NTMR reefs to identify microbial indicators of reef zoning status, and clarify their relationships to protection status through associations with physico-chemical variables (temperature, salinity, chlorophyll *a*, and dissolved/particulate nutrients), benthic cover (coral types, algae, and abiotic components) and fish abundances and biomass. Using supervised machine learning, microbial niche modeling, and network analysis, we assessed how reef zoning shapes reef bacterioplankton composition, functioning, and microbe-to-microbe interactions, identifying seawater microbial signatures that could inform future monitoring.

**Figure 1.**
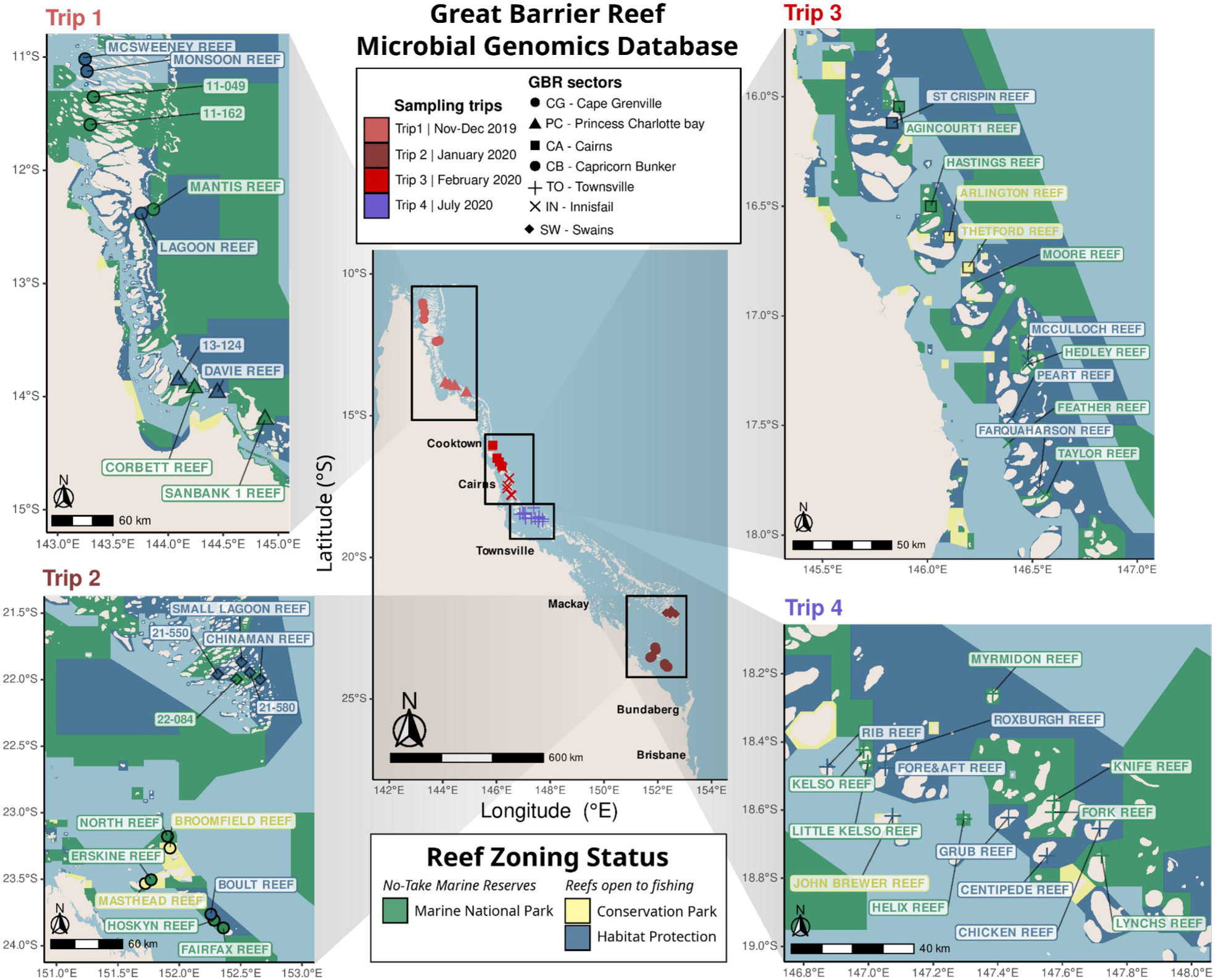
Sampling scheme used in the Great Barrier Reef Microbial Genomics Database (GBR-MGD). Microbial metagenomes and physico-chemical measurements were obtained from surface seawater samples collected at 48 offshore reefs across the GBR, concurrently with AIMS-LTMP *in situ* estimates of benthic cover and fish abundance and biomass. Sampling occurred in four trips between November 2019 and July 2020, with red tones indicating Austral summer/wet season (trips 1-3, Nov 2019–Feb 2020) and blue indicating winter/dry season (trip 4, July 2020). Samples were taken across seven GBR sectors, denoted on the maps with different symbols. Trip-specific map insets show that reefs in No-Take Marine Reserves (NTMRs, green zones) and fished reefs (dark-blue and yellow zones) were sampled in pairs to minimise confounding effects of geography.

## Results and Discussion

### Stable seawater indicators of GBR zoning status across season and geography

No-Take Marine Reserves (NTMRs) across the GBR have well-documented benefits for reef macroorganisms^7,12,13^, however, less is known about how reef zoning influences surrounding bacterioplankton communities. Community-level visualisation (PCA) and statistical testing (PERMANOVA and dbRDA) showed that reef protection status explained a small but statistically significant proportion of community variation (1.9%, p < 0.001), though sampling trip (58.6%), reef site (23.8%), and GBR sector (9.7%) had substantially larger influences (**Fig. 2A**; **Figs. S1-4; Table S1**). Having established this community-level signal, we then assessed which microbes are consistently indicative of NTMRs and fished reefs across the GBRMP. Zoning status was consistently predicted across 48 offshore reefs spanning seven sectors of the GBR (**Fig. 2**), with an average classification accuracy of ∼71% (**Fig. 2B**; **Figs. S10-13; Table S3-5**), significantly higher than expected by random chance (permutation test: p = 0.001; **Fig. S14; Table S6**). This was achieved using a subset of 350 indicator pMAGs (**Fig. 2C**) identified through MINT sPLS-DA^34–36^ as stable microbial markers of reef zoning that were spatially and temporally reproducible (**Fig. 2D; Fig. S12**).

**Figure 2.**
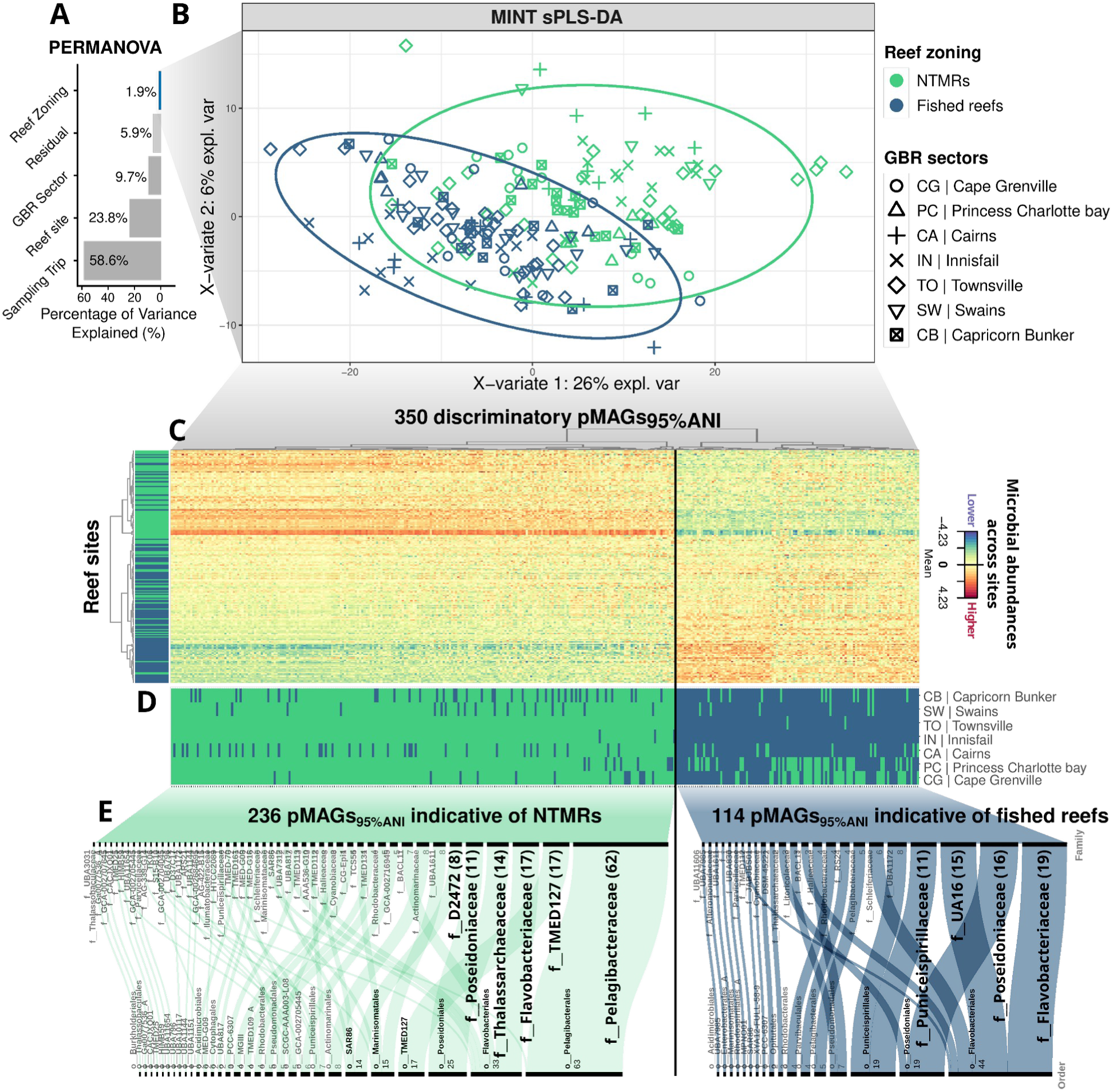
Visualization of seawater microbes selected to distinguish NTMRs from fished reefs across seven GBR sectors. (**A**) Proportion of variance in seawater microbial communities explained by different factors as inferred by the PERmutational Multivariate ANalysis Of VAriance (PERMANOVA). (**B**) Sample plot from MINT sPLS-DA showing clustering of reefs by reef zoning (fished vs NTMRs) based on indicator seawater prokaryotic metagenome-assembled genomes (pMAGs). Samples (48 reefs x four replicates) are projected in the first two components of the MINT sPLS-DA space, with ellipses indicating 95% confidence level. (**C**) Heatmap shows differential abundances for the pMAGs (columns) discriminating between NTMRs and fished reefs (rows). Euclidean clustering of microbes by abundance across reef sites splits the heatmap into two groups: 236 pMAGs indicating NTMRs (left), and 114 microbial indicators of fished reefs (right). (**D**) Heatmap indicating whether the pMAGs were more relatively abundant in NTMRs (green) or fished reefs (blue) in each GBR sector. (**E**) Alluvial diagrams summarise the taxonomy of indicator microbes for NTMRs (left) and fished (right) reefs, ordered by the most common taxa for each zone.

Taxonomic analysis of these indicators revealed consistent differences in microbial assemblages between NTMRs and fished reefs. Of the 350 indicator pMAGs, 236 were consistently enriched in NTMRs (**Fig. 2C-E**, green) with 63 belonging to the order Pelagibacterales (of which 62 were Pelagibacteraceae, 44/63 *Pelagibacter*). Additional indicator taxa were classified within orders TMED127 (in the class Alphaproteobacteria) and Flavobacteriales, with families TMED127 (including genus GCA-002690875) and Flavobacteriaceae being the second most abundant indicator taxa of NTMRs, each with 17 pMAGs. The orders Poseidoniales (25 indicators), Marinisomatales (15 indicators) and SAR86 (14 indicators) were other notable taxa indicative of NTMRs. In contrast, 114 pMAGs were consistently more abundant on fished reefs compared to NTMRs (**Fig. 2C-E**, blue). Of these, 44 indicators belonged to the order Flavobacteriales with 19 classified as Flavobacteriaceae (10 of which were classified as genus *Arcticimaribacter*, making this the most numerically abundant genus level indicator alongside UBA11663 in the family Sanyastnellaceae), and 16 from the family UA16 (at genus level: 10 from UBA11663, 2 from UBA8752, and one not identified to genus level) (**Fig. 2E**, blue). Other notable taxa indicative of fished reefs included 19 members of the archaeal order Poseidoniales (6 each belonging to the genera *Poseidonia* and MGIIa-L1), 18 in the order Puniceispirillales, 7 in the order Pseudomonadales and 4 in the Parvibaculales.

The average classification accuracy differed between reef zones (**Table S4**) and GBR sectors (**Fig. S12; Tables S3-5**). NTMR reefs were more accurately predicted compared to fished reefs (75% vs 68% accuracy; **Table S4**), a difference that may reflect variation in protection levels among the sampled fished reefs. Specifically, 20 reefs were sampled in the less protected dark blue ‘Habitat Protection’ zone, where most fishing is allowed except trawling, versus five reefs in the more protected yellow ‘Conservation Park’ zone^37^ (**Fig. 1**). While finer-scale comparisons of protection levels (e.g., green vs. dark blue vs. yellow zones) could offer additional insights, the limited number of reefs sampled in the yellow zone (n=5) precluded robust statistical comparisons at this granularity.

Among sectors, the highest prediction accuracy at 85% was detected in the Innisfail sector followed by Princess Charlotte Bay (80%), while the lowest were observed for Cape Grenville and Swains sectors (50% and 55%, respectively; Table S5), which may be attributed to particular characteristics of these sectors. In Cape Grenville, reefs are located close to the coast and subject to influences from the Torres Strait and northern Australia, including the presence of fish species with northern-truncated distributions that extend from the north and north-west Western Australia^38^. Additionally, hydrodynamic characteristics in Cape Grenville differ from those further south due to a predominant northerly flow from incoming Coral Sea currents north of Cape Flattery^39,40^, and fishing pressure is lower in the Far-North GBR due to its remoteness^41^. Poor zoning prediction within the Swains is likely also due to the remoteness of this sector, with reefs situated between 150 and 250 km from the coast, further offshore than any other GBR reefs and therefore subjected to minimal coastal influence^7^. Fish and benthic community composition are also somewhat distinct in the Swains compared to other GBR regions^38,42^, and oceanographic processes including southerly transport from the East Australian Current lagoonal branch and frequent upwelling events^43^ may have contributed to distinct environmental conditions and thus poor zoning predictions in the Swains.

Despite these sector-specific variations, our study demonstrates the potential of seawater microbes as stable and consistent indicators of reef zoning across the GBR (**Fig. 2D; Figs. S10–13**), with distinct microbial taxa associated with NTMRs and fished reefs, highlighting their utility for monitoring zoning status across broad spatiotemporal scales. Critically, convergence across multiple independent analytical approaches strengthens confidence in these indicators: while MINT sPLS-DA is the most appropriate framework for multi-study integration across sectors (as described in *Methods*), Random Forest (58.9% feature overlap with MINT; **Fig. S15**) and ALDEx2 (83.7% indicator overlap; **Figs. S16–S20**) nonetheless identified the same taxa as enriched in the same reef zone, with perfect agreement in enrichment direction across methods. Further, presence/absence analysis demonstrated near-universal detection of zoning indicator pMAGs across sites (>90%; **Fig. S21**), showing these taxa are widespread rather than detection-limited, a key consideration given potential genome size effects on abundance estimates (i.e. larger genomes accumulate more reads in the mapping step). Additionally, read-based metagenomic analysis corroborated these zoning signatures (**Fig. S22**) without requiring genome assembly or binning, further supporting practical monitoring applications where reduced sequencing and computational effort (comparatively to MAG-centric metagenomics) are advantageous for more rapid monitoring.

### Linking environmental variables with indicator microbes

Environmental drivers that shape the differences in microbial assemblages between NTMRs and fished reefs were identified through an integrated analysis of reef bacterioplankton and ecosystem conditions at these offshore GBR sites. MINT sPLS^36,44,45^ analysis identified that microbial indicators enriched in NTMRs were more abundant and correlated with environmental variables indicative of healthy reefs^46^, including elevated fish biomass, increased coral and crustose coralline algae (CCA) cover, reduced turf algae, and lower nutrient concentrations (**Fig. 3A–B**; **Table S7; Supplementary Data 5**). In contrast, microbial indicators of fished reefs were enriched under environmental features characteristic of degraded reef conditions^22,23^, including lower fish abundances, high turf-to-coral ratios, and nutrient enrichment (**Fig. 3A–B**; **Table S7; Supplementary Data 5**). Interestingly, even though the MINT sPLS approach is unsupervised and does not incorporate reef zoning information, it nevertheless revealed distinct clustering of microbial communities by zoning, with NTMR reefs (e.g., Myrmidon, Moore, and Hastings reefs) clustering separately from fished sites (e.g., Farquharson, John Brewer, and Masthead reefs), independent of reef sector (**Fig. 3A**).

**Figure 3.**
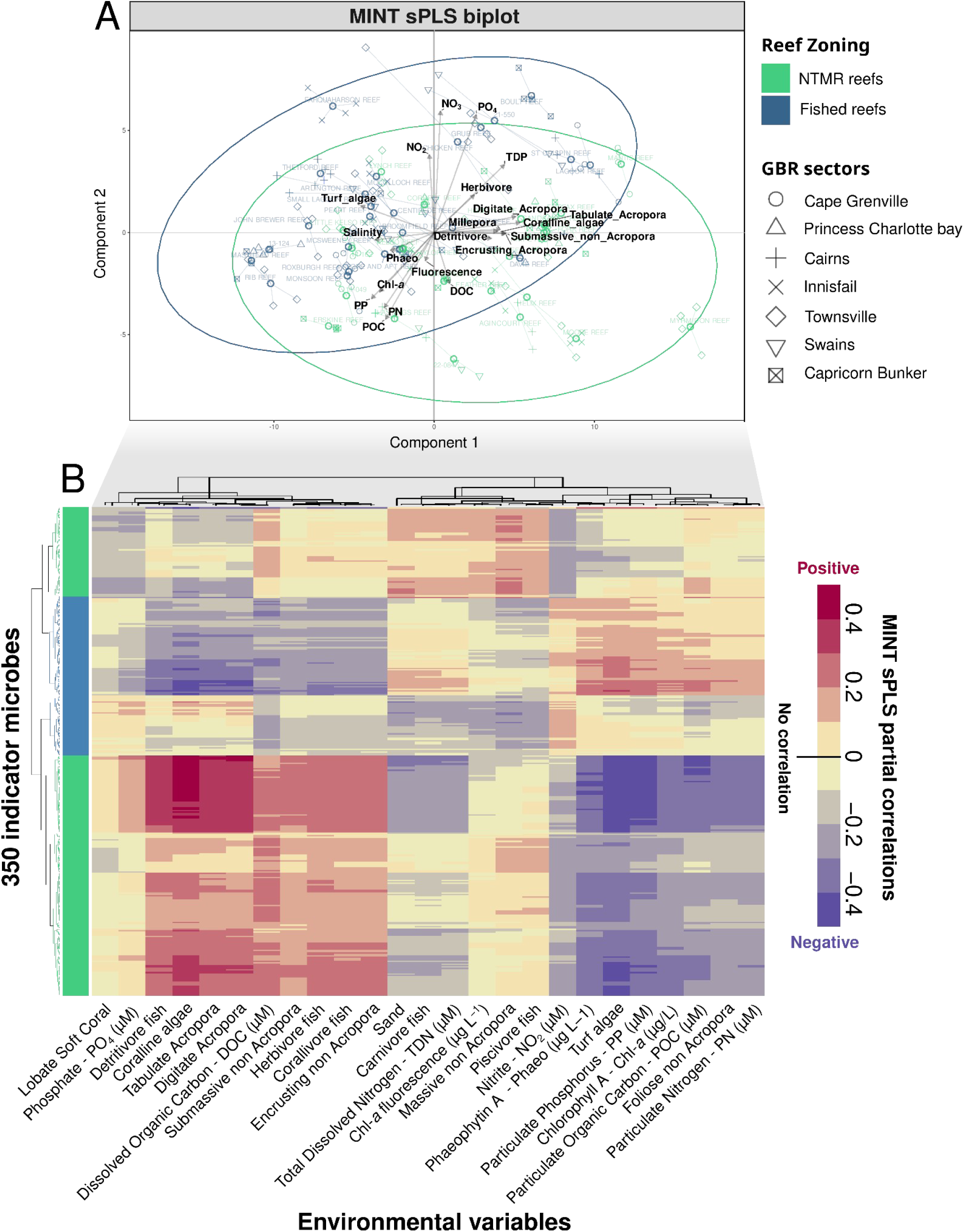
MINT sPLS integrates abundances of seawater microbes with continuous environmental data, while accounting for GBR sector-specific variation. (A) The biplot from MINT sPLS shows both samples (four replicates per reef; hollow circles represent site centroids connecting the four site replicates) and variables (environmental only, in black; microbial variables omitted for clarity) on the same plot. (B) A heatmap shows the MINT sPLS partial correlations between 350 pMAGs identified in MINT sPLS-DA as reef zoning indicators (rows; colored based on indicator status) and 25 most influential environmental variables (columns).

The establishment of NTMRs in 2004 increased fish biomass in the GBR^10–12^, a trend also reflected in our data, where biomass was approximately 1.28× greater in NTMRs (mean 65.57 ± S.E. 73.55 kg 1000 m^-2^) than in fished reefs (mean 51.28 ± S.E. 41.34 kg 1000 m^-2^; z=2.371, p=0.018; **Fig. S23A; Tables S8–S10**). While herbivorous fish densities (**Fig. S23B; Tables S11–S14**) and the cover of hard corals (**Fig. S23C; Tables S15–S18**) were not significantly different between NTMRs and fished reefs across all sites, fish abundances (including herbivores) and overall biomass were collinear and elevated together with CCA and hard coral cover in a subset of NTMR reefs (**Fig. 3A**; component 1), explaining why NTMR-associated microbial indicators were positively linked to all these variables simultaneously (**Fig. 3B**). Specifically, the 236 indicators enriched in NTMRs (for taxonomic annotations, refer to **Fig. 2E**, green) were positively associated with higher abundances of multiple fish groups including detritivores, corallivores, herbivores, invertivores, and piscivores (**Fig. 3B**; green cluster; **Table S7; Supplementary Data 5**). These NTMR-enriched microbes also showed positive relationships with hard coral cover, including different groups of *Acropora* (digitate, tabulate and encrusting), CCA and the hydrozoan *Millepora,* and were negatively associated with turf algae cover (**Fig. 3B**; green cluster; **Table S7; Supplementary Data 5**). Fish and coral are likely collinear because corals enhance reef 3D complexity to support diverse fish assemblages^47–50^. Additionally, elevated counts of herbivorous fish in some NTMR reefs (**Fig. 3A**; component 1) may have resulted in qualitative differences in grazing pressure and increased algae removal in NTMRs, thereby creating more space for coral and CCA to settle^38,51,52^.

Microbes enriched in NTMRs were also correlated with reduced nutrient concentrations, including particulate organic matter (POM) and dissolved inorganic nitrogen (specifically NO_2_ and NO_3_, collectively termed NO_x_), while showing a positive association with dissolved organic carbon (DOC) (**Fig. 3A–B**). The negative associations between coral cover and water column nutrients (POM and NOx concentrations, in addition to lower Chl-*a* and Phaeo; **Fig. 3A–B**) may reflect nutrient assimilation by coral and other benthic organisms^53^ in addition to reduced POM remineralisation into NOx due to the lower POM concentrations^54^. Corals supplement their nutrition through heterotrophic feeding on phytoplankton, zooplankton, and detritus^53^, with literature proposing that this efficient uptake of nutrients (including NO_x_) maintains the high productivity of reef ecosystems despite residing in nutrient-poor waters^29^. Viewed through the lens of the reef “microbialization” hypothesis, lower microbial depletion of DOC in NTMRs and the fact that DOC there likely originates from hard coral cover (given its collinearity with tabulate and digitate Acroporans and non-*Acropora* groups; **Fig. 3B**), suggests that microbial activity may be more tightly coupled to particulate and inorganic nutrient pools than to labile DOC. This is consistent with coral-dominated systems where DOC production is lower and microbial respiration is less stimulated than on algal-dominated reefs, where enhanced microbial respiration of DOC exudates from algae (including turfs) results in DOC drawdown^22^. Overall, nutrient cycling appears to be more efficient in protected coral-dominated reefs where higher benthic and fish biomass efficiently removes or limits the accumulation of organic particulates^55^. This aligns with findings from highly protected, near-pristine reefs like Jardines de la Reina, Cuba^56^, where the most strictly enforced protection zones resulted in some of the highest fish biomass and coral cover in the Caribbean^57^, maintaining oligotrophic conditions and picoplankton microbial communities with the highest alpha diversity and dominated by oligotrophic taxa like SAR11 and *Prochlorococcus*, despite geographic factors (i.e. site-specific effects) being the primary driver of community variation^56^.

Conversely, fished reefs with reduced herbivore counts may experience diminished grazing^58^, facilitating turf algae proliferation^22,23,59^. While turf algae cover was not significantly different between NTMRs and fished reefs when considering all sites (**Fig. S23D**; **Tables S19–22**), it was significant in some reefs (**Fig. 3A**, component 1), explaining why the 114 microbial indicators of fished reefs positively associate with these metrics collectively (**Fig. 3B**, blue cluster; **Table S7; Supplementary Data 5**). With respect to nutrient dynamics, microbial indicators of fished reefs were negatively correlated with DOC, suggesting enhanced microbial uptake of labile DOC in reefs open to fishing, likely driven by heterotrophic microbial processes and potentially indicative of early stages of reef microbialization^22^. DOC uptake may fuel microbial biomass production, potentially contributing both to POM accumulation and the release of NO_x_ (as a byproduct of microbial metabolism) into the water column. To explore the hypothesis that POM in reefs open to fishing is partly of microbial origin, we examined POC:PN ratios relative to the Redfield ratio, a well-established benchmark in oceanography describing the canonical C:N composition of marine phytoplankton^60^. The average POC:PN ratios were close to the Redfield ratio of 6.6, suggesting a largely phytoplanktonic origin of POM across all reefs, indicating that particulate organic matter in offshore GBR reefs is primarily produced *in situ* rather than derived from external detrital inputs^61^, consistent with relatively tight nutrient recycling. However, the wider range in POC:PN values observed in fished reefs (**Fig. S24**) suggests greater variability in POM composition. This could reflect more site-specific or diverse POM inputs in fished areas (i.e., some sites had N-enriched POM, consistent with microbially processed or more labile detritus), potentially due to altered trophic dynamics and reduced detritivore fish abundance (**Fig. 3B**), which may affect detritus consumption rates^24^. As further evidence for enhanced microbial remineralisation in fished reefs, higher DIN:DIP ratios were observed in fished reefs (9.9 ± 7.9) relative to NTMRs (7.2 ± 5.5; **Fig. S25**), suggesting more efficient nitrogen utilisation in NTMRs vs. enhanced release and retention of NO_x_ in fished reefs. While the exact cause of this pattern remains uncertain, it could reflect both bottom-up (nutrient availability) and top-down (fishing pressure) effects on nutrient dynamics.

Our results provide support that marine microbes can be used as indicators of benthic-pelagic coupling in reef ecosystems, with the synergistic effects of reduced herbivory, declining hard coral cover, and increased turf algae collectively altering nutrient cycling and driving divergent microbial assemblages between some NTMRs and fished reefs. Most notably, the influence of these benthic-pelagic couplings are measurable in the microbial data despite subtle changes in individual parameters, confirming the proposed sensitivity of seawater microbes to integrated ecosystem states^27^. Given that microbial indicators proved informative even under the GBR’s relatively low fishing pressure when compared to other reef ecosystems globally^37,41^, their utility may be even greater in reef ecosystems where fishing is a dominant driver of ecosystem degradation. Such systems could leverage microbial monitoring as a sensitive, early-warning tool for changing trophic cascades and benthic community shifts.

### Microbial indicators prevalent in NTMRs harbor signatures of genome streamlining via gene loss

NTMR-enriched microbes exhibited genomic features consistent with microbial streamlining, an adaptation that facilitates survival in nutrient-poor pelagic systems^62–64^. Specifically, NTMR-enriched microbes such as Pelagibacterales, SAR86, and Marinisomatales (see **Fig. 2E**, green) had significantly (p < 0.0001) smaller genomes (∼1.57×; **Figs. 4A-B**) and lower GC content (∼1.62×; **Figs. 4C-D**) compared to microbes prevalent in fished reefs, which were dominated by Flavobacteriales UA16 and Schleiferiaceae (**Fig. 2E**, blue). These genomic signatures are consistent with adaptation to oligotrophic conditions^62^ and align with the lower nutrient concentrations measured in NTMRs (e.g. ∼1.50–1.71× lower NO_2_ and NO_3_; **Fig. S26**; **Table S23**). NTMR indicator pMAGs additionally exhibited ∼1.53× lower average KEGG module completeness compared to the 114 microbes indicative of fished reefs (**Figs. 4E-F**), consistent with gene loss and reduced metabolic capacity, while microbial indicators enriched on fished reefs appear to have an increased pathway completeness and thus may be more metabolically independent^65^.

**Figure 4.**
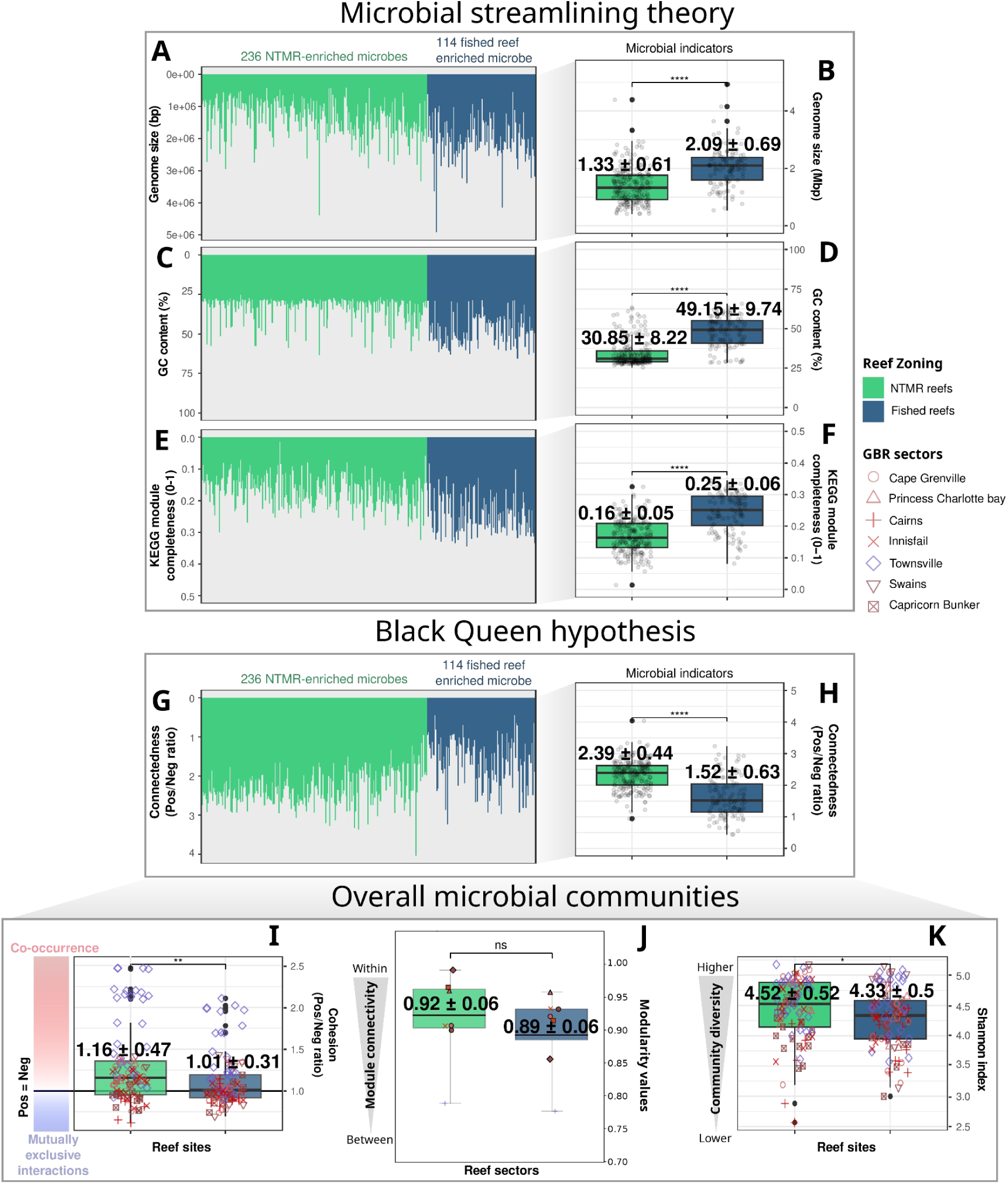
Genomic, functional, and network characteristics of indicator microbes associated with NTMRs (green) and fished reefs (blue). Genomic features (**A–D**): Barplots show genome size (**A**) and GC content (**C**) for indicator pMAGs, with boxplots comparing their distributions between NTMRs and fished reefs (**B, D**). Lower values are considered signatures of genome streamlining. Functional potential and interaction structure (**E–I**): (E) Barplots show average KEGG module completeness per pMAG, with (**F**) corresponding group-level comparisons. Lower values indicate gene loss and incomplete pathways. (**G**) Microbe-specific ratios of positive to negative co-occurrence edges represent cooperative versus antagonistic interactions, and (**H**) shows group-level comparisons of these ratios. (**I**) Sample-level ratios of positive to negative cohesion illustrate co-occurrence within overall bacterioplankton communities. Higher positive to negative ratios (both for connectedness and cohesion; **G-I**) indicate a prevalence of positive interactions, while lower values are a proxy of negative interactions. Diversity and community structure (**J–K**): (**J**) Comparison of modularity scores between NTMR and fished reefs computed for each sector × zone combination (n = 14 microbial co-occurrence networks). Higher modularity values indicate increased community connectivity within modules (greater compartmentalization), while lower values suggest higher between-module connectivity. (**K**) Alpha diversity (Shannon index) is shown for seawater microbiomes from NTMR and fished reef sites. For boxplots (**B, D, F, H, I, K**), significance levels from Wilcoxon rank sum tests are indicated as: *p < 0.05; **p < 0.01; ***p < 0.001; ****p < 0.0001; “ns” = not significant.

A key consequence of genome reduction and gene loss is that many streamlined microbes lose the ability to synthesise certain essential metabolites, rendering them dependent on neighboring community members for these compounds^66^. Cooperative interactions among free-living microbes are central to the Black Queen Hypothesis^67^, which posits that oligotrophic microbes will form more positively connected communities due to enhanced mutualistic metabolic exchanges^64,67,68^. Using species co-occurrence network analysis^69^, we found that NTMR indicator pMAGs with streamlined genomes were indeed ∼1.61× more positively connected to the broader community (**Figs. 4G–H**, green). In contrast, fished reef microbial indicators showed lower positive-to-negative connectedness ratios (**Figs. 4G–H**, blue), potentially implying more antagonistic microbial interactions such as competition, amensalism, or parasitism^70^. Supporting the predicted genomic underpinnings of these network patterns, regression analysis revealed significant (albeit weak) negative correlations between positive-to-negative connectedness and genome size (β = –2.67 × 10^-7^, R^2^ = 0.096, p < 0.001), GC content (β = –0.021, R^2^ = 0.143, p < 0.001), and KEGG module completeness (β = –0.265, R^2^ = 0.078, p < 0.001) (**Fig. S27**). This further supports the patterns that genome streamlining (**Figs. 4A–D**), marked by gene loss and reduced metabolic capacity (**Figs. 4E–F**), is linked to increased cooperative microbial interactions in NTMRs (**Figs. 4G–H**).

Fished reefs showed stronger signatures of antagonistic microbial interactions across entire reef bacterioplankton communities, with co-occurrence networks (inferred from all 876 pMAGs) exhibiting ∼12.9% lower positive:negative cohesion ratios (though with high variability; ±50% S.E.; **Fig. 4I**) compared to NTMRs. This patterns was consistent across GBR sectors (**Fig. S28**) suggesting a systemic shift toward negative (i.e., mutually exclusive) interactions in fished reef seawater microbiomes. As reef bacterioplankton indicators of fished reefs were shown to increase in prevalence with elevated nutrient concentrations (NO_2_^-^, NO_3_^-^, and POM; see **Fig. 3B**), this microbial competition may be linked to competition for available nutrients. Correspondingly, negative microbe-to-microbe interactions were also more prevalent during the austral summer (**Fig. S29**) when all nutrients apart from phosphate were elevated^71^ (**Fig. S30**). These observations align with previous findings from oligotrophic reefs in Porto Seguro (Bahia, Brazil) showing that reefs further from a river mouth and exposed to less organic pollution harboured more complex microbial networks, with a higher proportion of positive associations in the more pristine oligotrophic reefs^68^. The increased microbial positive connectedness in NTMRs (**Figs. 4G–I**) may also be the reason why we detected a higher number of indicator microbes for NTMRs than for fished reefs (236 vs 114 markers; **Fig. 2**), as more co-dependent microbes are stably associated and thus more consistently enriched, potentially also facilitating better predictions of NTMRs than fished reefs (∼7% increase in classification accuracy; **Table S4**). In contrast, the enrichment of pMAGs on fished reefs may reflect competitive dominance at the expense of other microbes, likely contributing to the ∼5% lower overall microbial diversity observed relative to NTMRs (**Fig. 4K**, consistent across most GBR sectors; **Fig. S31**). This is in line with findings by Hernandez et al. (2021)^70^ that negative microbial interactions strongly predict reduced microbial diversity.

Graph theory is increasingly used to quantify stability in microbial co-occurrence networks^72,73^. Recent frameworks^70^ suggest that healthy microbial communities show stronger partitioning (high connectivity within network modules, i.e., tightly linked microbial subgroups), which limits environmentally induced shifts within these modules. In contrast, unstable microbial communities (undergoing stress) are predicted to exhibit higher between-module connectivity which was proposed as a metric of destabilised networks^70^, as environmental fluctuations may propagate widely across the community, potentially resulting in network collapse if critical thresholds are exceeded^72^. Higher modularity and a greater proportion of within-module (versus between-module) edges therefore both indicate a more compartmentalised, and potentially more stable, network^74^. To test whether the elevated positive:negative cohesion we observed in NTMRs (**Fig. 4I**) corresponded to such a difference in compartmentalisation, we compared both metrics across the 14 sector×zone co-occurrence networks. Neither differed significantly between zones: modularity was marginally higher in NTMRs but not significantly so (**Fig. 4J**), and both the within-module edge fraction and the within:between edge ratio were near-identical between zones (p = 0.54; **Fig. S32; Table S24**). We therefore do not interpret the higher positive cohesion and connectedness in NTMRs as evidence of greater network stability, but instead as a shift in the balance of putatively cooperative versus competitive associations. Whether these contrasting microbial interaction structures carry community stability consequences may become clearer on more degraded reefs, as our NTMR and fished sites differed only modestly in reef-health metrics. Further experimental validation, such as controlled manipulation of nutrient levels in mesocosms coupled with microbial interaction assays or stable isotope probing to track metabolic exchanges, is required to explore this hypothesis.

### Microbes enriched on fished reefs are functionally primed for rapid carbohydrate uptake, energy production, and biosynthesis of complex compounds

To test if microbial indicators of fished reefs possess metabolic traits to compete for nutrients, hypothesised based on their associations with elevated nitrate, nitrite, and POM (**Fig. 3**), and network properties indicating microbial antagonism (**Fig. 4**), we compared the completeness of metabolic pathways (KEGG modules) between fished-reef and NTMR microbial indicators. Microbial indicators of fished reefs overall showed an enhanced potential for rapid nutrient utilisation as they had higher completeness in several metabolic pathways involved in carbohydrate metabolism, energy generation, and anabolic reactions, including biosynthesis of cofactors, vitamins, amino acids, and lipids (**Fig. 5**; Figs. S34-S40). This patterns appears to reflect true biological variation, as genome completeness estimates (Fig. S33) and reference genomes were consistent with predicted genome size (and associated metabolic capacity) variation.

**Figure 5.**
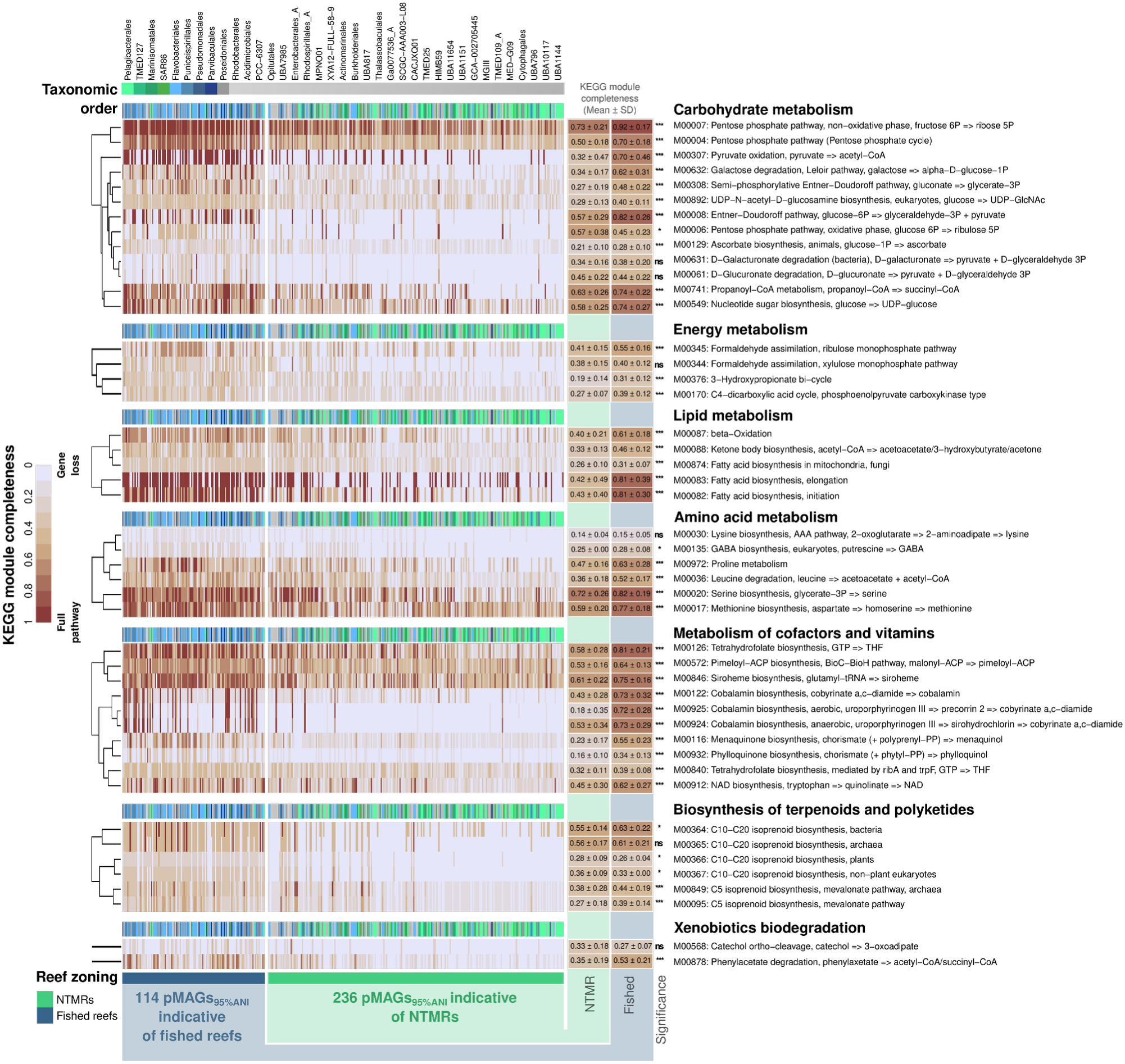
Differential metabolic potential of microbial indicator taxa in fished reefs and No-Take Marine Reserves (NTMRs). Heatmaps show the top 45 KEGG modules with the greatest differences in completeness scores between microbial indicator pMAGs from fished reefs and NTMRs. Rows represent KEGG modules (grouped by metabolic category), and columns represent indicator pMAGs clustered by reef zoning status. Module completeness scores (ranging from 0 to 1) are shown by color scale. Columns are annotated by the taxonomic order of each pMAG; only the four most abundant orders in each group are distinctly colored (green for NTMR-enriched, and blue for fished reef-enriched microbes; grey for others). Statistical differences in KEGG module completeness (mean ± SD) between groups were assessed using Wilcoxon rank-sum tests, with adjusted p-values indicated as: *p < 0.05; **p < 0.01; ***p < 0.001; ****p < 0.0001; “ns” = not significant.

Compared to NTMR-enriched pMAGs (e.g., Pelagibacterales, SAR86; **Fig. 2E**, green), the 114 microbes prevalent on fished reefs (primarily Flavobacteriales; **Fig. 2E**, blue) had more complete pathways for sugar metabolism, including the pentose phosphate, Entner-Doudoroff, and galactose degradation pathways (**Fig. 5**; **Fig. S34**). This may be reflective of ecological changes between NTMRs and fished reefs (though minor in our data) influencing the quantity and composition of organic matter in reef waters, potentially creating conditions that favour microbial assemblages capable of exploiting more labile and diverse carbon sources in fished reefs. Fished-reef microbial indicators likely process sugars derived from benthic algae, inferred from the collinearity between nutrients and turf cover (**Fig. 3A–B**) and the fact that our sampling sites are located on oblique or exposed reef slopes where seagrass is largely absent, although other organic matter sources (e.g., phytoplankton, detritus) may also contribute. Future work (e.g., isotopic tracing) will be necessary to confirm nutrient provenance and disentangle the relative contributions of these potential sources.

Microbes enriched on fished reefs also showed enhanced energy-generating capacity (**Figs. 5, S35**), with more complete carbohydrate utilisation pathways (M00004, M00308, M00008, M00631, M00632) converging on glyceraldehyde-3-phosphate for glycolysis^75^. Alongside enhanced acetyl-CoA synthesis (pyruvate oxidation M00307; β-oxidation M00087; **Fig. 5**), these pathways likely fuel the tricarboxylic acid cycle for ATP generation^75^, suggesting fished-reef seawater microbial assemblages are restructured for rapid energy harvest from organic substrates. This metabolic configuration supports their shift toward anabolic pathways, including the biosynthesis of lipids (**Fig. 5, S36**), amino acids such as serine, methionine, proline, and tryptophan (**Fig. 5, S37**), cofactors like vitamin B12 (cobalamin; **Fig. 5, S38**), and other secondary metabolites (**Figs. S39-S40**). Through *de novo* synthesis of essential macromolecules, fished-reef microbial indicators may exhibit enhanced metabolic independence^65^ providing a mechanistic explanation for the competitive advantage of fished-reef microbial indicators over other microbes (**Fig. 4G-I**). For example, the Entner–Doudoroff (ED) pathway, which is often more prevalent under elevated nutrient conditions, and the pentose phosphate pathway provide mechanisms by which dissolved organic carbon can be remineralised more quickly and less efficiently than through the Embden–Meyerhof–Parnas (glycolytic) pathway, which is more common in healthier coral-dominated reefs where nutrient levels are lower^22,59,76^. This metabolic shift, known as the ‘yield to power’ switch^22^, enables copiotrophic and opportunistic microbes to outcompete others by rapidly exploiting available resources^22,59^. Further, the ocean is globally depleted of vitamin B12 (cobalamin) despite it being a major cofactor required by most marine microbes^77,78^. Interestingly, many of the 114 pMAGs prevalent on fished reefs showed functional potential for cobalamin biosynthesis both via aerobic (with 93.9% and 52.6% of pMAGs possessing cob genes within KEGG modules M00122 and M00925, respectively) and anaerobic (cbi genes: M00924; 43.9%) pathways (**Fig. 5**: Metabolism of cofactors and vitamins). In contrast, these pathways were far less common in the 236 NTMR indicator pMAGs (e.g., M00122, 40.7%; M00924, 7.20%; and M00925, 7.63%), further reinforcing the functional autonomy, environmental adaptability, and potential of fished-reef microbial indicators to outcompete other microbes.

A shift towards anabolic metabolism was proposed as a key mechanism driving ‘microbialization’ on degraded reefs, linked with fishing pressures, macroalgae overgrowth and changed reef water chemistry that select for microbial biomass accumulation and abundance shifts towards higher copiotrophic and potentially pathogenic seawater microbes^17,22,59,79,80^. This enrichment of seawater microbial copiotrophs is not limited to chronic macroalgal dominance, with recent evidence showing “microbialization” also occurs following other severe disturbances like mass bleaching episodes, where thermally stress bleached corals release labile dissolved organic matter (DOM) into the water column, significantly increasing bacterioplankton growth and abundances of copiotrophic and putatively pathogenic microbes^81^. Critically, fishing pressure in the offshore GBR is comparatively low due to its remoteness^37,41^. Our detection of microbial shifts consistent with DDAM model predictions is novel, demonstrating that microbialization is a more universal ecological response than previously recognised, detectable even under mild anthropogenic disturbance and not only on severely degraded reefs. From a management perspective, this suggests that even pristine offshore reef systems may show early microbial signs of disturbance (due to fishing), making bacterioplankton communities a sensitive indicator for reef monitoring. To test if there is greater microbial biomass in these fished reefs, future studies should collect microbial count data using readily obtainable field methods such as flow cytometry^27^.

This picture of oligotrophic taxa in NTMRs and opportunistic microbes in fished reefs also suggests a restructuring of nutrient cycling and energy flow between reef zones. For example, despite positive correlations with dissolved nitrogen (**Fig. 3A-B**), microbial indicators of fished reefs lacked canonical transporter genes for nitrate (NRT1 and NRT2 family transporters), nitrite (NrtA/NrtB systems, NirC, and the ABC transporters NrtC/D), and ammonium (including Rh family proteins SLC42A, and Amt/MEP family proteins) (**Fig. S41**). Thus, microbial indicators of fished reefs may be associated with higher NOx (**Fig. 3B; Fig. S26**) not due to direct uptake, but because they drive remineralisation of (particulate) organic matter to produce NOx (as well as ammonium; **Fig. S43**), consistent with the higher DIN:DIP ratio in fished reefs (9.9 ± 7.9) compared to NTMRs (7.2 ± 5.5; **Fig. S25**). Further research is warranted to confirm this hypothesis of enhanced microbial decomposition of POM into dissolved nitrogen in fished reefs, in addition to understanding how these taxonomic (**Fig. 2**) and functional (**Fig. 5**) reef bacterioplankton shifts between NTMRs and fished zones cascade through reef productivity, carbon cycling networks, and overall reef health.

To provide an independent, assembly-free validation of these functional patterns, we performed a community-wide read-based analysis using 4,287 GO terms derived from the same metagenomic reads (**Fig. S42**) previously published within Terzin et al. (2025)^71^, which independently corroborated and extended the functional patterns identified with indicator pMAGs (**Fig. 5**). Specifically for NTMR reefs, enrichment of MglA-type sugar ABC transporters (IPR015862) and Energy-coupling factor (ECF) transporters (IPR024919) indicates that NTMR-enriched microbes rely on ultra-high-affinity uptake systems to scavenge dissolved monosaccharides^82^ and vitamins^83^ (including B12, B1, folate, and riboflavin) directly from nutrient-depleted seawater^84^. This is consistent with the known physiology of the streamlined microbes including Pelagibacterales and SAR86 (enriched in NTMRs in MAG-based - **Fig. 2;** also read-based analysis - **Fig. S22**), and represents a hallmark of oligotrophic microbial lifestyles^82^. This is directly supported by our MAG-based analysis, which showed that NTMR indicator pMAGs had lower completeness in cobalamin biosynthesis pathways than fished reef indicators (**Fig. 5; Fig. S38**), implying NTMR-enriched microbial oligotrophs are vitamin auxotrophs that depend on environmental scavenging/uptake rather than vitamin synthesis. In contrast, fished reef communities were enriched for functions reflecting metabolic autonomy and copiotrophic resource exploitation (similar to MAG-based pathway completeness findings; see **Fig. 5**), including: active energy harvest from organic substrates (GO:0006091, GO:0016491), carbon storage via PHB biosynthesis^85^ (IPR011283), nitrogen assimilation (for anabolic reactions) through the GS/GOGAT pathway^86^ (IPR027283), *de novo* cofactor biosynthesis including folate^87^ (IPR002678) — the functional mirror image of the vitamin scavenging strategy seen in NTMRs — and flagellar motility (IPR005503, IPR007412; see **Fig. S42**), potentially used for navigation toward nutrient hotspots^88^. Together, these patterns paint a coherent picture in which NTMR microbial communities are functionally streamlined for life in oligotrophic conditions and scavenging scarce resources at high affinity, while fished reef communities are primed for rapid exploitation of elevated organic matter and nutrients through autonomous biosynthesis and active foraging.

### Seawater microbes have the potential to predict continuous environmental variables

Having established seawater microbes as reliable predictors of reef zoning status (Fig. 2), we next evaluated their potential to predict continuous environmental variables structuring microbial populations, offering a scalable, biologically integrated approach to track reef conditions beyond categorical (e.g. NTMRs vs fished) classifications. Combining random forest models^89^ and microbial niche modeling^90^, we found that seawater microbes predict several environmental variables with high accuracy (R^2^ > 0.6; *see methods*) including: seawater temperature, salinity, particulate (>0.7 µm in size) nutrients, and dissolved inorganic phosphorus (**Fig. 6A**, green boxplots). In contrast, prediction accuracies were low (R^2^ < 0.3) to moderate (0.3 < R^2^ < 0.6) for dissolved nitrogen and silicate, benthic cover variables, fish biomass, and all fish groups (**Fig. 6A**, orange and red boxplots) apart from corallivore fish, for which prediction accuracy was high (median R^2^ = 0.72; **Fig. 6A**). This ranked assessment of microbial predictive utility (**Fig. 6A**) provides a roadmap for integrating seawater microbes into reef monitoring, with future validation across other reef systems essential to confirm their broader applicability.

**Figure 6.**
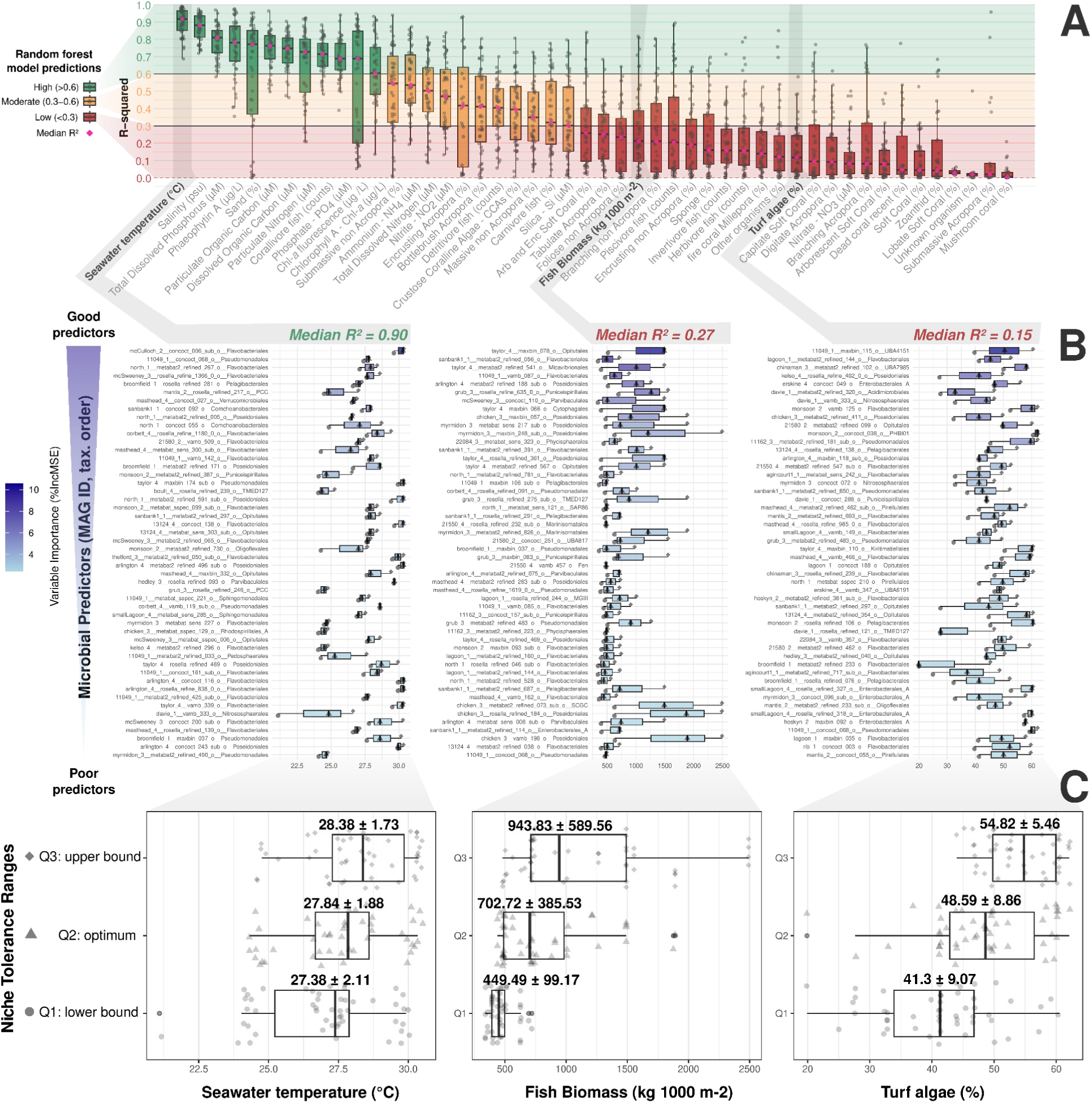
Microbial predictors of reef environmental variables, derived from random forest (RF) modeling and microbial niche analysis. (**A**) Boxplots show random forest model performance across environmental variables, based on R-squared (R^2^) values from 50 stratified permutation tests per variable. Cross-validation used an 80/20 train/test data split stratified by the GBR sector. Gray points represent individual permutations, and pink diamonds indicate median R^2^. Variables are ordered by decreasing median R^2^, with boxplots colored by performance category. The dashed line marks null performance (R^2^ = 0). (**B-C**) Boxplots show niche tolerance ranges (Q1: lower bound, Q2: optimum, Q3: upper bound) for the top 50 microbial predictors—( **B**) per pMAG and (**C**) combined across all 50 pMAGs—for three environmental variables: seawater temperature (left), fish biomass (middle), and turf algae cover (right). Niche bound values are visualised using distinct point shapes. In ( **B**), microbial predictors are additionally colored by random forest importance (%IncMSE).

Random forest predictions were validated by distinguishing microbial specialists (taxa with narrow environmental tolerance ranges that drove high prediction accuracy) from generalists, which exhibited broader niches and were thus associated with weaker predictive power. The top 50 microbial predictors for temperature (**Fig. 6B**, left), which showed a narrow thermal niche range (Q1–Q3: 27.38 ± 2.11°C to 28.38 ± 1.73°C) with an environmental optimum (Q2) at 27.84 ± 1.88°C ( **Fig. 6C**, left), were mostly Flavobacteriales (46%, primarily Flavobacteriaceae and Schleiferiaceae; **Fig. S44**). Flavobacteriaceae have previously been identified as a predictor of seawater temperature in inshore GBR reefs^91^, and temperature is well documented to be the main environmental driver structuring reef-associated bacterioplankton assemblages in the GBR^71,80^, a patterns that extends to other Pacific reef systems^92^ and to open-ocean microbial communities globally^93,94^. Accurate prediction of particulate organic carbon (median R^2^ = 0.74; **Fig. S45A**) and nitrogen (median R^2^ = 0.66; see **Fig. S45B**) is likely due to the direct role of seawater microbes (picoplankton) in producing particulate organic matter (POM) in the offshore GBR. Seasonal increases in POM during the austral summer^71^ (2.3–3.4× higher; **Fig. S30**) are associated with elevated microbial biomass, particularly cyanobacteria that contribute directly to higher POM levels^61,71,95,96^. This likely explains why summer-enriched microbes were strong predictors of summer-elevated POC (∼16% Flavobacteriales, including Arcticimaribacter, MED.G14, Croceivirga, and UBA10364) and particulate nitrogen (∼22% Flavobacteriales, ∼12% Enterobacterales, and ∼6% Pseudomonadales) (**Fig. S45**).

Unlike particulates, most dissolved nutrients and specifically nitrogen (NH_4_^+^, NO_2_^-^, NO_3_^-^, TDN) were poorly predicted by microbial communities (**Fig. 6A**). This reflects the nitrogen limitation common on the GBR outer shelf^97^, as evidenced by a DIN:DIP ratio of 8.55, which is well below the balanced Redfield ratio of 16:1^60^. Labile DIN is thus rapidly taken up by bacterioplankton, leaving nitrogen concentrations transient and unreliable as indicators of water quality on the offshore GBR reefs^96^. In contrast, phosphorus is rarely limiting and accumulates in winter (**Fig. S30**) due to reduced phytoplankton uptake^71,96^. Seasonal phosphorus dynamics in the GBR may explain the more accurate predictions of PO_4_^3-^ (R^2^ = 0.69; **Figs. 6A; S46A**) and total dissolved phosphorus (TDP; R^2^ = 0.79; **Figs. 6A; S46B**) both by winter-dominated taxa (predicting high dissolved phosphorus, e.g. *Prochlorococcus*, predominantly observed at the highest measured TDP concentrations of 0.26-0.28 µM; **Fig. S46B**) and summer-enriched taxa (e.g. Flavobacteriales, predicting lower PO_4_^3-^ and TDP concentrations; **Fig. S46**).

Spatial decoupling and niche differentiation potentially explains why benthic cover variables were difficult to predict from seawater microbial communities. While seawater microbes reflect broader, reef-scale conditions^27^, benthic cover (e.g., corals, algae) is often patchily distributed and shaped by microhabitats, contributing to spatial disconnect. Additionally, surface seawater microbial communities occupy a distinct ecological niche from those in the benthic boundary layer, where water chemistry and particle fluxes differ markedly^92,98,99^. As a result, interactions between benthic communities and seawater microbes are frequently mediated by dissolved organic matter or particles, which may dilute direct correlations and obscure mechanistic linkages. This disconnect between pelagic microbial signals and benthic cover highlights the importance of incorporating water chemistry data (**Fig. 3**) to contextualize microbial patterns, particularly in relation to nutrient and organic matter fluxes. It also points to future opportunities to improve benthic predictions by integrating spatially resolved ‘omics approaches (such as metatranscriptomics or single-cell sequencing) with benthic-proximal microbial sampling and process-based chemistry flux measurements. Explicitly incorporating hydrodynamic data including current movement, water residence time, and upwelling dynamics, would further resolve how water movement modulates the dispersal of seawater microbes, the flux of benthic-derived nutrients, and the spatial scaling of microbial-benthic coupling, ultimately refining our ability to predict reef-scale ecosystem states from microbial signatures^100^. Such an integrative approach could better capture microbial sensitivity to localised benthic inputs (e.g., coral and algal exudates) and ultimately support the development of more context-aware molecular monitoring strategies.

## Conclusions

We propose a possible mechanistic explanation for the effects of reef zoning on seawater microbial communities, proposing that zoning in the offshore GBR may shape seawater microbial communities through ecological feedbacks tied to nutrient dynamics (summarised in **Fig. 7**). In a subset of NTMR reefs, lower nutrient concentrations may be indicative of efficient nutrient cycling cumulatively driven by high fish biomass, increased algae removal due to herbivory, and enhanced coral cover, which selects for oligotrophic (microbial streamlining theory) and cooperative (Black Queen Hypothesis) microbes (**Fig. 7**, NTMRs). In contrast, fished reefs with higher turf-to-coral ratios tend to exhibit more nutrient-rich conditions, favoring metabolically independent and competitive microbes with larger genomes (**Fig. 7**, fished reefs). These changes in water chemistry and benthic cover, and their cumulative flow-on effects on microbial community composition, enabled reef zoning to be predicted from seawater microbial data alone with 71% accuracy in the offshore GBR. This high classification accuracy highlights the value of seawater microbial markers as complementary indicators of zoning effectiveness, which is relevant because zoning non-compliance is an ongoing challenge, with 500–600 annual zoning offenses (e.g., illegal fishing) potentially undermining NTMR benefits like fish spillover and biodiversity protection^37,101,102^. Considering the global success of NTMRs, future research should explore whether similar microbial shifts occur in marine reserves of other reef systems beyond the GBR. Microbial monitoring could complement existing in-water assessments by expanding spatial coverage across a broader range of reef locations and environmental conditions, offering a scalable and sensitive addition to current monitoring frameworks, ultimately informing more holistic conservation and management strategies.

**Figure 7.**
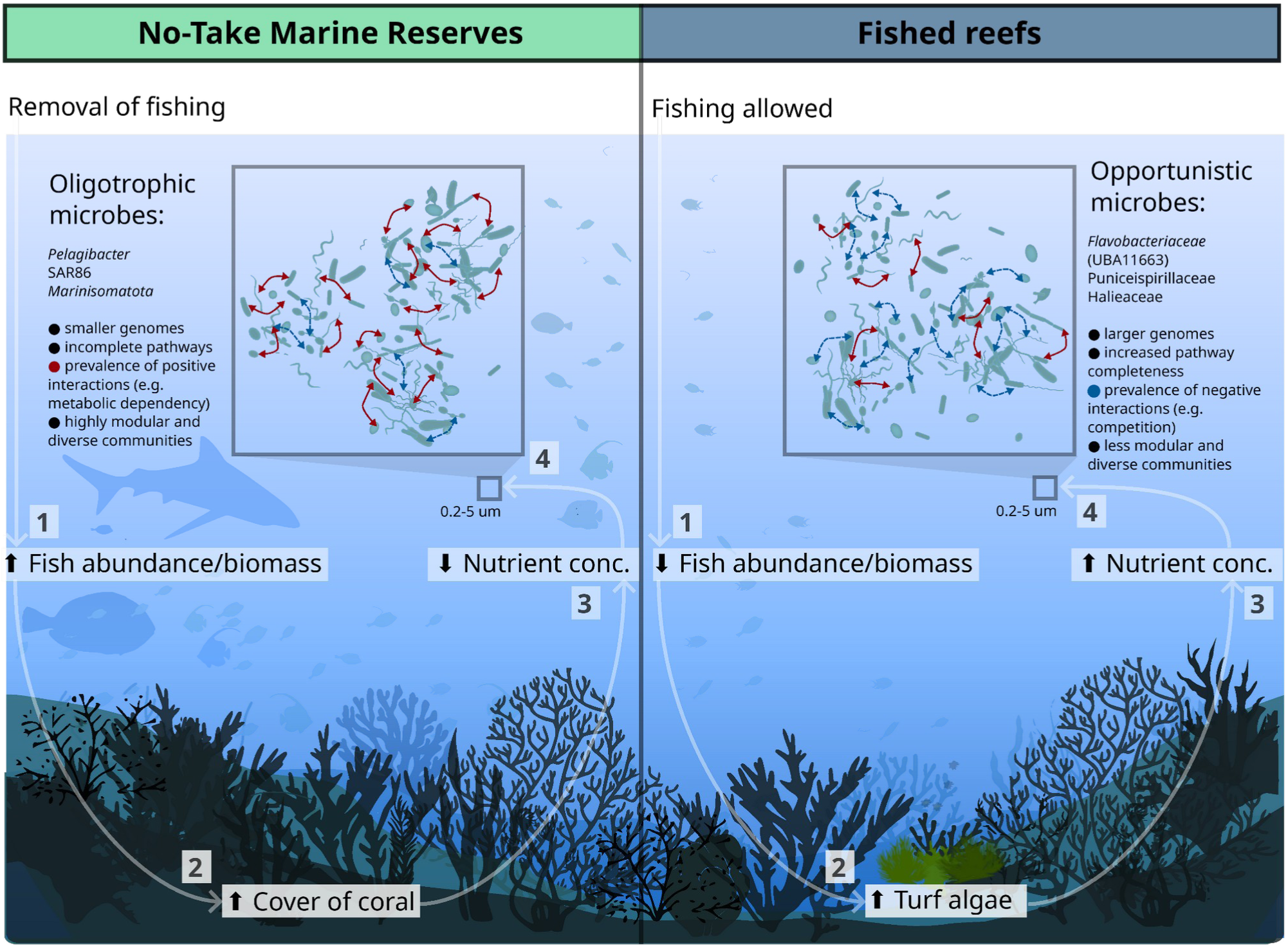
Conceptual model of the seawater microbial dynamics within No-Take Marine Reserves (NTMRs) and fished reefs in the Great Barrier Reef, in the context of physico-chemical, fish abundance, and benthic cover variables. (A) In some NTMR reefs, the removal of fishing pressure results in higher fish biomass, including herbivorous fish (1), which enhances grazing pressure on algae and promotes increased hard coral cover (e.g., *Acropora* spp., *Millepora*) and crustose coralline algae (CCA) abundance (2). Efficient nutrient cycling in such healthier reefs with elevated fish biomass and coral cover leads to lower nutrient availability (3). This reduction in nutrients imposes strong selective pressures on seawater microbial communities (4), enriching NTMRs with oligotrophic taxa (e.g., *Pelagibacterales*, SAR86, *Marinisomatota*) that exhibit streamlined genomes and incomplete biochemical pathways, as predicted by the microbial streamlining theory. These microbes form positively connected communities, likely reflecting metabolically co-dependent relationships where streamlined taxa rely on other bacterioplankton members to exchange missing metabolites, consistent with the Black Queen Hypothesis. (B) In fished reefs, lower fish biomass and reduced grazing pressure (1) result in higher turf-to-coral ratios (2) and higher nutrient concentrations (3), likely due to inefficient nutrient cycling and DOM release from turf algae, as predicted by the DDAM model. These conditions favor opportunistic microbial taxa (4), ultimately leading to distinct microbial community dynamics compared to NTMRs.

## Materials and Methods

### Seawater sampling

Water sampling was conducted on four vessel-based field trips by the Australian Institute of Marine Science (AIMS) between November 2019 and July 2020 for (1) microbial community profiling and (2) analysis of physico-chemical variables. The field trips covered the Northern (Trip 1: Princess Charlotte Bay and Cape Grenville sectors; November–December 2019), Southern (Trip 2: Swains and Capricorn Bunker sectors; January 2020), and Central Great Barrier Reef (Trip 3: Cairns and Innisfail sectors; February 2020; and Trip 4: Townsville sector; July 2020). The first three trips were conducted concurrently with Australian Institute of Marine Science Long Term Monitoring Program (AIMS-LTMP) *in situ* reef health surveys collecting data on (3) fish abundance and biomass, and reef benthic cover, with the fourth water sampling trip occurring 2 months after the AIMS-LTMP reef health surveys, at the same sites. At each reef site, surface seawater was collected at a target depth of approximately five metres (observed range: 2–10 m, reflecting natural variation in reef slope topography and tidal state across sites) using SCUBA (Trips 1–3) or Niskin bottles (Trip 4) (**Fig. 1**). Variation in sampling depth reflects differences in reef shelf proximity to the sea surface across sites, with collections standardised to a consistent clearance above the reef structure rather than a fixed vertical position in the water column, and SCUBA collections were additionally constrained within this range by standard non-ADAS dive protocols. Within this shallow depth range, the offshore GBR water column on exposed reef slopes is generally well-mixed, with minimal stratification in temperature, dissolved oxygen, or light penetration expected between 2 and 10 m.

During each trip, sampling sites were evenly distributed across fished reefs and NTMRs for a total of 48 offshore reefs (23 NTMRs and 25 fished reefs). Of the fished reefs, 20 were located in Habitat Protection zones (dark blue) where most fishing activities are allowed with the exception of trawling, while five were in Conservation Parks zones (yellow), where fishing for sea cucumbers, lobsters, and net fishing are also prohibited^37^.

### Metagenomics (field, laboratory, and data analysis) workflow for profiling seawater microbial communities

For microbial sampling, four 5 L seawater replicates from each reef were first filtered through 5 µm (Minisart® NML syringe pre-filter [Sartorius, Goettingen, Germany]) followed by 0.22 µm filters (Millipore® Sterivex-GP™ Pressure Filter [Merck Millipore, Darmstadt, Germany]) immediately after collection, and the 0.22 µm filters were snap frozen and stored at −75 °C. DNA was extracted from 0.22-µm Sterivex filters using a phenol:chloroform:isoamyl alcohol extraction with ethanol precipitation as in Botté et al (2019)^103^, with the addition of 18 µL (100 mg mL^−1^) lysozyme to the lysis buffer. Extracted DNA was quality assessed using a NanoDrop 2000 spectrophotometer (Thermo Fisher Scientific, Australia) and quantifed using a Qubit 3 fluorometer (Thermo Fisher Scientific, Australia) before submission for short-read metagenome sequencing (Nextera FLEX 2×150 bp library prep, Illumina NovaSeq) to the Australian Centre for Ecogenomics (ACE, University of Queensland, Australia) and Microba Life Sciences Ltd (Brisbane, Australia). Hybrid metagenome assemblies were generated from 27 samples (one replicate from 27 of the 48 sites) that underwent both long-read sequencing on the Oxford Nanopore PromethION platform and deep (i.e., 40 Gbp) Illumina sequencing for polishing, also at the ACE sequencing centre. High–molecular weight DNA was size-selected with the Circulomics SRE XS kit (PacBio), barcoded with the Native Barcoding Expansion kit (Oxford Nanopore Technologies), and sequenced on PromethION R9.4 flow cells using MinKNOW (v20.06.18) with default settings, with raw reads subsequently re-basecalled using Guppy (v5.0.16; Oxford Nanopore Technologies, RRID:SCR_023196) in superaccuracy mode. Adapter and barcode trimming was performed with Porechop (https://github.com/rrwick/Porechop) under default parameters.

The prokaryotic metagenome-assembled genomes (pMAGs) analysed in this study were generated as part of the Great Barrier Reef Microbial Genomics Database (GBR-MGD), using both hybrid Illumina–Nanopore and short-read-only assemblies, with hybrid long-read approaches markedly improving pMAG recovery, particularly for streamlined taxa with low GC content (<40%) and high nucleotide diversity (>0.02), and with full methodological details and code described in Robbins et al. (2025)^33^. Briefly, seawater metagenomes from 48 reefs across the GBR were assembled from the Illumina and Nanopore (27 sites) or Illumina-only data (21 sites), and pMAGs subsequently binned from the metagenome assemblies using the Aviary (https://github.com/rhysnewell/aviary; v0.3.3) assembly and genome binning pipelines. Aviary implements a hybrid metagenome assembly approach where Nanopore data are first assembled (metaFlye^104^) and polished (racon^105^, pilon^106^). Illumina data are then mapped to high-coverage Nanopore contigs to identify non-mapping reads (i.e. not represented in Nanopore assembly), and finally the non-mapping Illumina data are assembled using metaSPAdes^107^ and collated with the high-coverage Nanopore contigs to produce the final assembly. For Illumina only sites, Aviary directly assembles the Illumina data with metaSPAdes^107^. Aviary then bins pMAGs from each reef using several binning tools (MetaBAT1^108^, MetaBAT2^109^, MaxBin2^110,111^, CONCOCT^112^, Vamb^113^, and Rosella https://github.com/rhysnewell/rosella) and collates a representative set of bins with DAS Tool^114^. pMAGs were retained if they met a quality threshold of ≥50 (calculated as completeness – 3×contamination) as assessed using either CheckM1^115^ or CheckM2^116^.

The resulting 5,283 high-quality pMAGs (**Supplementary Data 1;** deposited under EBI BioProject PRJEB82623) were taxonomically classified with the Genome Taxonomy Database Toolkit (GTDB-Tk^117^, release R214) and subsequently dereplicated at 95% ANI using CoverM^118^ (v0.6), resulting in a total of 876 “species-resolved” pMAGs_95%ANI_ (**Supplementary Data 2**) used in downstream analysis (and hereinafter referred to as pMAGs for brevity). Read mapping counts and relative abundances were inferred by mapping Illumina short reads (≥95% identity, ≥75% aligned length) to the dereplicated set of pMAGs using minimap2^119^ (as implemented in CoverM). To account for sparsity and compositionality issues inherent to microbial metagenomics data, raw read counts were transformed using a center log-ratio (CLR) transformation in the microbiome^120^ (v1.24.0) R package. All statistical analysis and visualization were performed on CLR-transformed and relative abundance data, and final figures were compiled using Inkscape^121^ (v0.92.5).

The de-replicated set of 876 pMAGs were imported into anvi’o^122^ (v8) for functional annotation. Using the anvi’o *contigs* workflow (https://anvio.org/help/main/workflows/contigs/), open reading frames (ORFs) in each pMAG were first predicted using Prodigal^123^ (v2.6.3) and then compared against anvio’s set of protein hidden markov models (HMMs) for functional annotation with the *anvi-run-hmms* command. Using the *anvi-run-kegg-kofams* program, ORFs that matched to HMMs were then mapped against the Kyoto Encyclopedia of Genes and Genomes (KEGG) Orthology (KO) database^124^ using HMMER (v3.3.2, http://hmmer.org/). These mappings were used to assess the stepwise completeness of metabolic pathways in each pMAG with the *anvi-estimate-metabolism* program using KEGG ‘modules’. The resulting KO annotations and KEGG module completeness statistics were exported from anvi’o (with *anvi-export-functions*) for downstream indicator analyses of NTMRs and fished reefs (**Supplementary Data 3**). From 358 detected KEGG modules identified in our data (**Supplementary Data 3**), Sparse Partial Least Squares Discriminant Analysis (sPLS-DA)^34,35^ was used to identify the top 50 modules showing differential completeness (0 - KEGG module absent; 1 - complete module) between NTMR-associated (n=236) and fished reef-associated (n=114) microbial indicators, which was further validated with Wilcoxon rank sum tests in R.

### Seawater processing for physico-chemical variables

Water chemistry was measured in three of the four seawater replicates using established methods, following the standard operating procedures of the AIMS Water Quality Monitoring Program^125^, while the fourth replicate was reserved exclusively for microbial community profiling. Fourteen variables were measured, including dissolved nutrients (total dissolved nitrogen – TDN; ammonium – NH_4_^+^; nitrite – NO_2_^−^; nitrate – NO_3_^−^; total dissolved phosphorus – TDP; phosphate – PO_4_^3−^; dissolved organic carbon – DOC; silicate – Si), particulate fractions (particulate organic carbon – POC; particulate nitrogen – PN; particulate phosphorus – PP), pigments (chlorophyll a – Chl-a; phaeophytin a – Phaeo), and total suspended solids – TSS. Immediately following collection, dissolved nutrient samples (NH_4_^+^, NO_2_^-^, NO_3_^-^, PO_4_^3-^, TDN, TDP, DOC, and Si) were filtered (0.45 **μ**m cellulose acetate Sartorius Minisart NML) into 10 mL acid-washed vials (triple pre-rinsed with filtered site water) and all analytes except DOC and Si were stored frozen (−18 °C) until laboratory processing. Samples for DOC and Si analyses were stored refrigerated (4 °C) until analysis, with DOC having been acidified with 100 μL AR-grade hydrochloric acid immediately following sampling. Particulate variables (POC, PN, PP, and Chl-*a*) were filtered onto pre-combusted (450 °C for 4 h) 25 mm glass fibre 0.7 µm filters (Whatman GF/F), wrapped in foil, and stored frozen (−18 °C) until analysis. TSS samples were filtered onto pre-weighed 47 mm polycarbonate filters with 0.4 **μ**m pore size (GE Water & Process Technologies), triple-rinsed with ultrapure water, and stored refrigerated until analysis.

Laboratory analyses were conducted at the AIMS Analytical Technology and Water Quality Laboratories within one month (Chl-*a*, DOC, TSS) or three months (all other variables) of collection. Dissolved nutrient concentrations (NH_4_^+^, NO_2_^-^, NO_3_^-^, PO_4_^3-^, Si, TDN, and TDP) were determined using wet chemical methods^126–128^ on a segmented flow analyser (Seal AA3 and AA500). Concentrations of NH_4_^+^ (in the forms of both NH_3_ and NH_4_^+^) were analysed by fluorescence spectroscopy (o-phthalaldehyde or OPA method)^129^. Concentrations of PO_4_^3-^, NO_2_^-^, NO_3_^-^+NO_2_^-^, and Si were analysed by colourimetry. For both PO_4_^3-^ and Si, colour formation is based on the molybdenum blue reaction; for NO_2_^-^ and NO_3_^-^+NO_2_^-^, colour formation is based on the Griess reaction, with the use of a cadmium reduction column. Nitrate (NO_3_^-^) is then calculated as the difference between the concentrations of NO_2_^-^ and NO_3_^-^+NO_2_^-^. Alkaline persulfate digestion^130^ was applied to TDN and TDP samples prior to analysis following methods for NO_3_^-^+NO_2_^-^ and PO_4_^3-^ as described above. Organic carbon (DOC and POC) and PN were analysed via high temperature catalytic combustion (Shimadzu TOC-L) with solid sample module (SSM-5000A) and nitrogen module (TNM-L). Concentrations of PP were determined via hot acid persulfate digestion^131^ to release free phosphate, followed by colourimetric analysis (antimony-phosphomolybdate reaction) and analysis on a spectrophotometer (Shimadzu UV-1900i)^126^. Chl-*a* filters were ground and incubated for 2 h in 90% acetone prior to reading on a fluorometer (Turner 10AU). Samples were then acidified and re-read to determine the concentration of Phaeo^132^. TSS samples were dried (60 °C overnight) immediately upon returning to the laboratory and post-weighed for gravimetric analysis on a 6 decimal place balance (A&D BM-20 micro analytical balance). AIMS uses traceable pure chemicals for its nutrient calibrations and performs quality checks with each analytical run. Reference materials include in-house quality controls and certified reference materials (KANSO Reference Material for Nutrients in Seawater), and the laboratory participates in bi-annual Quasimeme proficiency testing.

In addition, underway physico-chemical data (temperature, salinity, Chl-*a* fluorescence) were obtained from IMOS Ships of Opportunity sensors (SBE 38 thermometer, SBE 21 Thermosalinograph, WET Labs ECO-FLNTU-RT fluorometer) at 1.9 m (RV Cape Ferguson) or 2.5 m (RV Solander), recording the closest measurement to sampling time^133,134^. This resulted in 17 physico-chemical variables analysed in this study: 14 water chemistry parameters, temperature, salinity, and Chl-*a* fluorescence. A more detailed protocol can be found in Terzin et al. (2025)^71^.

### Fish abundance and biomass, and reef benthic cover

Standardised protocols were used to survey benthic and fish assemblages at 48 reefs as part of the Long-Term Monitoring Program (LTMP) by AIMS, further described below. Trained divers conducted fish and benthic surveys on SCUBA at three sites per reef in a standard reef slope habitat, simultaneously with collections of seawater samples for metagenomic sequencing and physico-chemical analyses. At each site, five 50 m permanently marked transects were surveyed, set parallel to the reef crest, and at depths between 4 and 12 m (depending on site bathymetry).

Along each transect, large-bodied, mobile fishes were surveyed within a 5 m belt (250 m^2^ area) to capture diurnal, non-cryptic reef fish species. Non-cryptic, diurnal small-bodied fishes (e.g., Pomacentridae, small Labridae) were also surveyed using a transect width of 1 m (50 m^2^ area). Fish were identified to species and abundance counts were subsequently aggregated into trophic groups and at family level based on the latest available classification. For each individual fish, length was estimated using predefined size classes (5 cm bins for large mobile fishes counted on the 5 m belt, 2 cm bins for species counted on the 1 m belt). Observer consistency was maintained through annual cross-calibration trips, following the procedures described in Emslie et al. (2018)^135^.

Fish biomass was calculated from length estimates using species-specific length-weight relationships, and based on the formula: Biomass = Abundance × *a* × (Midpoint)^b^, where Abundance is the number of fish, and *a* and *b*are species-specific coefficients derived from FishBase^136^. The midpoint refers to the average length of fish within each size category. Total biomass (kg per 1000 m^2^) for each transect was summed to estimate the overall fish biomass for the surveyed area (i.e., per reef).

Benthic surveys were conducted concurrently along the same transects using digital imagery. Digital images of the substrate were taken on the up-slope side of each transect at 50 cm intervals. Estimates of proportional benthic cover were subsequently derived from the identification of the benthos beneath five fixed points arranged in a quincunx patterns digitally overlaid onto these images. A total of 40 images from each transect (n = 3000 points reef^-1^) were randomly selected and analysed using the Image Classifier software, with analyses for this study conducted prior to transition to the Reef Cloud platform (https://reefcloud.ai). Hard corals were identified to the lowest taxonomic resolution possible, usually genera^137^. A total of 37 *in situ* variables were used, encompassing benthic cover (abiotic substrates, soft corals, hard corals by morphology, algae, sponges, and other biota), and fish community data (aggregated per trophic group, family-level taxonomy, and overall biomass); a complete list is provided in Supplementary Data 4.

### Reef zoning influences on seawater microbes: community-based and feature selection approaches

Principal Components Analysis (PCA) in mixOmics^35^ (v6.26.0) (R version 4.3.2; R Core Team, 2023) was first used to explore major sources of variation in microbial community composition (Figs S1-2). To test whether reef zoning explains significant community-level differences while accounting for spatiotemporal structure, two complementary approaches were applied on Aitchison distances calculated from centred log-ratio (CLR)-transformed abundance data: PERmutational Multivariate ANalysis Of VAriance (PERMANOVA), using the *adonis2()* function, and Distance-Based ReDundancy Analysis (dbRDA) using the *dbrda()* function in vegan^71,72^ (v2.6-4). Both models included reef protection status as the predictor of interest, with sampling trip, geographic sector, and reef identity as covariates to account for spatiotemporal confounding (model formula: aitchison_dist ∼ Open_or_Closed_to_fishing + Sampling_trip + SECTOR_N_S + REEF_NAME), using 9,999 permutations for significance testing. The significant community-level signal of reef zoning provided the statistical foundation for subsequent supervised analyses to identify specific indicator taxa (Figs. S3-4; Table S1), and microbial community data were partitioned by GBR sector since AIMS-LTMP benthic cover and fish abundance data have historically been analysed across GBR sectors^7,138^.

To identify microbial indicators that consistently discriminated NTMRs from fished reefs across GBR sectors, we used Multivariate INTegration Sparse Partial Least Squares Discriminant Analysis (MINT sPLS-DA)^34–36^. For comparison, a conventional (i.e., without sector integration) sPLS-DA was also performed to identify microbial indicators of reef zoning, which yielded discriminatory features but retained strong spatiotemporal batch effects (**Figs. S5–S9; Table S2**), supporting the use of MINT for this dataset.

MINT sPLS-DA was run on microbial abundance data (CLR-transformed), with model tuning performed via Leave-One-Group-Out Cross-Validation in which models were iteratively trained on six sectors and validated on the remaining sector. The optimal number of components and number of features to retain per component were selected based on cross-validation performance, and the final MINT sPLS-DA model was fitted using the selected parameters (**Fig. S10–S11; Tables S3–S5**). Results were visualised in mixOmics^35^ and ggplot2^139^ with sample-level ordination plots (to visualise classification structure) and heatmaps to show abundances of discriminatory microbes across sites, and the reproducibility of indicator microbes across sectors.

To validate the statistical significance of the zoning signal, we implemented zone-label shuffling to generate a null distribution against which observed MINT sPLS-DA performance could be benchmarked. Reef protection status labels were randomly permuted 999 times within each sector (preserving spatial structure) to break any association between microbial community composition and zoning status, after which the MINT sPLS-DA model was re-run with identical parameters (as in **Fig. S10–S13; Tables S3–S5**) and classification accuracy was evaluated via LOGOCV. The observed classification accuracy was compared against the resulting null distribution using a permutation test, and effect size was quantified using Cohen’s *d*(**Fig. S14; Table S6**).

Other approaches, such as Random Forest (RF) was also compared with MINT following the same leave-one-sector-out cross-validation and permutation testing procedures (999 permutations with zone-label shuffling within sectors). RF showed poorer classification performance given the inability of the method to handle the variability between independent sector sampling. However, a small core microbial signal confirmed our MINT results (Supplementary Methods; Fig. S15).

Per-pMAG differential abundance testing was also conducted using ALDEx2 (ANOVA-Like Differential Expression tool v1.34.0)^140^, which accounts for compositional data structure through Dirichlet-multinomial modeling. Two ALDEx2 approaches were implemented: (1) pairwise comparisons using Welch’s t-test and Wilcoxon rank-sum tests on CLR-transformed abundances (128 Monte Carlo samples, denominator = "all") without covariates, and (2) Generalized Linear Models (GLMs) with spatiotemporal covariates (model formula: abundance ∼ Open_or_Closed_to_fishing + Sampling_trip + SECTOR_N_S). Both models used Benjamini-Hochberg false discovery rate (FDR) correction (α = 0.05) (**Figs. S16-S20**).

To further evaluate whether indicator pMAG selection could be confounded by genome size bias (wherein longer genomes naturally accumulate more mapped reads, potentially introducing spurious abundance patterns in CLR-transformed data), we performed presence/absence analysis of all 350 MINT sPLS-DA-selected indicator pMAGs across the 190 samples. Using raw count data (prior to CLR transformation), we calculated detection frequencies separately for NTMR and fished zones (Fig. S21).

To validate findings using read-based metagenomics as an assembly-independent approach^141^, raw Illumina reads were (1) mapped against the NCBI nr database using DIAMOND^142^ (v2.0.9) with an e-value threshold of <10^-5^, (2) taxonomic profiles were generated in MEGAN^143^ (v6.23.0), and (3) low-abundance prokaryotic taxa (relative abundance <0.0001%) were removed to minimize spurious signals, following the methods detailed in Terzin et al. (2025)^71^. To provide a community-wide functional perspective complementary to the KEGG module completeness analysis of indicator pMAGs [see section: *Metagenomics (field, laboratory, and data analysis) workflow for profiling seawater microbial communities*], GO term annotations were additionally assigned to each read in MEGAN and collapsed at GO rank 5. Similar to taxonomic profiling, pre-filtering steps included removal of non-annotated reads, eukaryotic and viral reads, and rare GO terms (relative abundance <0.0001%), yielding a final dataset of 4,287 GO terms, which were then were CLR-transformed (after the addition of pseudocounts)^71^. MINT sPLS-DA was rerun (as described above) to identify if similar indicator microbes and GO terms for NTMRs and fished reefs were selected across both MAG- and read-based analyses (**Fig. S22** for microbial taxa; **Fig. S42** for microbial functional features)

### Correlating microbial abundance data with 54 continuous environmental variables

To further investigate environmental factors that may be driving microbial community patterns in NTMRs and fished reefs, we applied MINT Sparse Partial Least Squares (sPLS) analysis to identify important associations between 350 seawater microbes indicating reef zoning with continuous environmental data. Unlike MINT sPLS-DA, which is used for classification with a categorical outcome, MINT sPLS^36,44,45^ integrates two continuous datasets (i.e., microbial abundances and environmental variables) while accounting for study-specific (i.e., GBR sectors) variation. First, median values were calculated per reef site for each of the 54 environmental variables (physico-chemical measurements and measures of benthic cover, fish assemblage abundances, and overall fish biomass) to account for differences in the number of replicates between microbial (n = 4) and environmental (n = 3) samples. Then, MINT sPLS was used to identify important associations between 350 indicator seawater microbes (CLR-transformed abundances) and the 25 environmental variables that exhibited the highest covariance with the microbial data, as determined by the sPLS model’s variable selection procedure^36,44,45^. MINT sPLS results were visualised in ggplot2^139^ with a heatmap (displaying microbiome-environment associations) and a biplot reintroducing the context of the samples, thus visualising how well the latent components separate groups of interest (i.e., NTMRs and fished reefs).

### Generalised Linear Mixed Models (GLMMs)

After identifying environmental variables significantly associated with microbial indicators of NTMRs versus fished reefs, we tested whether these key metrics themselves differed between protection zones using generalised linear mixed models (GLMMs). To analyse differences between NTMRs and fished areas, all models included protection status as a fixed effect (NTMR vs. fished) as well as the random terms of latitudinal sector and position across the continental shelf, with nested random effects including reef, site nested within reef, and transect nested in site. Herbivore density (abundance numbers per 1000 m^2^) was modelled against a negative binomial distribution to handle overdispersion, total fish biomass with a Gamma distribution with a log link function. Hard coral and turf algae were modelled against binomial distributions representing the number of points classified as hard coral or turf algae out of the total number of points surveyed. All models were implemented in R using the glmmTMB (v1.1.10) package^144^, with significance assessed via Wald z-tests (α = 0.05) and model diagnostics confirming appropriate fit using the DHARMa (0.4.7) R package^145^.

### Microbial species co-occurrence networks between NTMRs and fished reefs

Graph theory is increasingly used to quantify stability in microbial co-occurrence networks^72,73^. Recent frameworks^70^ suggest that healthy microbial communities show stronger partitioning (high connectivity within network modules, i.e., tightly linked microbial subgroups), which limits environmentally induced shifts within these modules. In contrast, unstable microbial communities (undergoing stress) are predicted to exhibit higher between-module connectivity which was proposed as a metric of destabilised networks, as environmental fluctuations may propagate widely across the community, potentially resulting in network collapse if critical thresholds are exceeded^70^. To assess network stability in reef bacterioplankton within NTMRs and fished reefs, we computed two complementary network metrics: cohesion (sample-level) and connectedness (microbe-level) following Herren & McMahon (2017)^69^, as well as modularity (network-level) following Hernandez et al. (2021)^70^.

Network connectedness and cohesion are defined as network metrics that quantify the degree of positive and negative connectivity within a microbial community^69^. Briefly, connectedness (microbe-specific metric) is derived by first computing pairwise Pearson correlations between taxa from relative abundance data. To correct for biases inherent in compositional data, a null model (e.g., "taxon shuffling") randomises abundances for all taxa except the focal one across 200 iterations, generating expected correlations that are subtracted from observed correlations. The resulting corrected correlations are then averaged separately for positive and negative values, yielding taxon-specific positive and negative connectedness with the broader community. Cohesion (sample-specific metric) is then calculated for each sample by weighting the relative abundance of each microbe by its connectedness values and summing these products across all microbial taxa, producing two metrics per sample (i.e., community): positive cohesion (sum of positive contributions) and negative cohesion (sum of negative contributions). In contrast to existing correlation detection methods which aim to identify significant pairwise associations (i.e., between two taxa), connectedness and cohesion therefore evaluate connectivity at the community level, and were used in our study to compare the prevalence of positive (i.e., mutualism, commensalism, co-dependency due to metabolic exchange) and negative (i.e., competition, predator/prey, pathogen/host, parasite/host, and etc.) correlations in reef bacterioplankton between NTMRs and fished reefs. It should be noted however that these correlations may also reflect indirect microbial associations such as shared responses to the same environmental drivers or shared interactions with a third microbe, and are not always a reliable proxy of direct biological interactions^69^.

For modularity, we constructed co-occurrence networks for each sector × zone combination (n = 14 networks) using FlashWeave^146^ (v0.19.2), which infers direct associations via conditional dependencies. Raw counts were CLR-transformed with pseudocounts added, and networks were built in sensitive mode (α = 0.01, max_k = 2) with a minimum of 4 samples per network. Positive partial correlations (weight > 0) were retained as edges, consistent with the predominance of positive associations in global plankton interactomes (Chaffron et al., 2021)^72^. Modularity was computed on binary (unweighted) networks using the Clauset-Newman-Moore greedy algorithm^147^ (measuring the degree of compartmentalization), as implemented in NetworkX^148^ (v3.6.1). Statistical comparisons between NTMR and fished reefs used Mann-Whitney U tests on the seven sector-specific modularity values per protection status. To further assess network compartmentalisation, we quantified, for each of the 14 sector×zone networks, the proportion of edges falling within versus between modules (using the same Clauset-Newman-Moore module assignments), and compared the within-module edge fraction and the within:between edge ratio between NTMRs and fished reefs using Mann-Whitney U tests. Results were visualised as barplots for connectedness (expressed as positive/negative edges ratio, for the 350 indicator pMAGs discriminating NTMRs and fished reefs) and as boxplots for cohesion (expressed as positive/negative cohesion ratio) and modularity. For each of the 350 indicator pMAGs, linear regression was used to find a relationship between positive/negative connectedness ratio (as a response variable) and (1) genome size, (2) GC content, and (3) potential for metabolic independence (expressed for each pMAG as an average completeness across obtained KEGG modules, similar to Veseli et al. 2024^65^) as the three predictor variables.

### Predictions of continuous environmental variables using seawater microbial data

To identify if microbial markers can predict environmental variables in the surrounding seawater, Random Forest (RF) models^89^ were trained on CLR-transformed microbial abundances to predict continuous environmental response variables (physico-chemical measurements, benthic cover, and fish abundance and biomass). RF models (500 trees; mtry = √p, where p = number of microbial features; node size = 5) were validated using stratified (strata = GBR sectors) and site-aware (all replicates from a given site were grouped within each split) 80/20 train-test splits repeated 50 times, and with fixed random seeds for reproducibility. This repeated subsampling approach provides robust performance estimates by mitigating geographic (sector and site-specific) and temporal biases, while also avoiding information leakage (i.e., training the data on a subset of site replicates and validating on left-out replicates of the same site). RF models were implemented via the randomForest (v4.7-1.2) R package^149^.

For each environmental variable, model performance was quantified using mean R^2^ values (i.e., by aggregating results across 50 permutations). While no universal R^2^ interpretation exists for marine microbiome-environment prediction studies, we defined RF model predictive performance as high (R^2^ ≥ 0.6), moderate (0.3 ≤ R^2^ < 0.6), or poor (R^2^ < 0.3) based on existing literature ^91,150,151^. Predictor importance was quantified using the percentage increase in mean squared error (%IncMSE), with higher values indicating stronger predictive contributions (as shuffling the values of that microbe significantly lowered the RF model’s performance). Microbe-specific %IncMSE scores were averaged across 50 permutations, and the top (i.e., the highest average %IncMSE) 50 microbes per variable were selected.

### Inferring microbial niches

To further validate these predictions, we computed the microbial niche tolerance ranges for consensus markers, defined as the 25th–75th percentile of environmental values beyond which that microbe is typically not observed. Microbial niches were calculated following Chaffron et al. (2021)^72^, using the robust optimum (RO) method^90^. In this approach, the ecological optimum (Q2) represents the ideal living conditions for a microbe concerning a given environmental parameter (i.e., where a specific pMAGs will be found at its highest relative abundance), whereas the lower (Q1) and upper (Q3) niche bounds correspond to the minimum and maximum values of that environmental variable beyond which the microbial taxon is rarely observed. These bounds represent the environmental ranges within which each microbial taxon can be found at varying levels of relative abundance, and for each microbe, its tolerance (niche) range for a specific environmental variable is calculated as the interquartile range (Q3–Q1). Microbial niches (i.e., Q1, Q2, and Q3 values for each of the 876 pMAGs, and for each of the 54 environmental variables) were computed three times to estimate global microbial niche across all samples, and separately for NTMRs and fished reefs.

Relative abundance data for each of the 876 microbial pMAGs was normalised in a two-step manner: by introducing pseudocounts (i.e., adding 1 to abundance values to avoid issues with zero counts) and then dividing abundances by the geometric mean, scaling the data to account for variations between microbial features and ensuring the data was standardised across samples. Based on obtained relative abundance distributions across samples, niche tolerance ranges were computed for each of the 876 pMAGs—specifically the lower bound (Q1, i.e., 25th percentile), ecological optimum (Q2, i.e., 50th percentile), and upper bound (Q3, i.e., 75th percentile)—relative to each of the 54 continuous environmental variables. Niches were computed only based on the samples where the microbial feature was present, which was done by identifying prevalent samples where the microbial feature was observed (i.e., where the abundance value of that pMAG was greater than zero), and the environmental variables associated with these prevalent samples were then extracted to compute microbial niche bounds. Finally, the resulting niche bounds (Q1, Q2, and Q3) for each microbial feature (876 pMAGs) and each of the 54 continuous environmental variables were compiled into a comprehensive dataset, providing detailed information on the optimal environmental conditions for each microbe. We visualised the niche tolerance ranges for microbial predictors identified in RF models using boxplots generated within ggplot2^139^ (v3.5.1).

## DECLARATIONS

### Ethics approval and consent to participate

Samples were collected under the permit G12/35236-1 issued by the Great Barrier Reef Marine Park Authority.

### Consent for publication

Not applicable.

### Availability of data and material

Sequencing data and primary metagenomic assemblies have been uploaded to the European Nucleotide Archive (ENA) under project accession PRJEB82623 under the project name “Great Barrier Reef seawater microbiomes genome database”. The 5,283 prokaryotic metagenome-assembled genomes (pMAGs) are accessible from Zenodo (DOI: 10.5281/zenodo.17109887).

The associated physico-chemical variables have been uploaded to the IMOS-AODN repository and are available from the Australian Institute of Marine Science (AIMS) (2022): Great Barrier Reef Microbial Genomics Database: Seawater Illumina Reads. https://doi.org/10.25845/Q4XH-YN10.

The benthic cover, fish abundance, and biomass data are managed by the AIMS Long-Term Monitoring Program (LTMP) and can be accessed via the AIMS data portal (https://apps.aims.gov.au/metadata/view/a17249ab-5316-4396-bb27-29f2d568f727).

Metagenomic analysis including metagenome hybrid assembly, binning, taxonomic annotation, and abundance estimation of MAGs is described in Robbins et al. (2025).

All additional code including indicator analysis, microbial networks, integration of microbial and environmental data, and random forest machine learning is available at: https://github.com/mterzin/fishy_microbes (Terzin, 2025). The microbial niche analysis was performed following the protocol from Chaffron et al. (2021), and the specific code for this analysis is available from the authors of that study upon request.

### Authors’ contributions

NSW obtained funding for the project. NSW, DGB, PWL, RKG, and SJR conceived the sampling design. SCB collected seawater in the field (for metagenomics and physico-chemical data) and processed all samples in the laboratory for metagenomic sequencing. LTMP data (benthic cover and fish abundance/biomass) were processed by MJE and DMC as part of the LTMP, with GLMMs run by MJE. Metagenomics analyses including hybrid assembly, binning, taxonomic annotation, and abundance estimation were performed by SJR, YKY, KED, and JZ, with the guidance of PH, PWL, and DGB. MT performed the functional annotation of metagenomes and carried out all subsequent analyses, including dataset integration and data visualisation, with assistance from PWL, KALC, YKY, DGB, SJR, SC, RKG, and MJE. MT wrote the original draft of the manuscript, and all authors made substantial contributions to its form. All authors critically reviewed the manuscript before submission.

## Acknowledgements

The seawater samples analysed in this study for metagenomics and physico-chemical variables were collected across 48 reefs, from the sea country of various Indigenous groups who are Traditional Owners (TOs) of that land. We acknowledge the Gudang Yadhaigana TOs, custodians of the McSweeney, Monsoon, 11-049, and 11-162 reefs, which lie within their sea country estate. We pay our respects to the Kuuku Ya’u TOs of the Mantis and Lagoon reefs, and the Lama Lama TOs of the 13-124, Davie, and Corbett reefs in the western half of their territory. We also recognise the Cape Melville, Howicks, and Flinders Island TOs of the eastern half of Corbett and Sand bank #1 reefs, as well as the Eastern Kuku Yalanji TOs of St Crispin and Agincourt #1. We extend our respect to the Yirrgandji TOs of Hastings reef and the Gunggandji as TOs of Arlington, Thetford, and Moore reefs. We acknowledge the Gunggandji-Mandingalbay Yidinji TOs of McCulloch, Hedley, Peart, and Feather reefs, and the Mandubarra TOs of Farquaharson Reef. We honour the Girringun Aboriginal Corporation TUMRA for their connection to Taylor Reef, and the Manbarra TOs of Rib, Kelso, Little Kelso, and John Brewer reefs. We also recognise the Wulgurukaba TOs of Myrmidon, Grub, and Helix reefs, the Bindal Traditional Owners of Knife, Fork, Centipede, Chicken, and Lynchs reefs, and the continuing connection of the Manbarra TOs to Roxburgh, and Fore and Aft reefs. Lastly, we acknowledge the PCCC TUMRA for their stewardship of North, Bloomfield, Eskine, Mast Head, Hoskyn, Fairfax, and Boult reefs. We pay our respects to their Elders, past, present, and emerging, and acknowledge their enduring connection to land and sea. Further, our desktop / lab research took place at the Australian Institute of Marine Science (AIMS) headquarters at Cape Ferguson, and we wish to acknowledge the Wulgurukaba and Bindal peoples as the Traditional Owners of that land. This research was also undertaken at the JCU Townsville Bebegu Yumba campus, and the authors acknowledge that the Australian Aboriginal and Torres Strait Islander peoples are the original inhabitants and traditional custodians of this continent and have unique cultural and spiritual relationships to the land and waters. We acknowledge the AIMS Water Quality team, especially Ulysse Bove, Keeley Glasson, and Daniel Moran for logistics, training, and processing of water chemistry samples. We acknowledge the AIMS-LTMP team and others involved in field collection and preparation of samples including Emmanuelle Bove, Johnston Davidson, Veronique Mocellin, and Josephine Nielsen. We thank the crew of the RV Solander and RV Cape Ferguson for their excellent logistical support in the field. We also acknowledge Gene Tyson for his support in facilitating the use of the NovaSeq at Microba Life Sciences Ltd. (Brisbane, QLD, Australia). We extend our gratitude to Murray Logan for his insightful discussions on the appropriate statistical handling of the data. AI tools (ChatGPT and DeepSeek) were used exclusively for proofreading (grammar, syntax, and clarity checks) and code assistance (debugging and statistical script optimization). No AI tools were used to generate original content, interpret data, or formulate conclusions. KALC was supported in part by the National Health and Medical Research Council (NHMRC) Investigator Grant (GNT2025648). MT extends his gratitude to the members of the Lê Cao Lab at Melbourne Integrative Genomics (MIG) for the supportive environment and valuable scientific discussions, particularly Dr Vinicius Salazar, Dr Saritha Kodikara, and Dr Jiadong Mao.

