## Supplementary_Material for "No-take marine reserves promote oligotrophic reef bacterioplankton communities across the Great Barrier Reef"

### Field sampling design

The coordinates of the 48 surveyed reefs were visualized in R Studio (R version 4.3.2)[1] using the R packages: raster (version 3.6.26)[2], tidyverse (version 2.0.0)[3], ggspatial (version 1.1.9)[4], sf (version 1.0.15)[5,6], dataaimsr (version 1.1.0)[7], gisaimsr (version 0.0.1) (https://github.com/open-AIMS/gisaimsr), and ggrepel (version 0.9.5)[8].

### PCA - Principal Components Analysis | What are the main clustering patterns across our samples?

Principal Components Analysis (PCA) was applied in an R package mixOmics[9] as an unsupervised approach to visualise the main clustering patterns between reef sites based on microbial community profiles. The number of optimal PCA components was determined using the tune.pca() function in mixOmics.

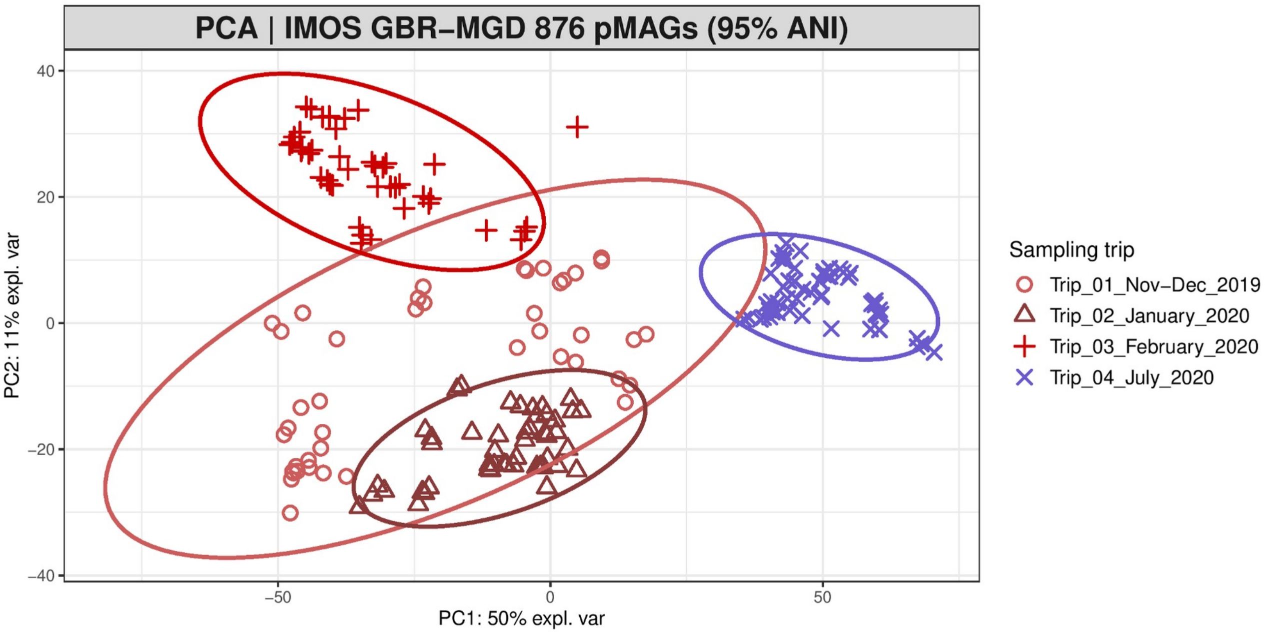


**Figure S1. Main clustering patterns of seawater microbial communities, colored per sampling transect**. The PCA ordination plots show clear differences between microbial communities sampled during the summer/wet season (red) and winter/dry season (blue), with 50% of variance being attributable to dimension 1. Samples collected in the peak of summer (Trip 3) additionally separate from early summer sampling (Trips 1 and 2) on PCA dimension 2.

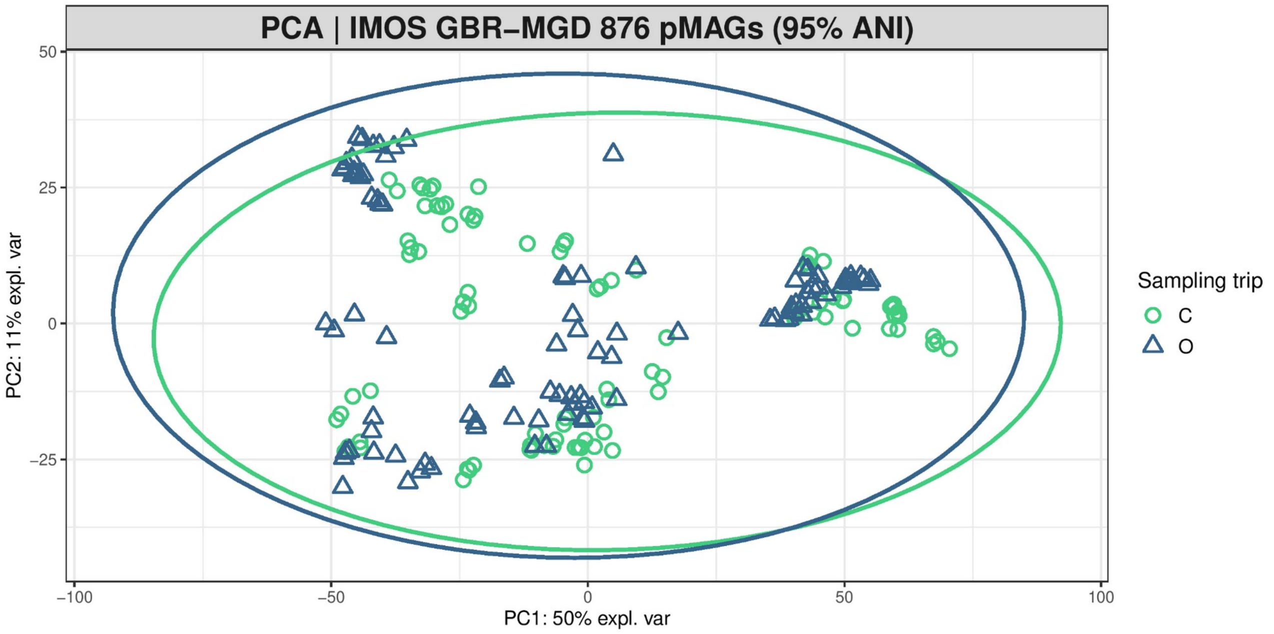


**Figure S2. Clustering patterns of seawater microbial communities based on reef zoning.** PCA ordination does not show clear clustering between No-Take Marine Reserves (C - closed to fishing, green) and fished reefs (O - open to fishing, blue).

### Community-level tests

#### **PER**mutational **M**ultivariate **AN**alysis **Of** **VA**riance (**PERMANOVA**)

To formally test whether reef zoning explains significant community-level differences after accounting for spatiotemporal structure, PERmutational Multivariate ANalysis Of VAriance (PERMANOVA; adonis2() function in vegan[10,11] v2.6-4) was used to test the statistical significance of reef protection status as a predictor of community composition. Reef protection status was included as the predictor of interest, with sampling trip, geographic sector, and reef name as covariates to account for spatiotemporal confounding (model formula: aitchison_dist ~ Open_or_Closed_to_fishing + Sampling_trip + SECTOR_N_S + REEF_NAME), using 9,999 permutations for significance testing.

When accounting for spatiotemporal factors, reef protection status remains a highly significant predictor of microbial community composition, though its explanatory power is considerably smaller than spatiotemporal factors like sampling trip, reef site, and Great Barrier Reef sectors (**Fig. S3; Table S1**).

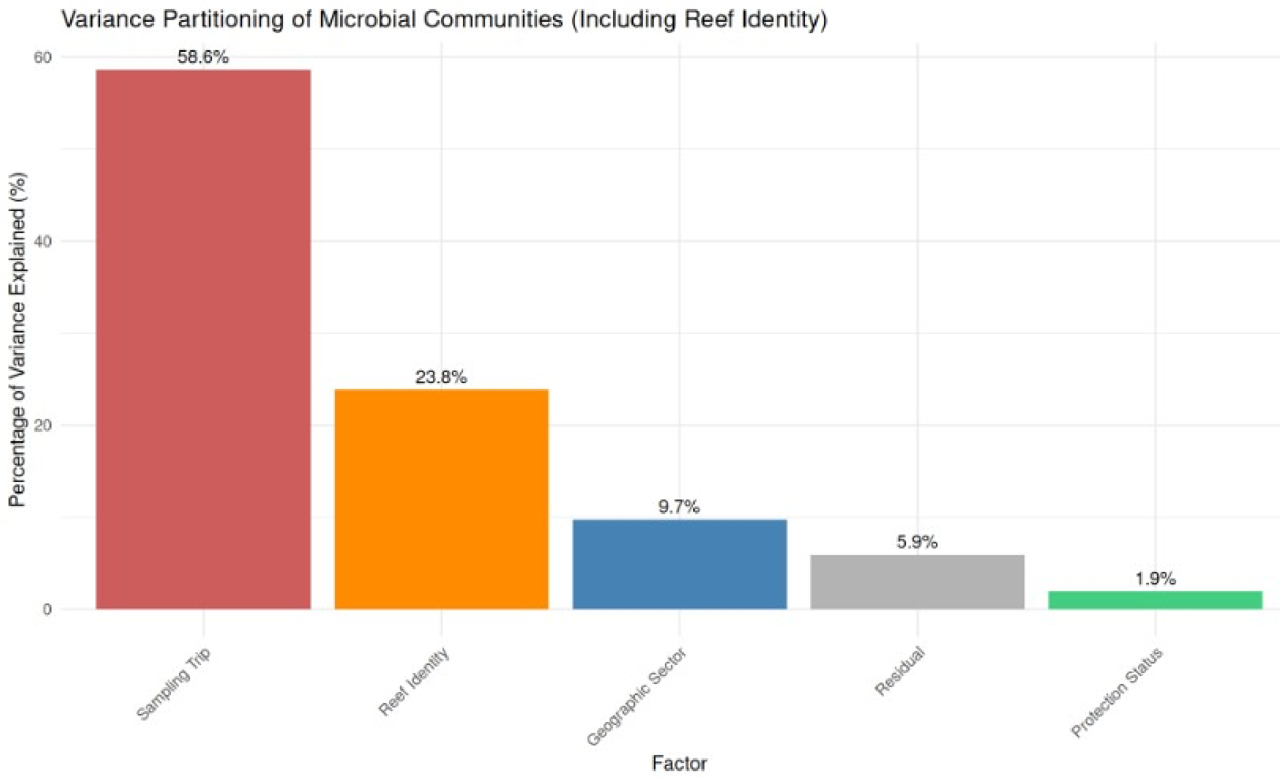


**Figure S3. PERmutational Multivariate ANalysis Of VAriance (PERMANOVA) Variance Partitioning**. Proportion of variance in seawater microbial communities explained by different factors, with sampling trip explaining most of variance (58.6%; $R^2 = 0.58616$, $F = 467.789$, $p < 0.001$), followed by reef site (23.8% of variance; $R^2 = 0.23833$, $F = 14.631$, $p < 0.001$), Great Barrier Reef sector (9.7% of variance; $R^2 = 0.09712$, $F = 58.133$, $p < 0.001$), and lastly, reef protection status explaining 1.91% of variance ($R^2 = 0.01907$, $F = 45.648$, $p < 0.001$). Total explained variance: 94.1% - Residual/unexplained variance: 5.9%.

**Table S1**. PERMANOVA results testing the effects of reef protection status (as predictor variable) and sampling trip, sector, and reef name (as covariates) on seawater microbial community composition.

| Term | Df | SumOfSqs | $R^2$ | F | p.value | Significance |
|---|---|---|---|---|---|---|
| Open_or_Closed_to_fishing | 1 | 9464.96 | 0.01907 | 45.648 | < 0.001 | *** |
| Sampling_trip | 3 | 290984.89 | 0.58616 | 467.789 | < 0.001 | *** |
| SECTOR_N_S | 4 | 48214.63 | 0.09712 | 58.133 | < 0.001 | *** |
| REEF_NAME | 39 | 118314.20 | 0.23833 | 14.631 | < 0.001 | *** |
| Residual | 142 | 29443.39 | 0.05931 | NA | NA | |
| Total | 189 | 496422.06 | 1.00000 | NA | NA | |

### Distance-Based ReDundancy Analysis - dbRDA

To complement PERMANOVA and visualise the constrained ordination space, we performed Distance-

Based ReDundancy Analysis (dbRDA; dbrda() function in vegan[10,11] v2.6-4). Unlike PERMANOVA which tests statistical significance, dbRDA provides a visual representation of how community composition varies along gradients defined by explanatory variables. To avoid overconstraining the model (which occurred when including reef identity as 47 separate levels), we constructed a simpler model focusing on broader-scale patterns: aitchison_dist ~ Open_or_Closed_to_fishing + Sampling_trip + SECTOR_N_S. This model explained 70.2% of total variance across 8 constrained axes (**Fig. S4**), with sampling trip explaining 58.6%, geographic sector 9.7%, and reef protection status 1.9% of variance. All factors were statistically significant ($p < 0.001$). The dbRDA ordination revealed clear seasonal separation along the first axis, with partial separation between protected and fished reefs, particularly within seasonal clusters.

This influence of reef zoning on community-level structuring of microbial communities provided the basis for downstream feature selection analyses.

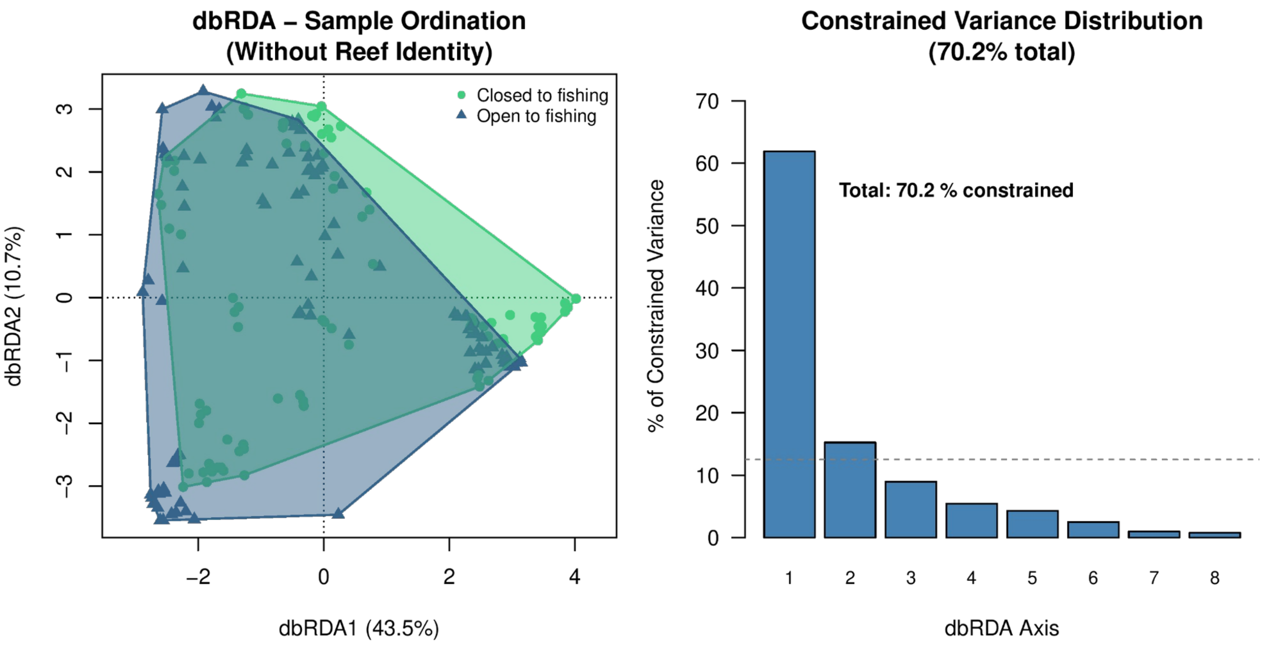


**Figure S4. Distance-Based ReDundancy Analysis (dbRDA) ordination of seawater microbial communities**. (**Left**) dbRDA ordination showing samples colored by protection status (green = closed to fishing; blue = open to fishing). Convex hulls represent group boundaries. The first two constrained axes explain 54.2% of constrained variance (dbRDA1: 43.5%, dbRDA2: 10.7%). (**Right**) Total variance (70.2%) explained by each constrained axis, showing rapid decay after the first two axes.

### (s)PLS-DA - (Sparse) Partial Least Squares Discriminant Analysis | Can we discriminate between No-Take Marine Reserves (NTMRs) and fished reefs using a supervised approach?

#### PLS-DA

As PCA ordination shows that our sites cluster based on geographic proximity (i.e. sector) and time (i.e.

sampling trip) (**Fig. S1**) and not based on reef zoning (**Fig. S2**) as our categorical outcome of interest, we then explored if sPLS-DA[12], as a supervised approach, will identify microbial indicators of No-Take Marine Reserves (NTMRs) vs fished reefs.

#### Tuning the number of components in PLS-DA

The perf() function evaluates the performance of PLS-DA - i.e., its ability to rightly classify 'new' samples into their category (NTMRs and fished zones) using repeated cross-validation. We initially choose a large number of components (here ncomp = 10) and assess the model as we gradually increase the number of components. Here, we used a 4-fold CV repeated 50 times.

The plot (**Fig. S5**) shows that the error rate keeps dropping as we increase the number of components, which may suggest strong batch effects in the data as this many dimensions would typically not be needed to discriminate only two categorical outcomes (NTMRs and fished reefs).

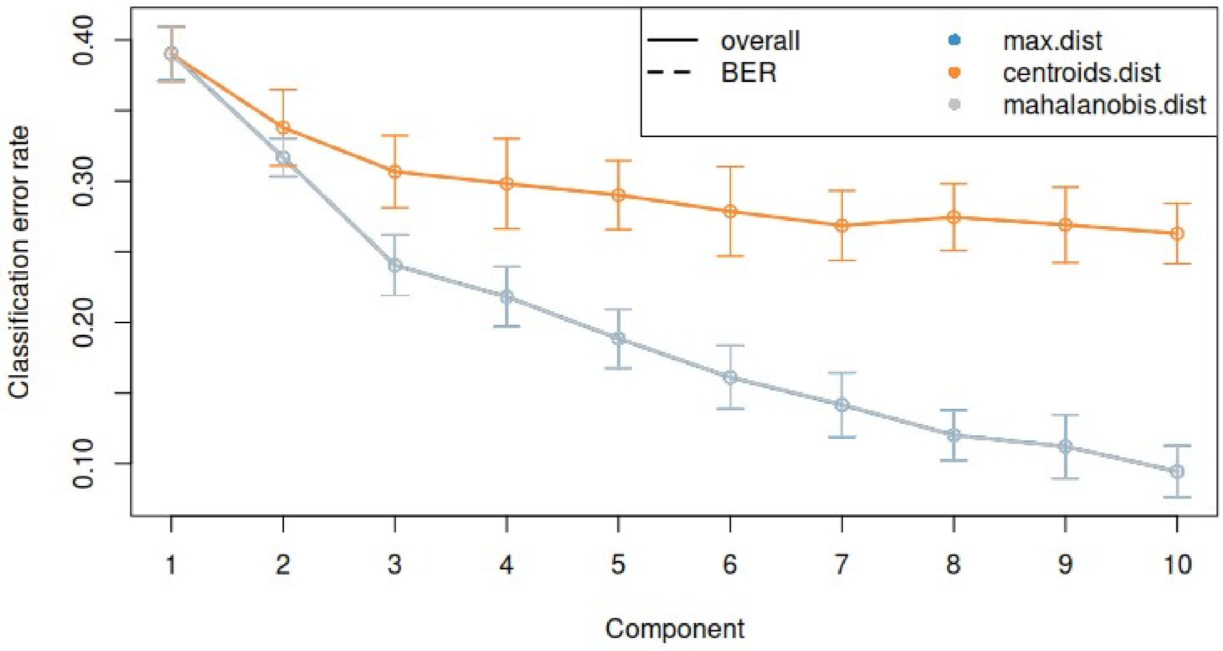


**Figure S5. Tuning the number of components in PLS-DA on the IMOS GBR-MGD microbial data (876 pMAGs, de-replicated at 95% ANI)**. For each component, repeated cross-validation (50 ×4–fold CV) is used to evaluate the PLS-DA classification performance (overall and balanced error rate BER, and for each type of prediction distance: max.dist, centroids.dist and mahalanobis.dist) to discriminate between No-Take Marine Reserves (NTMRs) and fished reefs based on the seawater microbiomes. Bars show the standard deviation across the repeated folds.

In PLS-DA sample plots (**Fig. S6**), we can observe improved clustering according to reef protection status (**Fig. S6**; top), compared with PCA (**Fig. S2**). This is to be expected since PLS-DA is a supervised approach and includes the class information of each sample, and aims to discriminate between them. From the *plotIndiv()* function, we observe some discrimination between NTMRs and fished reefs mostly on component 1 (x-axis), however we can still see the trip effect (**Fig. S6**; bottom). The axis labels indicate the amount of variation explained per component, however, the interpretation of this amount is not as important as in PCA, as PLS-DA aims to maximise the covariance between components associated to X (predictor dataset, i.e. the 876 IMOS-MGD MAGs) and Y (categorical "response", i.e. no-take and take zones), rather than the variance of X (shown in PCA plots).

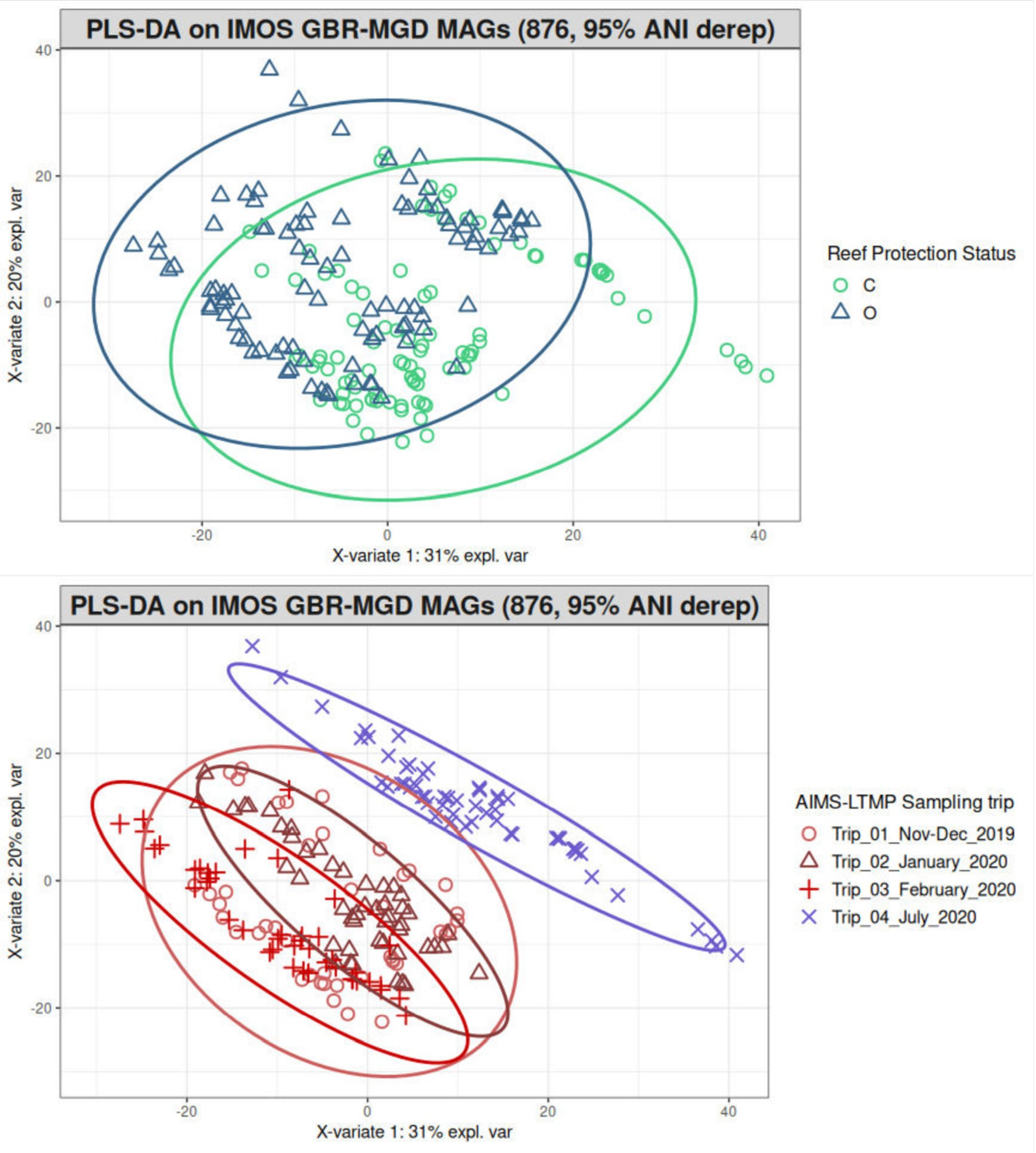


**Figure S6. Sample plots from PLS-DA performed on the IMOS GBR-MGD microbial data (876 pMAGs, dereplicated at 95% ANI) as X, to discriminate reef zoning status as Y**. Samples are projected into the space spanned by the first two components, coloured by their reef zoning status (above) or sampling trip (below). While we do observe separation of samples based on reef zoning (above), samples also cluster based on time of sampling and geographic proximity (below), suggesting batch effects due to confounding effects of space and time.

### sPLS-DA - can we refine these clusters by selecting the most influential pMAGs to classify reef zoning?

As many of the pMAGs in X may be noisy or uninformative to discriminate between NTMRs and fished reefs, an sPLS-DA analysis (sparse variant) may help refine the sample clusters and select a small subset of variables relevant to discriminate each class.

#### Tuning the number of variables to select

We estimate the classification error rate with respect to the number of selected variables in the model with the function *tune.splsda()*. The tuning is being performed one component at a time inside the function and the optimal number of variables to select is automatically retrieved after each component run.

Previously, we determined the optimal number of components to be ncomp = 10 with PLS-DA. Here we set ncomp = 15 to further assess if this would be the case for a sparse model, and use 4-fold cross validation repeated 50 times. We first define a grid of keepX values, and we tested 34 keepX values in total: a fine grid (1-10) followed by a coarser sequence (20-250 in increments of 10).

In (**Fig. S7**), we display the mean classification error rate on each component, bearing in mind that each component is conditional on the previous components calculated with the optimal number of selected variables. The diamond in the figure below indicates the best keepX value to achieve the lowest error rate per component. This type of graph helps not only to choose the 'optimal' number of variables to select, but also to confirm the number of components ncomp. From the following code (tune.splsda.open.closed_IMOS_P$choice.ncomp$ncomp), we can assess that the optimal number of components was 10 according to a one-sided T−test.

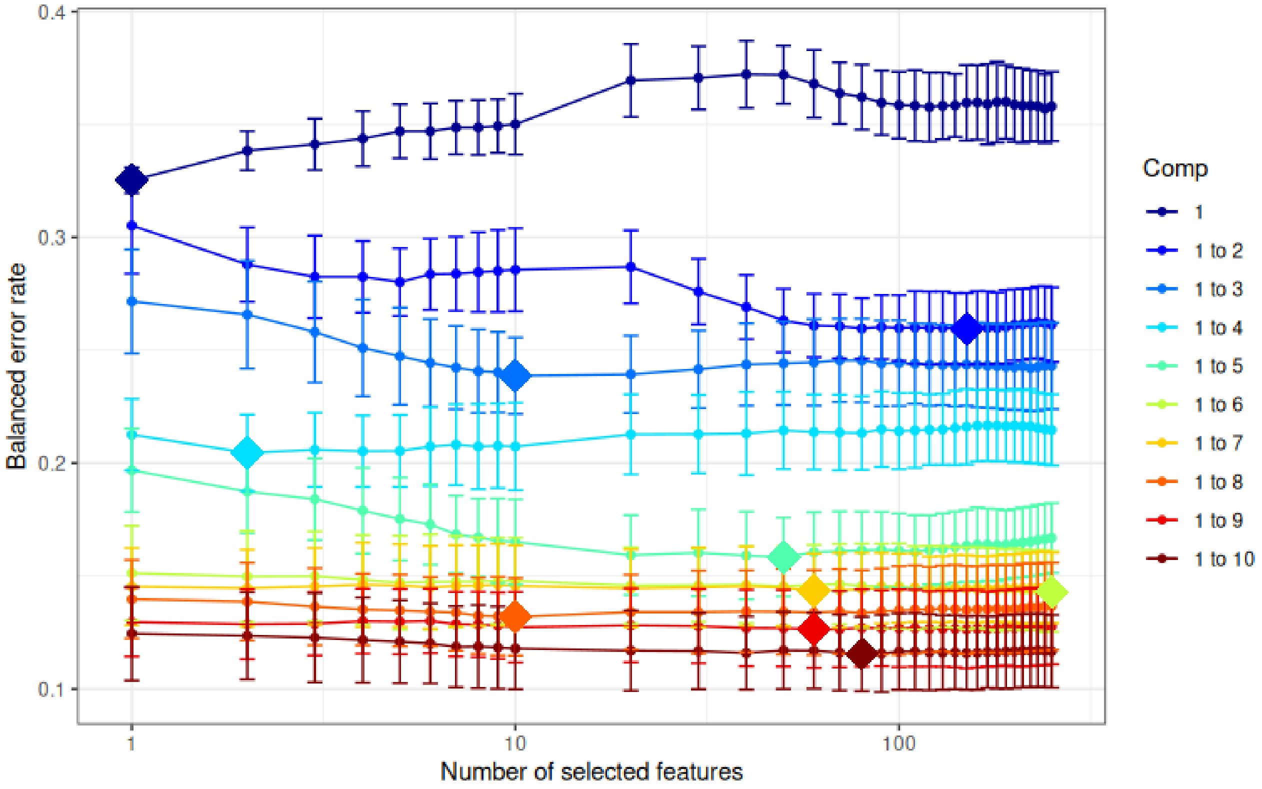


**Figure S7. Tuning keepX for the sPLS-DA performed on the IMOS GBR-MGD pMAGs (876 genomes, drep at 95% ANI).** Each coloured line represents the balanced error rate (y-axis) per component across all tested keepX values (x-axis) with the standard deviation based on the repeated cross-validation folds (4-fold x 50 repeats). The diamond indicates the optimal keepX value on a particular component which achieves the lowest classification error rate as determined with a one-sided t–test. As sPLS-DA is an iterative algorithm, values represented for a given component (e.g. comp 1 to 2) include the optimal keepX value chosen for the previous component (comp 1).

The numerical output from *tune.splsda()* (**Table S2**) globally shows that the classification error rate continues to decrease after the second component in sparse PLS-DA, yet since we are only discriminating between two categorical outcomes, retaining a small number (i.e. one or two) components is recommended to avoid overfitting. In (**Figs. S8-S9**), we further confirm the spatiotemporal batch effects in the data.

**Table S2. Numerical output associated with Fig. S7**, showing sPLS-DA tune() results: mean error rate for each component and each tested keepX value given the past (tuned) components.

| comp1 | comp2 | comp3 | comp4 | comp5 | comp6 | comp7 | comp8 | comp9 | comp10 |
|---|---|---|---|---|---|---|---|---|---|
| 0.33 | 0.31 | 0.27 | 0.21 | 0.20 | 0.15 | 0.15 | 0.14 | 0.13 | 0.12 |
| 0.34 | 0.29 | 0.27 | 0.20 | 0.19 | 0.15 | 0.14 | 0.14 | 0.13 | 0.12 |
| 0.34 | 0.28 | 0.26 | 0.21 | 0.18 | 0.15 | 0.15 | 0.14 | 0.13 | 0.12 |
| 0.34 | 0.28 | 0.25 | 0.21 | 0.18 | 0.15 | 0.15 | 0.14 | 0.13 | 0.12 |
| 0.35 | 0.28 | 0.25 | 0.21 | 0.18 | 0.15 | 0.15 | 0.13 | 0.13 | 0.12 |
| 0.35 | 0.28 | 0.24 | 0.21 | 0.17 | 0.15 | 0.14 | 0.13 | 0.13 | 0.12 |

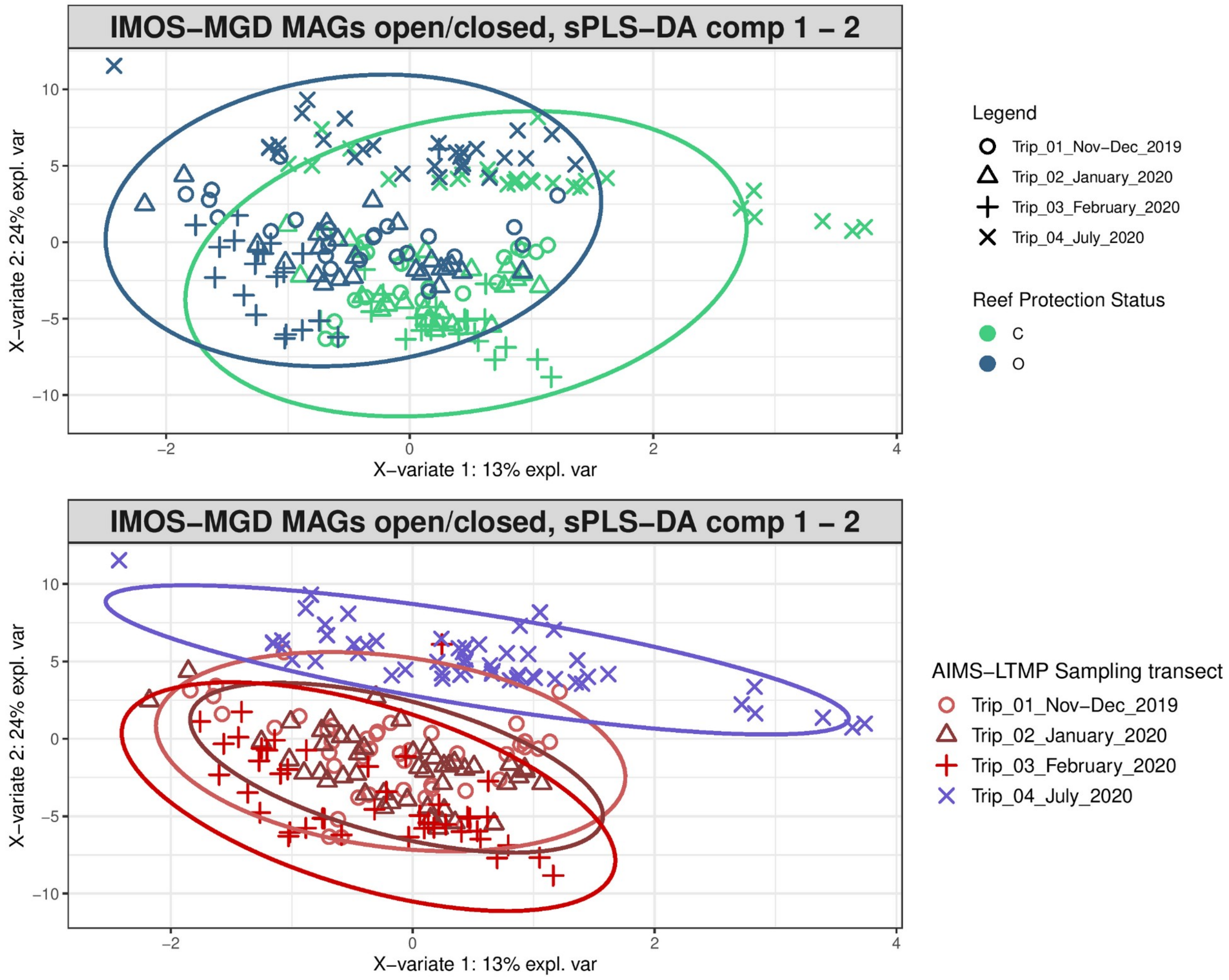


**Figure S8. Sample plots from sPLS-DA performed on the IMOS GBR-MGD microbial data (876 pMAGs, dereplicated at 95% ANI) as X, to discriminate reef zoning status as Y**. Samples are projected into the space spanned by the first two components. The plots represent 95% ellipse confidence intervals around each sample class: (above) NTMRs (in green) vs fished reefs (in blue); and (below) sampling trip. While we do see separation between NTMRs and fished reefs as our outcome of interest (above), we can also observe spatio-temporal batch effects (below).

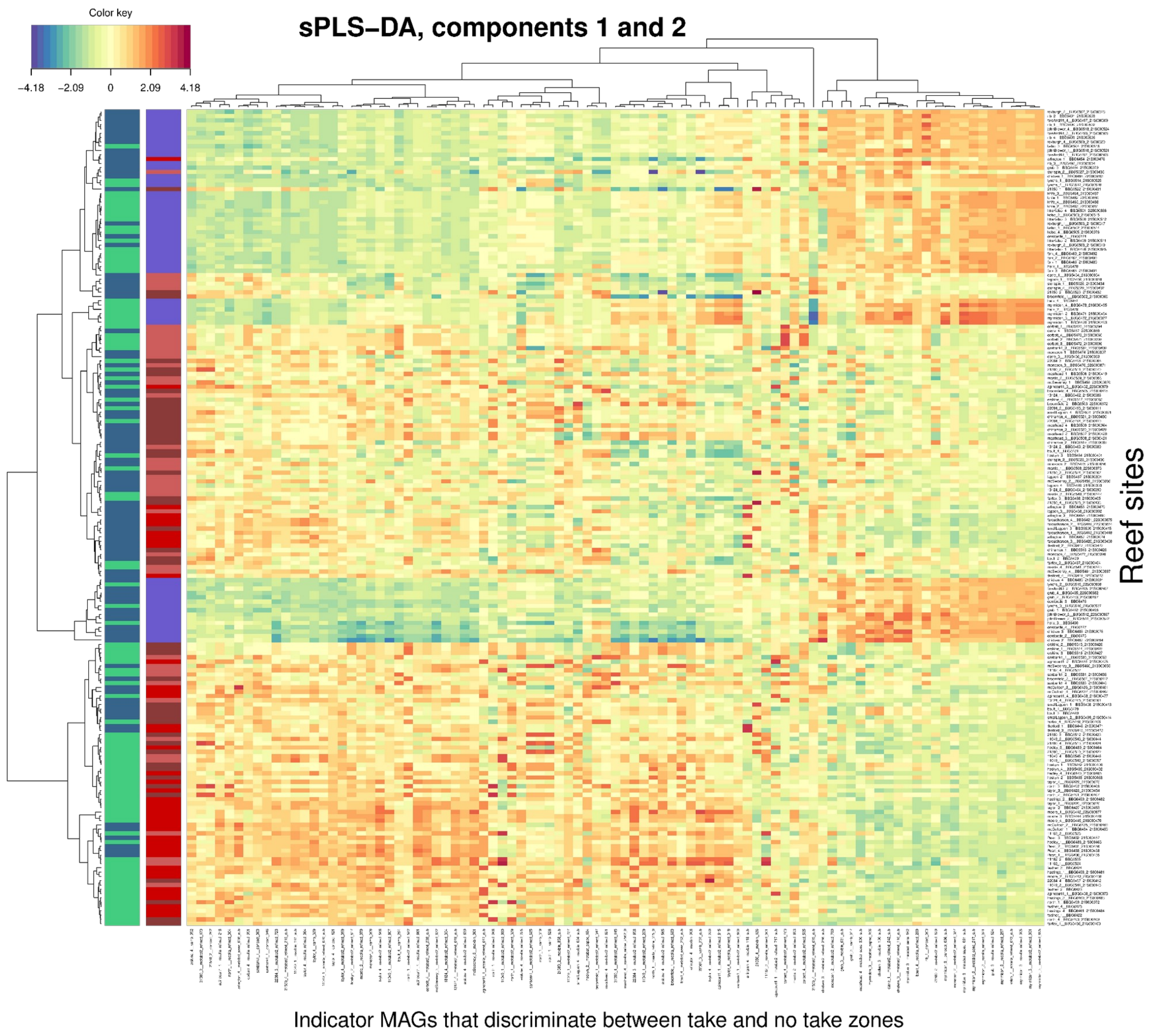


**Figure S9. Clustered Image Map showing abundance patterns of reef zoning microbial indicators, as inferred from sPLS-DA performed on the IMOS GBR-MGD microbial data (876 pMAGs, de-replicated at 95% ANI) as X, to discriminate No-Take Marine Reserves (NTMRs) and fished reefs as categorical Y**. A hierarchical clustering based on the pMAG enrichment values for the selected indicator microbes, with samples (48 reef sites x 4 replicates) in rows coloured according to their reef protection status (NTMRs in green, fished reefs in blue) and sampling trip (Austral summer/wet season samples in red tones, winter samples in blue). The heatmap is clustered using Euclidean distance with Complete agglomeration method. As previously observed, spatio-temporal patterns are stronger drivers than zoning and thus represent batch effects that need to be corrected for.

### **M**ultivariate **INT**egration **(MINT) sPLS-DA** | Discriminating between reefs that are open or closed to fishing (**sPLS-DA**), while accounting for sector-specific effects (**MINT**)

#### Tuning the number of dimensions

The perf() function is used to estimate the performance of the MINT-sPLS-DA model using Leave One Group Out Cross Validation (LOGOCV, i.e. by training MINT sPLS-DA on six out of seven sectors and validating the model performance on the left-out subset, hence seven times until each of the seven GBR sectors is left out once), and to choose the optimal number of components for our final model.

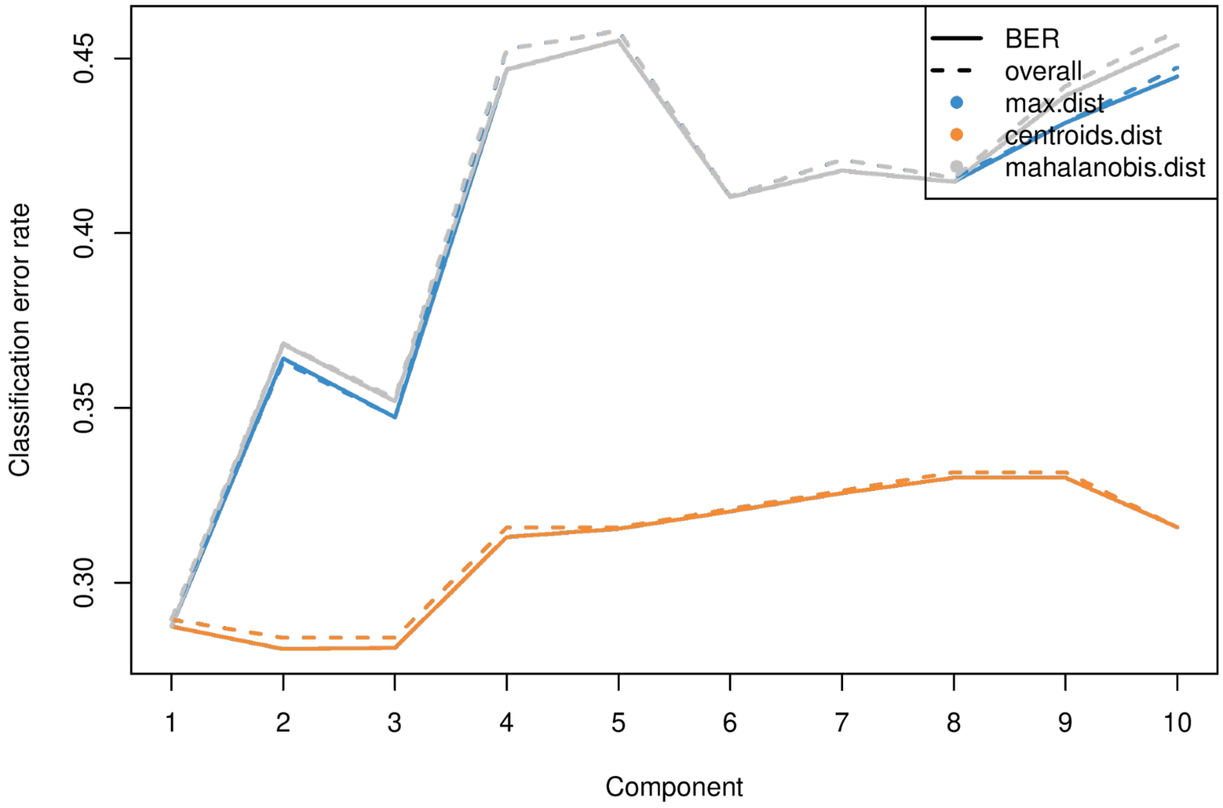


**Figure S10. Choosing the number of components in mint.splsda using perf() with LOGOCV to discriminate between reefs that are open or closed to fishing, using a dataset of 876 IMOS GBR-MGD microbial genomes (pMAGs$_{95\%ANI}$).** Classification error rates (overall and balanced - BER) are represented on the y-axis with respect to the number of components on the x-axis for each prediction distance. Overall and balanced error rates show largely the same trend as the design is balanced (i.e. the same number of NTMR and fished reefs in each GBR sector). The plot shows that the error rate reaches a minimum (~29%) with two or three dimensions with the centroids prediction distance. We therefore retained 2 PCs in downstream analysis.

**Table S3. Numerical output associated with Fig. S10**. Here, we show overall MINT sPLS-DA error rates when discriminating between reefs that are open or closed to fishing (Y), using 876 IMOS GBR-MGD pMAGs$_{95\%ANI}$ as X, using 3 prediction distances (max.dist, centroids.dist, mahalanobis.dist), and across the seven GBR sectors. Sector-specific model accuracies were expressed as 1 – error (with centroids dist), and for each of the 10 tested MINT sPLS-DA components, we also show MINT sPLS-DA classification accuracy averaged across 7 GBR sectors.

| MINT sPLS-DA comp | Study (GBR sector) | Max dist | Centroids dist | Mahalanobis dist | Accuracy (1 – error with centroids dist) | Average accuracy |
|---|---|---|---|---|---|---|
| comp1 | 01_Cape_Grenville | 0.42 | 0.42 | 0.42 | 0.58 | |
| comp1 | 02_Princess_Charlotte_bay | 0.20 | 0.20 | 0.20 | 0.80 | |
| comp1 | 03_Cairns | 0.35 | 0.35 | 0.35 | 0.65 | |
| comp1 | 04_Innisfail | 0.07 | 0.07 | 0.07 | 0.93 | 0.71 |
| comp1 | 05_Townsville | 0.30 | 0.30 | 0.30 | 0.70 | |
| comp1 | 06_Swains | 0.35 | 0.35 | 0.35 | 0.65 | |
| comp1 | 07_Capricorn_Bunker | 0.32 | 0.32 | 0.32 | 0.68 | |
| comp2 | 01_Cape_Grenville | 0.58 | 0.42 | 0.54 | 0.58 | 0.71 |
| comp2 | 02_Princess_Charlotte_bay | 0.67 | 0.20 | 0.73 | 0.80 | |
| comp2 | 03_Cairns | 0.10 | 0.25 | 0.10 | 0.75 | |
| comp2 | 04_Innisfail | 0.22 | 0.15 | 0.22 | 0.85 | |

| | | | | | | |
|---|---|---|---|---|---|---|
| comp2 | 05_Townsville | 0.32 | 0.27 | 0.32 | 0.73 | |
| comp2 | 06_Swains | 0.45 | 0.40 | 0.50 | 0.60 | |
| comp2 | 07_Capricorn_Bunker | 0.36 | 0.32 | 0.36 | 0.68 | |
| comp3 | 01_Cape_Grenville | 0.63 | 0.38 | 0.63 | 0.63 | |
| comp3 | 02_Princess_Charlotte_bay | 0.60 | 0.27 | 0.60 | 0.73 | |
| comp3 | 03_Cairns | 0.25 | 0.25 | 0.25 | 0.75 | |
| comp3 | 04_Innisfail | 0.30 | 0.15 | 0.30 | 0.85 | 0.71 |
| comp3 | 05_Townsville | 0.30 | 0.29 | 0.30 | 0.71 | |
| comp3 | 06_Swains | 0.35 | 0.40 | 0.35 | 0.60 | |
| comp3 | 07_Capricorn_Bunker | 0.18 | 0.29 | 0.21 | 0.71 | |
| comp4 | 01_Cape_Grenville | 0.63 | 0.42 | 0.63 | 0.58 | |
| comp4 | 02_Princess_Charlotte_bay | 0.93 | 0.27 | 0.93 | 0.73 | |
| comp4 | 03_Cairns | 0.25 | 0.30 | 0.25 | 0.70 | |
| comp4 | 04_Innisfail | 0.48 | 0.22 | 0.48 | 0.78 | 0.68 |
| comp4 | 05_Townsville | 0.39 | 0.29 | 0.39 | 0.71 | |
| comp4 | 06_Swains | 0.45 | 0.40 | 0.45 | 0.60 | |
| comp4 | 07_Capricorn_Bunker | 0.29 | 0.36 | 0.29 | 0.64 | |
| comp5 | 01_Cape_Grenville | 0.71 | 0.42 | 0.71 | 0.58 | 0.67 |

| | | | | | | |
|---|---|---|---|---|---|---|
| comp5 | 02_Princess_Charlotte_bay | 1.00 | 0.33 | 1.00 | 0.67 | |
| comp5 | 03_Cairns | 0.30 | 0.30 | 0.30 | 0.70 | |
| comp5 | 04_Innisfail | 0.48 | 0.19 | 0.48 | 0.81 | |
| comp5 | 05_Townsville | 0.38 | 0.29 | 0.38 | 0.71 | |
| comp5 | 06_Swains | 0.45 | 0.40 | 0.45 | 0.60 | |
| comp5 | 07_Capricorn_Bunker | 0.21 | 0.36 | 0.21 | 0.64 | |
| comp6 | 01_Cape_Grenville | 0.63 | 0.42 | 0.63 | 0.58 | |
| comp6 | 02_Princess_Charlotte_bay | 0.80 | 0.40 | 0.80 | 0.60 | |
| comp6 | 03_Cairns | 0.30 | 0.30 | 0.30 | 0.70 | |
| comp6 | 04_Innisfail | 0.41 | 0.19 | 0.41 | 0.81 | 0.67 |
| comp6 | 05_Townsville | 0.34 | 0.29 | 0.34 | 0.71 | |
| comp6 | 06_Swains | 0.40 | 0.40 | 0.40 | 0.60 | |
| comp6 | 07_Capricorn_Bunker | 0.25 | 0.36 | 0.25 | 0.64 | |
| comp7 | 01_Cape_Grenville | 0.63 | 0.46 | 0.63 | 0.54 | 0.66 |
| comp7 | 02_Princess_Charlotte_bay | 0.87 | 0.40 | 0.87 | 0.60 | |
| comp7 | 03_Cairns | 0.20 | 0.30 | 0.20 | 0.70 | |
| comp7 | 04_Innisfail | 0.41 | 0.19 | 0.41 | 0.81 | |
| comp7 | 05_Townsville | 0.34 | 0.29 | 0.34 | 0.71 | |
| comp7 | 06_Swains | 0.45 | 0.40 | 0.45 | 0.60 | |

| | | | | | | |
|---|---|---|---|---|---|---|
| comp7 | 07_Capricorn_Bunker | 0.32 | 0.36 | 0.32 | 0.64 | |
| comp8 | 01_Cape_Grenville | 0.63 | 0.46 | 0.63 | 0.54 | |
| comp8 | 02_Princess_Charlotte_bay | 0.93 | 0.40 | 0.93 | 0.60 | |
| comp8 | 03_Cairns | 0.20 | 0.30 | 0.20 | 0.70 | |
| comp8 | 04_Innisfail | 0.44 | 0.19 | 0.44 | 0.81 | 0.65 |
| comp8 | 05_Townsville | 0.32 | 0.29 | 0.32 | 0.71 | |
| comp8 | 06_Swains | 0.35 | 0.45 | 0.35 | 0.55 | |
| comp8 | 07_Capricorn_Bunker | 0.32 | 0.36 | 0.32 | 0.64 | |
| comp9 | 01_Cape_Grenville | 0.67 | 0.46 | 0.67 | 0.54 | |
| comp9 | 02_Princess_Charlotte_bay | 0.93 | 0.40 | 0.93 | 0.60 | |
| comp9 | 03_Cairns | 0.20 | 0.30 | 0.20 | 0.70 | |
| comp9 | 04_Innisfail | 0.44 | 0.19 | 0.52 | 0.81 | 0.65 |
| comp9 | 05_Townsville | 0.30 | 0.29 | 0.30 | 0.71 | |
| comp9 | 06_Swains | 0.35 | 0.45 | 0.40 | 0.55 | |
| comp9 | 07_Capricorn_Bunker | 0.43 | 0.36 | 0.39 | 0.64 | |
| comp10 | 01_Cape_Grenville | 0.75 | 0.46 | 0.75 | 0.54 | 0.67 |
| comp10 | 02_Princess_Charlotte_bay | 0.80 | 0.33 | 0.80 | 0.67 | |
| comp10 | 03_Cairns | 0.20 | 0.30 | 0.20 | 0.70 | |

| | | | | | |
|---|---|---|---|---|---|
| comp10 | 04_Innisfail | 0.56 | 0.19 | 0.56 | 0.81 |
| comp10 | 05_Townsville | 0.29 | 0.29 | 0.29 | 0.71 |
| comp10 | 06_Swains | 0.50 | 0.45 | 0.55 | 0.55 |
| comp10 | 07_Capricorn_Bunker | 0.36 | 0.29 | 0.39 | 0.71 |

**Table S4. MINT sPLS-DA - error rate (centroids distance) across GBR sectors, and separately for C (reefs closed to fishing) and O (open to fishing).**

| | Study | comp1 | comp2 | comp3 | comp4 | comp5 | comp6 | comp7 | comp8 | comp9 | comp10 |
|---|---|---|---|---|---|---|---|---|---|---|---|
| C | 01_Cape_Grenville | 0.58 | 0.58 | 0.58 | 0.58 | 0.58 | 0.58 | 0.58 | 0.58 | 0.58 | 0.58 |
| O | 01_Cape_Grenville | 0.25 | 0.25 | 0.17 | 0.25 | 0.25 | 0.25 | 0.33 | 0.33 | 0.33 | 0.33 |
| C | 02_Princess_Charlotte_bay | 0.00 | 0.00 | 0.13 | 0.13 | 0.25 | 0.38 | 0.38 | 0.38 | 0.38 | 0.25 |
| O | 02_Princess_Charlotte_bay | 0.43 | 0.43 | 0.43 | 0.43 | 0.43 | 0.43 | 0.43 | 0.43 | 0.43 | 0.43 |
| C | 03_Cairns | 0.25 | 0.00 | 0.00 | 0.13 | 0.13 | 0.13 | 0.13 | 0.13 | 0.13 | 0.13 |
| O | 03_Cairns | 0.42 | 0.42 | 0.42 | 0.42 | 0.42 | 0.42 | 0.42 | 0.42 | 0.42 | 0.42 |
| C | 04_Innisfail | 0.07 | 0.13 | 0.13 | 0.20 | 0.13 | 0.13 | 0.13 | 0.13 | 0.13 | 0.13 |
| O | 04_Innisfail | 0.08 | 0.17 | 0.17 | 0.25 | 0.25 | 0.25 | 0.25 | 0.25 | 0.25 | 0.25 |
| C | 05_Townsville | 0.39 | 0.32 | 0.36 | 0.36 | 0.36 | 0.36 | 0.36 | 0.36 | 0.36 | 0.36 |
| O | 05_Townsville | 0.21 | 0.21 | 0.21 | 0.21 | 0.21 | 0.21 | 0.21 | 0.21 | 0.21 | 0.21 |
| C | 06_Swains | 0.25 | 0.38 | 0.38 | 0.38 | 0.50 | 0.50 | 0.50 | 0.50 | 0.50 | 0.50 |
| O | 06_Swains | 0.42 | 0.42 | 0.42 | 0.42 | 0.33 | 0.33 | 0.33 | 0.42 | 0.42 | 0.42 |
| C | 07_Capricorn_Bunker | 0.31 | 0.31 | 0.25 | 0.38 | 0.38 | 0.38 | 0.38 | 0.38 | 0.38 | 0.25 |
| O | 07_Capricorn_Bunker | 0.33 | 0.33 | 0.33 | 0.33 | 0.33 | 0.33 | 0.33 | 0.33 | 0.33 | 0.33 |

| | | | | | | | | | | |
|---|---|---|---|---|---|---|---|---|---|---|
| Average error [C] | 0.27 | 0.25 | 0.26 | 0.31 | 0.33 | 0.35 | 0.35 | 0.35 | 0.35 | 0.31 |
| Average error [O] | 0.31 | 0.32 | 0.31 | 0.33 | 0.32 | 0.32 | 0.33 | 0.34 | 0.34 | 0.34 |
| Average accuracy [C] | 0.73 | 0.75 | 0.74 | 0.69 | 0.67 | 0.65 | 0.65 | 0.65 | 0.65 | 0.69 |
| Average accuracy [O] | 0.69 | 0.68 | 0.69 | 0.67 | 0.68 | 0.68 | 0.67 | 0.66 | 0.66 | 0.66 |

### Tuning the number of features per dimension

We can choose the keepX parameter using the *tune()* function for a MINT object. The function performs LOGOCV for different values of test.keepX (we specified test.keepX = seq(10, 300, 10)) provided on each component (we tested 5 components), and no repeat argument is needed. Based on the mean classification error rate (overall error rate or BER) and a centroids distance, we output the optimal number of variables keepX to be included in the final model.

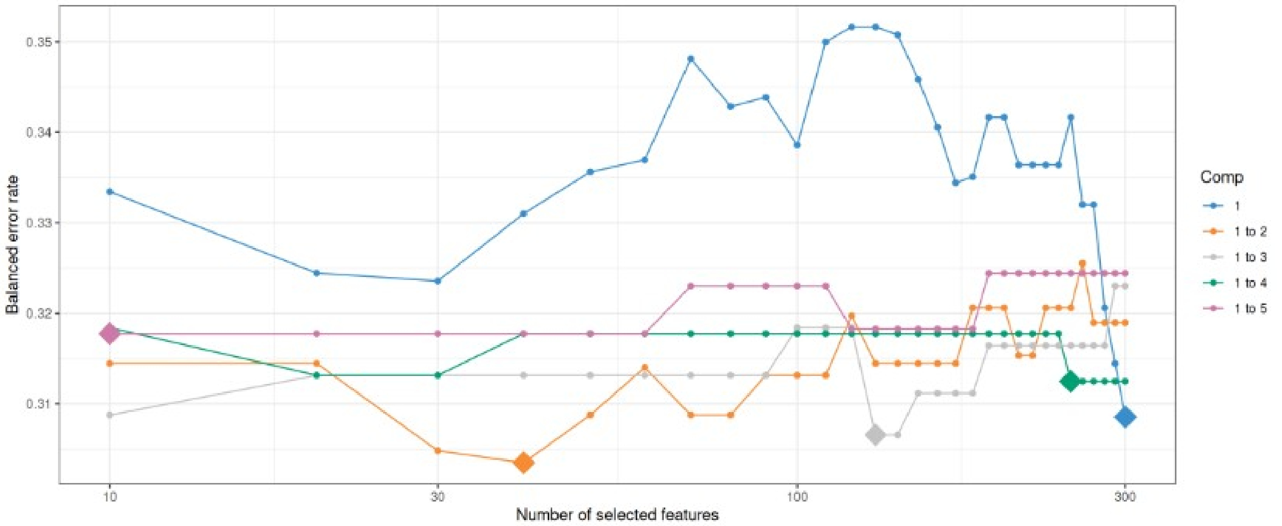


**Figure S11. Tuning plot of the MINT sPLS-DA models with up to 5 components, testing a grid value of 10 to 300 indicators (with sequential increases of 10)**. Diamonds represent the optimal number of features on a given component. Balanced error rate found on the vertical axis and is the metric to be minimised.

**Table S5. Numerical output associated with Fig S11**, also showing sector-specific MINT sPLS-DA classification errors and average error/accuracy across sectors.

| GBR_sector | **comp1** | **comp2** | **comp3** | **comp4** | **comp5** |
|---|---|---|---|---|---|

| | | | | | |
|---|---|---|---|---|---|
| CA | 0.33 | 0.27 | 0.21 | 0.27 | 0.27 |
| CB | 0.32 | 0.32 | 0.32 | 0.35 | 0.35 |
| CG | 0.42 | 0.42 | 0.46 | 0.42 | 0.46 |
| IN | 0.18 | 0.23 | 0.23 | 0.23 | 0.23 |
| PC | 0.21 | 0.21 | 0.21 | 0.21 | 0.21 |
| SW | 0.44 | 0.4 | 0.44 | 0.44 | 0.44 |
| TO | 0.29 | 0.29 | 0.29 | 0.29 | 0.29 |
| Average Error | 0.31 | 0.31 | 0.31 | 0.32 | 0.32 |
| Accuracy (1 − average error) | 0.69 | 0.69 | 0.69 | 0.68 | 0.68 |

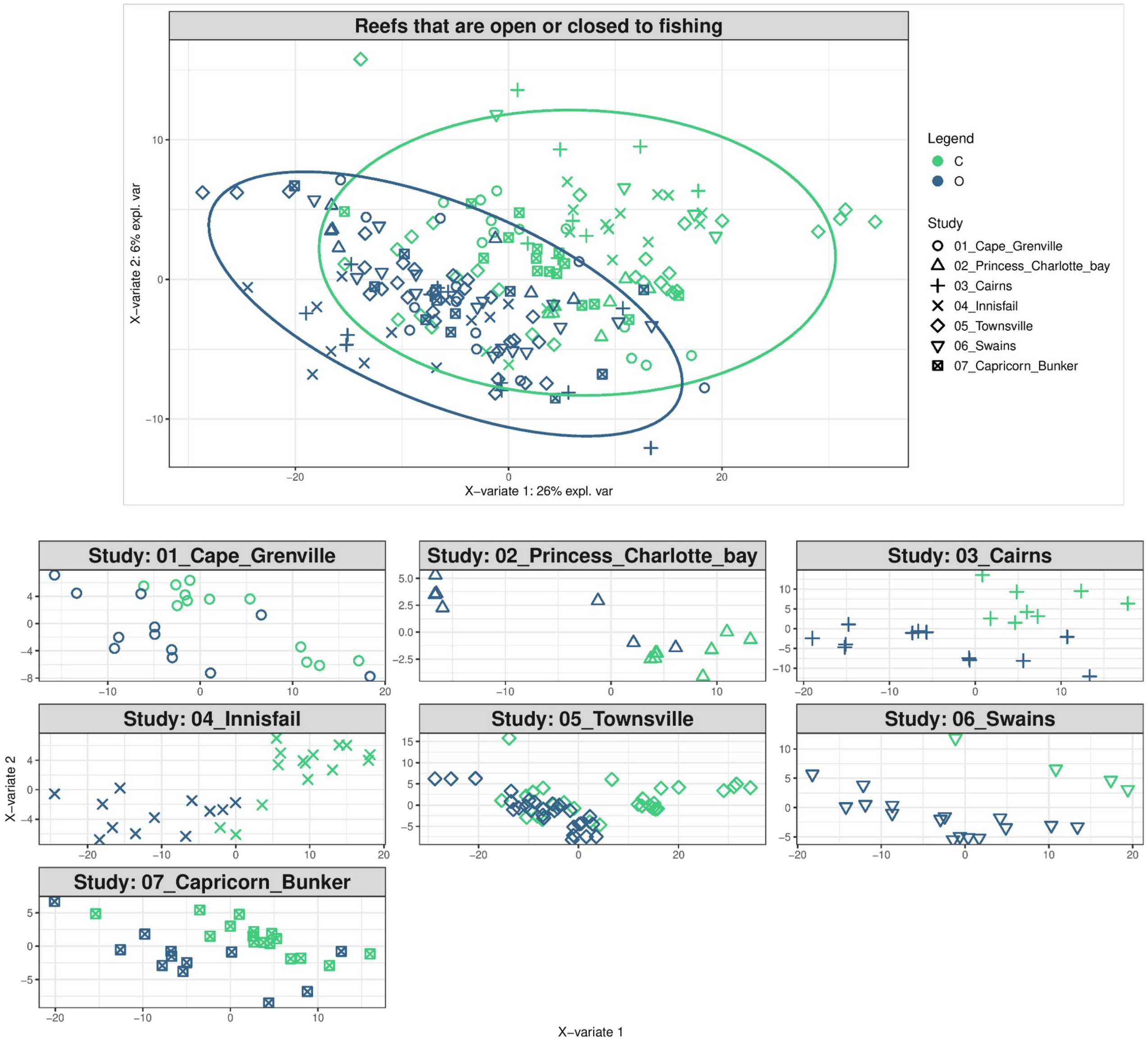


**Figure S12. Sample plots from the MINT sPLS-DA performed on the 876 IMOS GBR-MGD seawater $pMAGs_{95\%ANI}$, aiming to find discriminatory microbes between reefs that are open or closed to fishing**. Samples (48 reef sites x 4 replicates) are projected into the space spanned by the first two components. Reef sites are coloured by their protection level (open or closed to fishing) and symbols indicate the membership of reef sites to their corresponding LTMP trip/transect. (**top**) Global components from the model with 95% ellipse confidence intervals around each sample class. (**bottom**) Partial components per study show a good agreement across GBR sectors. Component 1 discriminates between reefs that are open or closed to fishing.

### Performance of the final MINT sPLS-DA model

Use of the auroc() function will yield a visualisation of classification performance when undergoing the LOGOCV procedure from above. The interpretation of this output may not be particularly insightful in relation to the performance evaluation of mixOmics methods, but can complement the statistical analysis. For example, the MINT sPLS-DA model of fished vs. NTMR sites yielded an AUC score of 0.73 (**Fig. S13**), which complements the overall ~71% classification accuracy observed during cross-validation.

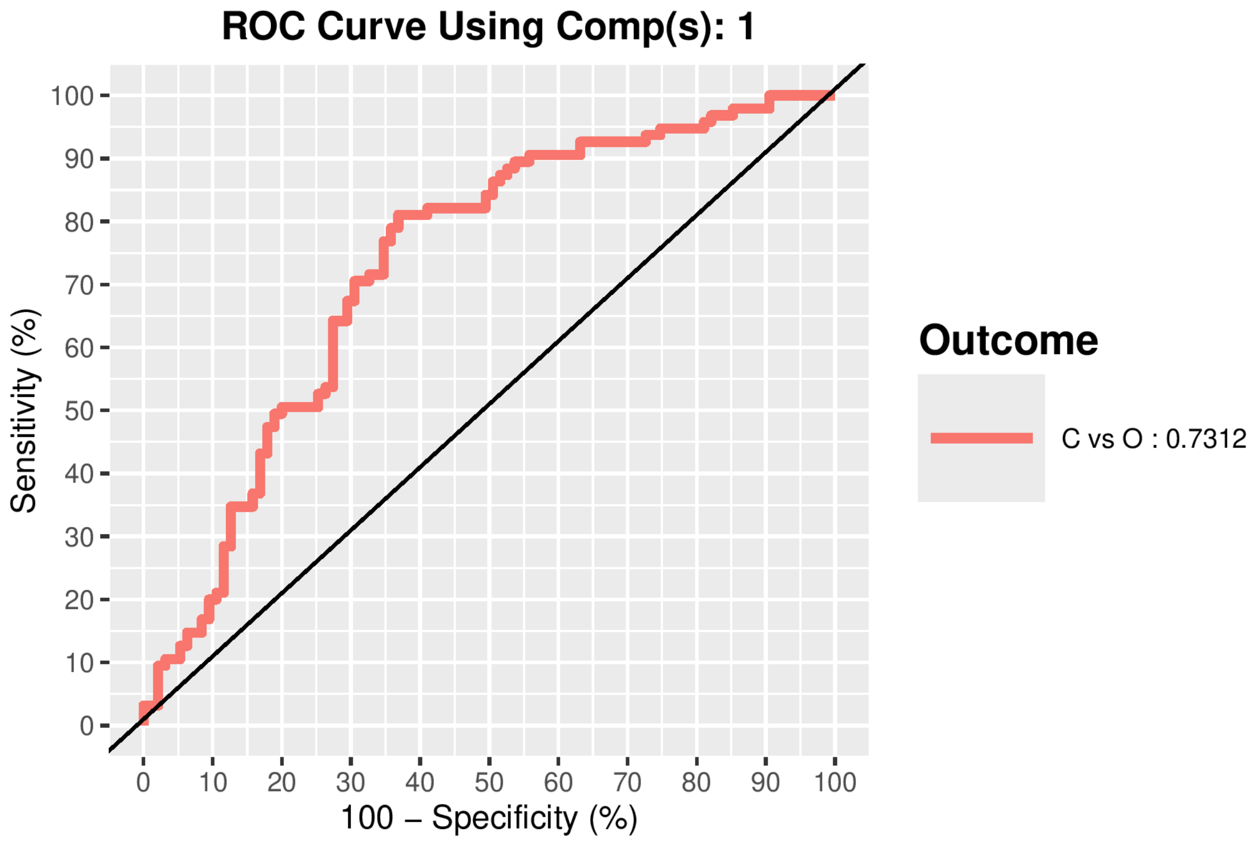


**Figure S13. ROC curve and AUC from the MINT sPLS-DA performed on the IMOS MGD MAGs (876 pMAGs$_{95\%ANI}$) for global component 1 for the fished vs. NTMRs reefs comparison**. Numerical outputs include the AUC (0.7312) and a Wilcoxon test p–value (p = $3.69 \times 10^{-8}$) for fished vs. NTMRs reefs class comparison that are performed per component.

### Zone-label shuffling validation for MINT sPLS-DA

We implemented zone-label shuffling to generate a null distribution against which observed model performance could be benchmarked. This approach tests whether the indicator taxa identified by MINT sPLS-DA reflect genuine biological associations with reef zoning rather than chance findings.

Specifically, reef protection status (fished reefs vs. No-Take Marine Reserves - NTMRs) labels were randomly shuffled within each GBR sector for a total of 999 permutations. This within-sector shuffling preserves the spatial structure of the data while breaking any association between microbial community composition and zoning status. All other aspects of the data, including microbial abundances, sample sizes per sector, and the total number of open and closed reefs, were held constant.

For each permuted dataset, we reran the full MINT sPLS-DA pipeline with identical parameters to the original model (ncomp = 2; keepX = c[350, 180], for component 1 and 2 respectively; and using centroids.dist, as determined optimal in model tuning), and validated classification accuracies using Leave-One-Group-Out Cross-Validation (LOGOCV) with sectors as groups. Classification accuracy was extracted from each permuted model using the mixOmics *perf()* function applied to the original model.

The p-value was calculated as p = (number of permutations with accuracy ≥ observed accuracy + 1) / (total permutations + 1). This formulation includes the observed value in the null distribution, providing a valid permutation test. Effect size was quantified using Cohen's *d* = (observed accuracy - mean(null accuracy)) / sd(null accuracy), which represents the difference between observed and null mean accuracies in units of standard deviation.

The null distribution comprised 999 successful permutations (100% success rate), with a mean accuracy of 49.9% (SD = 4.76%), closely centered on the theoretical random expectation of 50% for a model with only 2 categorical outcomes. The 95% confidence interval of the null distribution ranged from 40.4% to 59.2%. The observed classification accuracy (70.4%) was significantly higher than expected by chance ($p = 0.0010$), with none of the 999 permuted models exceeding the observed value (**Fig. S14; Table S6**). The effect size (Cohen's *d* = 4.32) indicates that model performance falls >4 standard deviations above the null mean, confirming that the identified indicator taxa reflect genuine biological differences between NTMRs and fished reefs rather than random chance (**Table S6**).

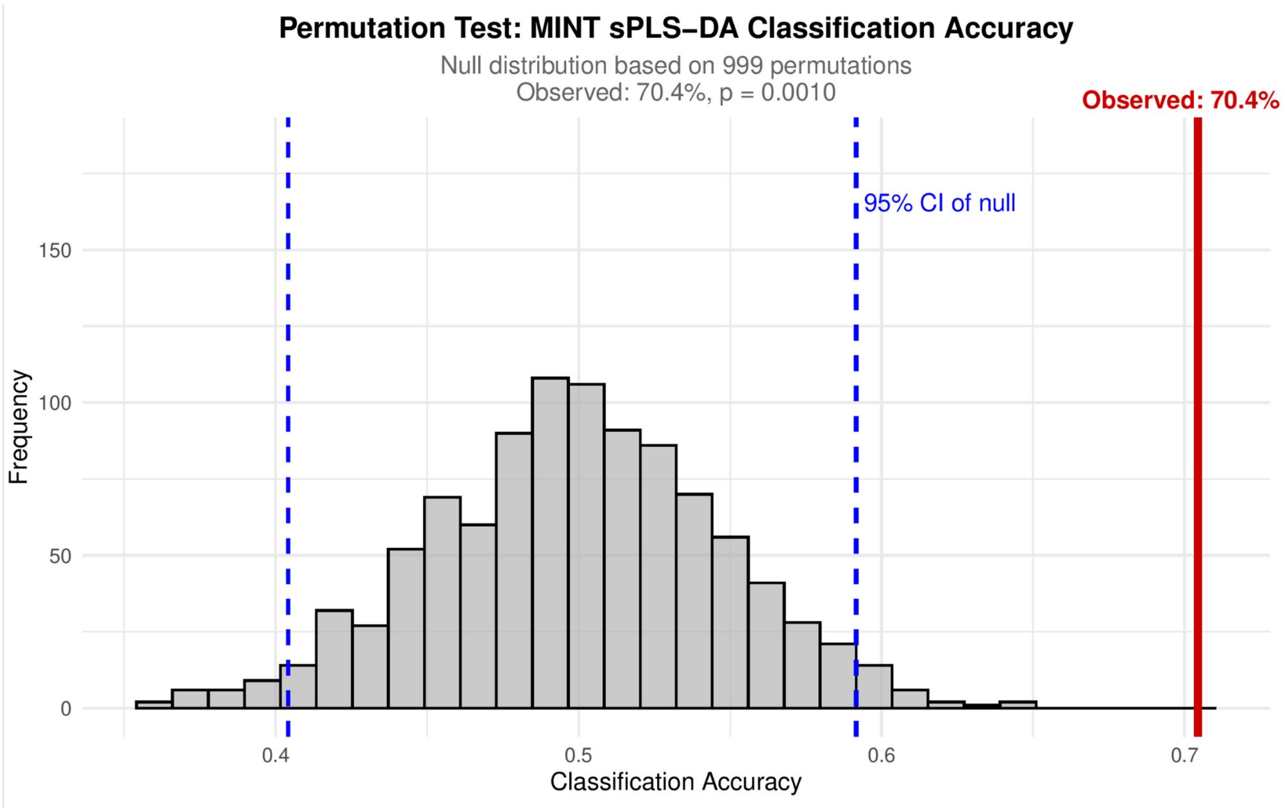


**Figure S14. Permutation test null distribution for MINT sPLS-DA classification accuracy**. Histogram showing the distribution of classification accuracies from 999 permutations where reef zoning status labels (fished vs. NTMR) were randomly shuffled within each GBR sector. For each permutation, MINT sPLS-DA was rerun with identical parameters (ncomp = 2, keepX = c(350, 180)) and evaluated using leave-one-group-out cross-validation (LOGOCV). Red vertical line: observed classification accuracy from the true model, i.e. before label shuffling (70.4%). The null distribution mean was 49.9%, with blue dashed lines representing 95% confidence interval of the null distribution [40.4%, 59.2%]. The null distribution represents the range of accuracies expected by chance when all association between microbial communities and zoning status is broken, while preserving the spatial structure of the data through within-sector label shuffling.

**Table S6. Summary statistics from zone-label permutation testing**. Results from 999 permutations where reef zoning status labels were randomly shuffled within each GBR sector to generate a null distribution of classification accuracies under the hypothesis of no association between microbial communities and protection status. All MINT sPLS-DA parameters were held constant across permutations (ncomp = 2, keepX = c(350, 180)), with classification accuracy evaluated via leave-one-group-out cross-validation.

| Metric | Value |
|---|---|
| Observed accuracy | 70.44% |
| Null distribution mean (±SD) | 49.90% (±4.76%) |
| 95% CI of null | [40.41%, 59.16%] |
| p-value | 0.0010 |
| Effect size (Cohen's d) | 4.32 |

### Random Forest Models

To provide an independent, non-linear supervised framework alongside MINT sPLS-DA and validate the robustness of microbial indicators of reef zoning, we implemented Random Forest (RF) classification[13]. This approach complements MINT sPLS-DA by offering a fundamentally different modelling paradigm, decision trees that capture non-linear relationships and complex interactions among predictors, while maintaining the same cross-validation structure for direct comparability. RF models were trained using CLR-transformed abundance data (190 samples × 876 pMAGs) with reef protection status (Open_or_Closed_to_fishing; 91 NTMR samples, 99 fished samples) as the response variable, and the GBR sector selected for the stratified cross-validation, identical to the MINT framework.

To ensure direct comparability with MINT sPLS-DA, we employed leave-one-sector-out cross-validation, iteratively training on six GBR sectors and predicting on the seventh, thus testing whether microbial communities can predict zoning status in entirely unseen geographic regions with distinct environmental conditions. For each training fold, Random Forest models were constructed using the randomForest R package[14] (v4.7-1.2) with the following parameters: (ntree = 500); square root of the total number of predictors (mtry = floor(sqrt(p)), where p = 876 pMAGs); stratified bootstrap sampling with equal representation of fished and NTMR reefs in each training iteration (sampsize = rep(min(table(Y_train)), 2)) for class imbalance handling; and feature importance recorded for each fold using importance = TRUE. Two importance metrics were computed: (1) Mean Decrease Accuracy, which represents the reduction in prediction accuracy when the values of a given pMAG are randomly permuted, and (2) Mean Decrease Gini: the reduction in node impurity attributable to each pMAG (higher values indicate that a variable is more crucial for separating classes).

Following the same approach applied to MINT sPLS-DA, we implemented permutation testing to assess whether the observed classification accuracy exceeded expectations under the null hypothesis of no association between microbial communities and zoning status. For 999 permutations, reef zoning labels were randomly shuffled within each sector, preserving the spatial structure of the data while breaking any association between microbial communities and protection status. Classification accuracy was recorded for each permutation, generating a null distribution of accuracies expected by chance, and the permutation p-value was calculated as: p = (number of permutations with accuracy ≥ observed accuracy + 1) / 999 + 1. This formulation includes the observed value in the null distribution, providing a valid permutation test[15]. The 95% confidence interval of the null distribution was derived from the 2.5th and 97.5th percentiles of permuted accuracies.

To identify the most influential pMAGs for zoning classification, we averaged importance scores (Mean

Decrease Accuracy) across all seven cross-validation folds to account for sector-specific variation and provide stable, generalizable importance estimates. pMAGs were ranked by descending Mean Decrease Accuracy. To enable direct comparison with MINT sPLS-DA (which selected 350 indicator MAGs on component 1), we extracted the top 350 most important features based on their importance scores. Methodological concordance between RF and MINT sPLS-DA was assessed through: (1) overlap analysis, by calculating the intersection (i.e. overlap percentage) between the 350 MINT indicators (component 1) and the top 350 RF-important features; and via (2) the Jaccard similarity index: a standardised measure of set similarity ranging from 0 (no overlap) to 1 (perfect overlap).

For the overlapping pMAGs identified by both methods, we assessed whether they showed consistent enrichment direction (NTMR-enriched vs. Fished-enriched). Enrichment direction was determined as follows: (1) for MINT sPLS-DA, direction was derived from loading values on component 1 (Positive loadings = NTMR-enriched; negative loadings = Fished-enriched), and (2) for Random Forest, mean CLR abundance was calculated for each MAG in NTMR and fished reefs, and the difference (mean_NTMR_CLR - mean_Fished_CLR) was computed (Positive difference = NTMR-enriched; negative difference = Fished-enriched). Direction agreement was quantified using: (1) percentage agreement: proportion of overlapping MAGs with identical direction assignments; and (2) Cohen's Kappa, which is a chance-corrected measure of inter-method agreement[16], calculated using the *kappa2()* function from the irr (v0.84.1) R package[17]. Cohen's Kappa interpretation follows standard guidelines[18]: < 0.20 = slight agreement; 0.21-0.40 = fair agreement; 0.41-0.60 = moderate agreement; 0.61-0.80 = substantial agreement; 0.81-1.00 = almost perfect agreement. Taxonomy of overlapping indicators was visualised at family level, separated by enrichment direction. The top 15 families by total MAG count were displayed to highlight the dominant taxonomic groups associated with each reef zone.

Parallel processing was implemented using doParallel[19] (v1.0.17) (Microsoft Corporation and Weston 2022) and foreach (v1.5.2) (Microsoft Corporation and Weston 2022) to expedite the computationally intensive permutation testing (999 iterations × 7 CV folds), utilizing N-5 cores to leave resources for system operations. All random processes were initialized with set.seed(123) to ensure reproducibility.

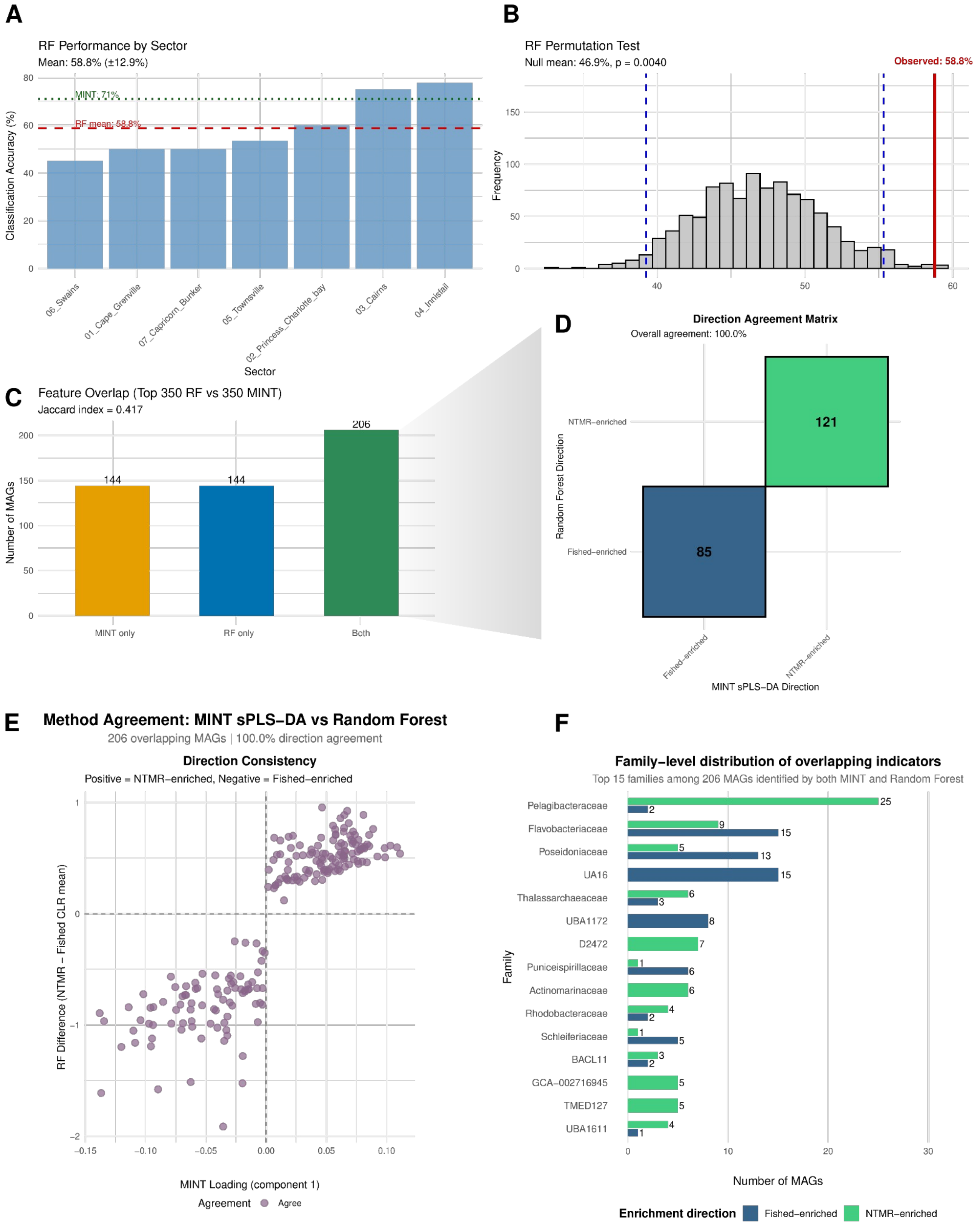


**Figure S15. Random Forest validation of reef zoning classification**. (a) Leave-one-sector-out cross-validation accuracy across the seven GBR sectors. The red dashed line indicates mean RF accuracy (58.8%) averaged across the 7 GBR sectors, the green dotted line shows MINT sPLS-DA accuracy (71%) for comparison. (b) Permutation test results showing the null distribution of classification accuracies from 999 permutations with zone labels shuffled within sectors. The red vertical line indicates observed RF accuracy (58.8%), blue dashed lines show the 95% confidence interval of the null distribution [39.3%, 55.3%]. p = 0.004, confirming significant predictive performance above chance. (c) Overlap between MINT sPLS-DA indicators (n = 350) and the top 350 most important RF features. Of the 350 MINT indicators, 206 (58.9%) appear in the RF top 350, representing substantial methodological concordance (Jaccard index = 0.417). (d) Confusion matrix showing perfect agreement in enrichment direction assignment between methods. From the 206 overlapping indicators between MINT sPLS-DA and RF models, all 121 NTMR-enriched and 85 Fished-enriched MAGs identified by both approaches show identical direction (Cohen's Kappa = 1.000, $p < 0.001$), confirming that the core microbial signal is robust across fundamentally different modelling frameworks. (e) Scatter plot of MINT loadings vs. RF difference (NTMR - Fished CLR mean) for the 206 overlapping MAGs. Points in the upper-right and lower-left quadrants indicate perfect directional concordance (positive = NTMR-enriched, negative = Fished-enriched). (f) Family-level taxonomic distribution of the 206 overlapping pMAGs, separated by enrichment direction. NTMR-enriched MAGs (green) are dominated by oligotrophic taxa including Pelagibacteraceae, Marinisomatota, and SAR86, while Fished-enriched MAGs (blue) are dominated by copiotrophic taxa including Flavobacteriaceae, UA16, and Schleiferiaceae, consistent with the ecological patterns observed in the full MINT analysis.

### ANOVA-like Differential EXpression - ALDEx2

To complement our primary MINT sPLS-DA analysis, we performed per-MAG differential abundance testing using ALDEx2[20] (v1.34.0) with two approaches: (1) a simple comparison without covariates, and (2) a generalized linear model (GLM) including sampling trip and GBR sector as fixed effects to control for spatiotemporal variation. All analyses used 128 Monte Carlo samples and FDR < 0.05 significance threshold, applied to the 876 pMAGs after CLR transformation.

#### Simple ALDEx2

We first performed a simple (i.e. without accounting for spatiotemporal covariates) ALDEx2 analysis comparing pMAG abundances between 91 NTMR (Closed) and 99 fished (Open) reef samples. The analysis was conducted with 128 Monte Carlo samples using 'all' as the denominator for CLR transformation. We computed both Welch's t-test and Wilcoxon rank-sum test statistics separately, then calculated effect sizes with confidence intervals. Significance was assessed using Benjamini-Hochberg FDR correction, and we performed comprehensive validation including test consistency checks, effect size categorization, and genus-level taxonomic analysis of significant MAGs.

The simple ALDEx2 analysis identified 304 significantly differentially abundant pMAGs at FDR < 0.05 (Welch's test) (**Fig. S16A**), with 196 enriched in NTMRs and 108 in fished reefs. The distribution of raw p-values showed deviation from the null expectation (**Fig. S16B**), confirming enrichment of true positive signals. Effect sizes ranged from -0.59 to 0.64 (mean = -0.034), with the majority (862/876 pMAGs) showing very small effect sizes (<0.5) (**Fig. S16C**). Overlap statistics indicated high effect certainty, with most pMAGs showing low overlap values suggesting reproducible differential abundance patterns across Monte Carlo samples (**Fig. S16D**).

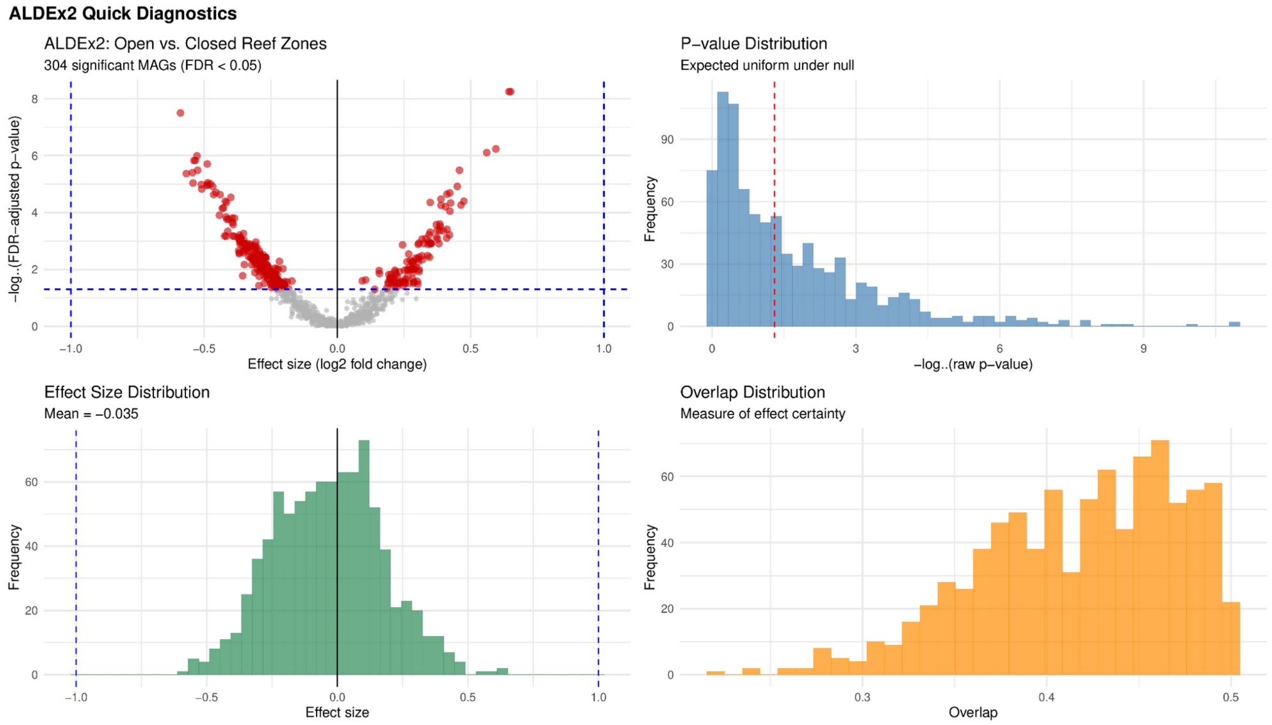


**Figure S16. ALDEx2 analysis (simple model) comparing pMAG abundances between No-Take Marine Reserves (NTMRs) and fished reefs (n = 876 MAGs)**. (**a**) Volcano plot showing effect sizes ($log_2$ fold change) against FDR-adjusted p-values. Red points indicate significant pMAGs (FDR < 0.05), with positive effects enriched in fished reefs and negative effects in NTMRs. (**b**) Distribution of raw p-values (Welch's t-test). (**c**) Distribution of effect sizes. (**d**) Distribution of overlap statistics, measuring effect certainty.

Furthermore, Welch's and Wilcoxon tests showed high agreement (282/304 MAGs concordant; Spearman $\rho = 0.954$, $p < 0.001$) (**Fig. S17**), despite effect sizes being predominantly small (862 MAGs < 0.5).

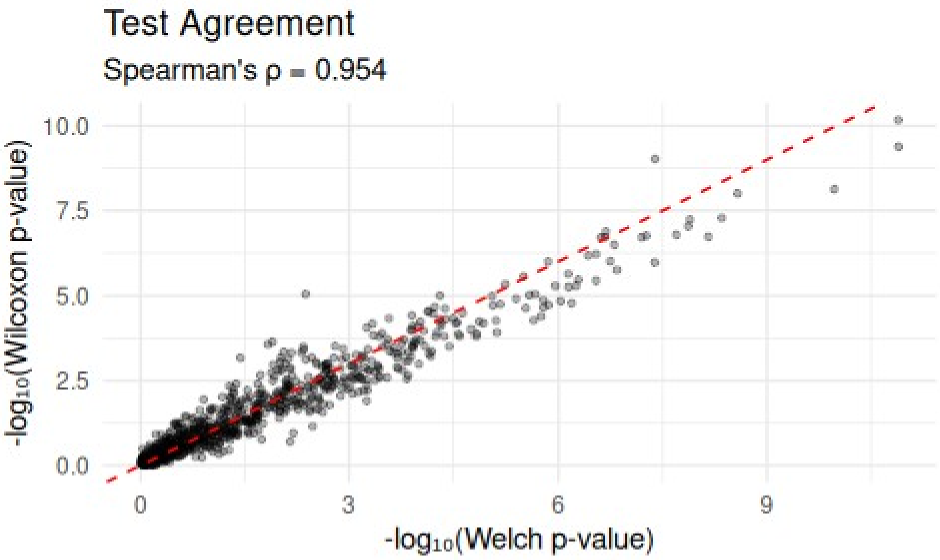


**Figure S17. Consistency between ALDEx2 statistical tests for differential abundance detection**. Scatter plot comparing $-\log_{10}$ transformed p-values from Welch's t-test (x-axis) and Wilcoxon rank-sum test (y-axis) for detecting differentially abundant pMAGs between reef protection zones. The red dashed line indicates perfect agreement (y = x). 282 of 304 significant MAGs (FDR < 0.05) were identified by both tests, supporting robust identification of zoning-associated taxa.

Lastly, taxonomic analysis revealed distinct enrichment patterns: *Pelagibacter* dominated NTMR-enriched MAGs (46 MAGs), while Poseidonia (11 pMAGs) and UBA11663 (10 pMAGs) were prominent in fished reefs (**Fig. S18A**). The analysis also identified more indicators of NTMRs vs fished reefs (**Fig S18B**), capturing 47 unique genera enriched in fished reefs and 80 in NTMRs, with 22 genera showing mixed patterns (**Fig S18C**), demonstrating specific microbial signatures associated with reef protection status.

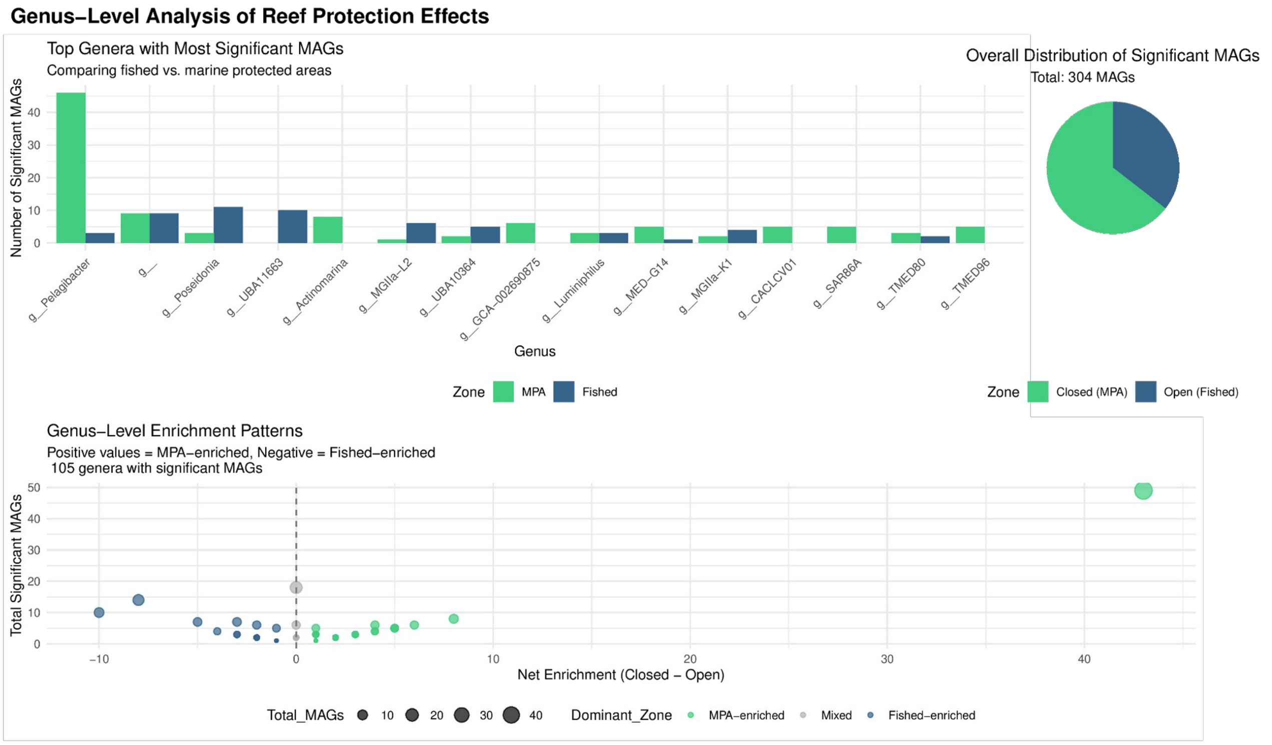


**Figure S18. Genus-level patterns of differentially abundant pMAGs between reef protection zones**. (**A**) Bar chart showing the number of significant pMAGs belonging to the 15 most abundant genera, separated by enrichment in fished (blue) versus No-Take Marine Reserve (NTMR; green) zones. (**B**) Pie chart showing the overall distribution of significant indicator pMAGs between zones. (**C**) Bubble plot displaying genus-level enrichment patterns, where point size represents total significant pMAGs per genus, position on the x-axis shows net enrichment (positive = NTMR-enriched, negative = fished-enriched), and color indicates the dominant zone of enrichment.

### Complex ALDEx2 GLM model with spatiotemporal covariates

The complex (i.e., controlling for spatiotemporal variation) ALDEx2 GLM model was implemented using a design matrix with reef protection status as the primary predictor, while explicitly including sampling trip and GBR sector as fixed effects. The comparison results with the simple ALDEx2 results strongly confirm the robustness of our core findings. The GLM model accounting for covariates identified 316 differentially abundant MAGs (FDR < 0.05), with 214 enriched in No-Take Marine Reserves (NTMRs) and 102 enriched in fished reefs. Critically, 237 of the 304 MAGs identified as significant in the simple model (77.7%) remained significant in the covariate-adjusted GLM (**Fig. S19A**), showing 100% agreement in the direction of enrichment (e.g., *Pelagibacter* enriched in NTMRs, *Poseidonia* and UBA11663 enriched in fished reefs). The effect sizes between models were highly correlated (Spearman $\rho = 0.882$, $p < 0.001$), indicating stable signals (**Fig. S19B**) and similar statistical power between models (**Fig. S19C**).

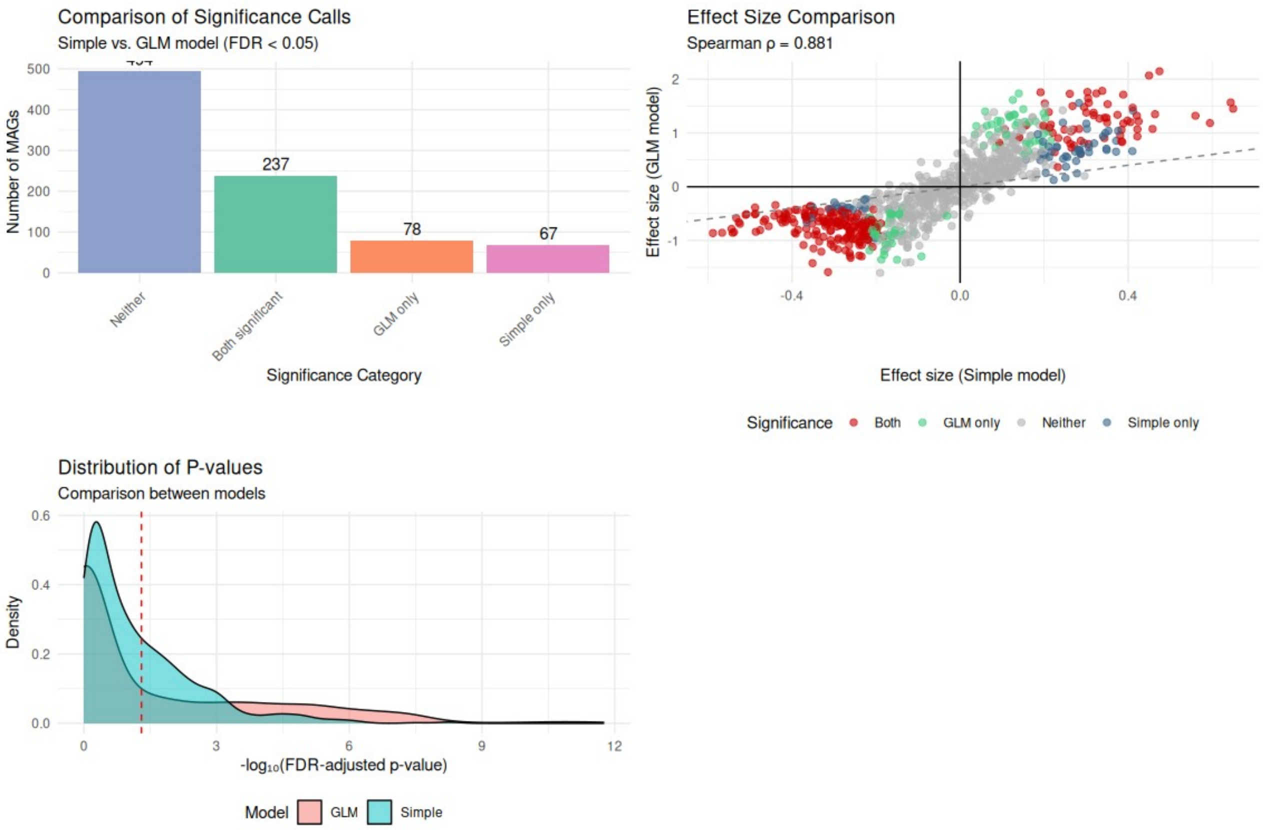


**Figure S19. Comparison of simple versus covariate-adjusted ALDEx2 models for detecting reef protection indicators**. (**A**) Agreement in significance calls between ALDEx2 models with and without spatiotemporal covariates. (**B**) Correlation of effect sizes between simple and GLM models. (**C**) Distribution of FDR-adjusted p-values across both models.

### Comparison between ALDEx2 + GLM and MINT sPLS-DA selected indicators

To validate MINT sPLS-DA-selected indicators using orthogonal statistical testing, we compared the 350 MAGs identified by MINT sPLS-DA (component 1 loadings) with those showing significant differential abundance in the ALDEx2 GLM (FDR $< 0.05$), examining both overlap magnitude and enrichment direction consistency. Enrichment direction was determined from MINT loadings (negative = fished-enriched, positive = NTMR-enriched) and ALDEx2 effect sizes (positive = fished-enriched, negative = NTMR-enriched).

The comparison revealed exceptional concordance between methods (**Fig. S20**). Of the 350 MAGs selected by MINT sPLS-DA as reef protection indicators, 293 (83.7%) were independently validated as significantly differentially abundant by ALDEx2 GLM models that accounted for spatiotemporal covariates (**Fig. S20A**). The overlapping indicators represented diverse taxonomic groups, with the top 12 families including Pelagibacteraceae (NTMR-enriched), Flavobacteriaceae (mixed), and Rhodobacteraceae (fished-enriched) (**Fig. S20B**). pMAGs significant in both methods showed stronger effect sizes than those identified by ALDEx2 alone, indicating that multivariate feature selection with MINT sPLS-DA preferentially captured pMAGs with more pronounced reef protection signals (**Fig. S20C**). Lastly, among these overlapping indicators, 100% showed consistent enrichment direction between methods, with pMAGs identified as NTMR-enriched by MINT also found to be NTMR-enriched by ALDEx2, identical for fished-reef enriched pMAGs showing agreement between MINT and ALDEx2 (**Fig. S20D**).

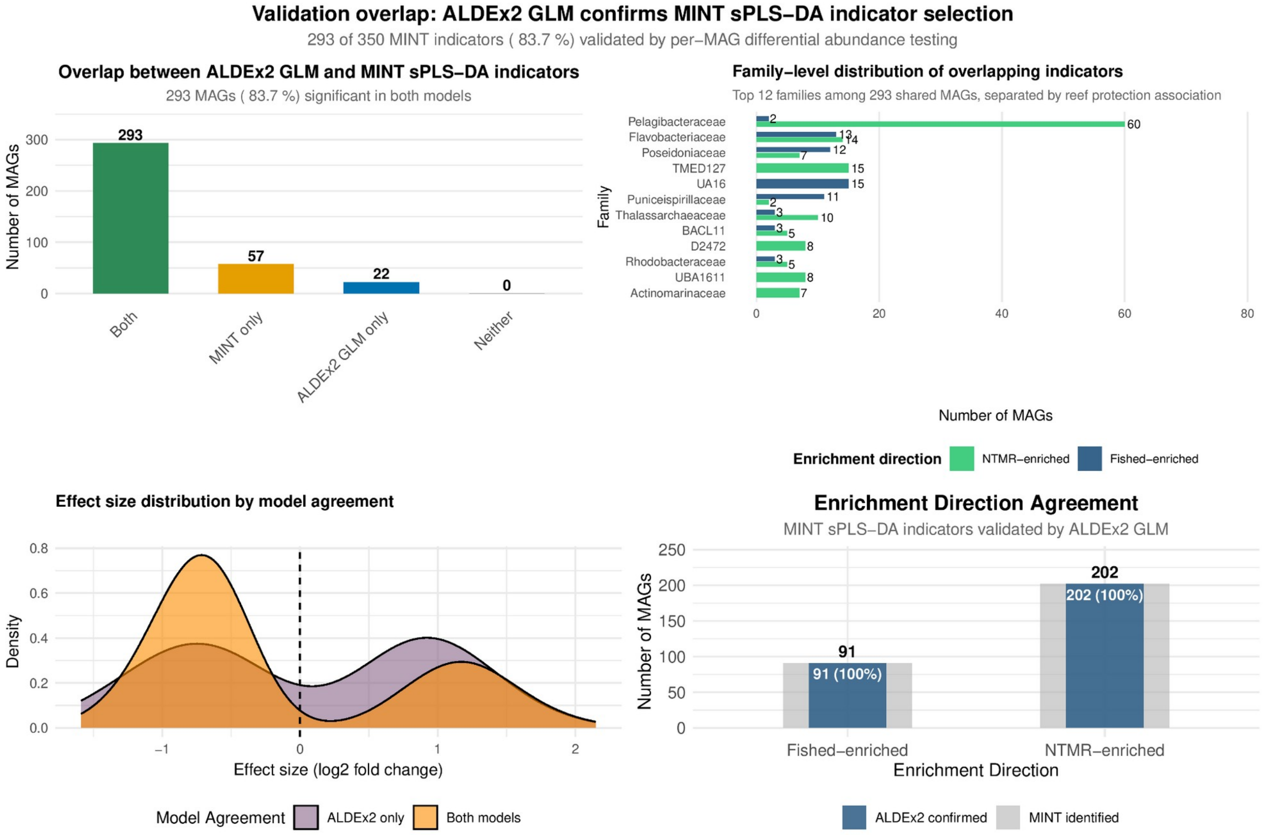


**Figure S20. Methodological concordance between multivariate feature selection (MINT sPLS-DA) and per-MAG statistical testing (ALDEx2) for identifying reef protection indicators**. (**A**) Overlap analysis comparing pMAGs selected by MINT sPLS-DA (component 1) with those showing significant differential abundance in ALDEx2 GLM models. (**B**) Taxonomic distribution of overlapping MAGs showing the top 12 families, separated by enrichment direction (NTMR-enriched vs. fished-reef enriched). (**C**) Density distributions of effect sizes for pMAGs identified by both methods (orange) versus ALDEx2 only (purple). (**D**) Comparison of enrichment direction agreement between methods for overlapping indicators.

### Ruling out genome size bias

#### Presence/absence analysis

We analysed the presence/absence patterns of the 350 MINT sPLS-DA indicator pMAGs to evaluate whether observed abundance differences might be driven by genome size bias rather than ecological patterns. Our analysis reveals near-universal presence across all sites: of the 350 indicator pMAGs, 317 (90.6%) were detected in all 190 seawater samples across the Great Barrier Reef, with the remaining 33 pMAGs present in 91-99% of samples (**Fig. S21**). This demonstrates these are not rare taxa with detection-limited signals, but widespread members of the GBR seawater microbiome. This universal presence indicates that our MINT sPLS-DA results reflect genuine ecological variation in microbial abundance values, not detection artifacts related to genome size (i.e. genome size remains consistent across sites) or (under)sampling.

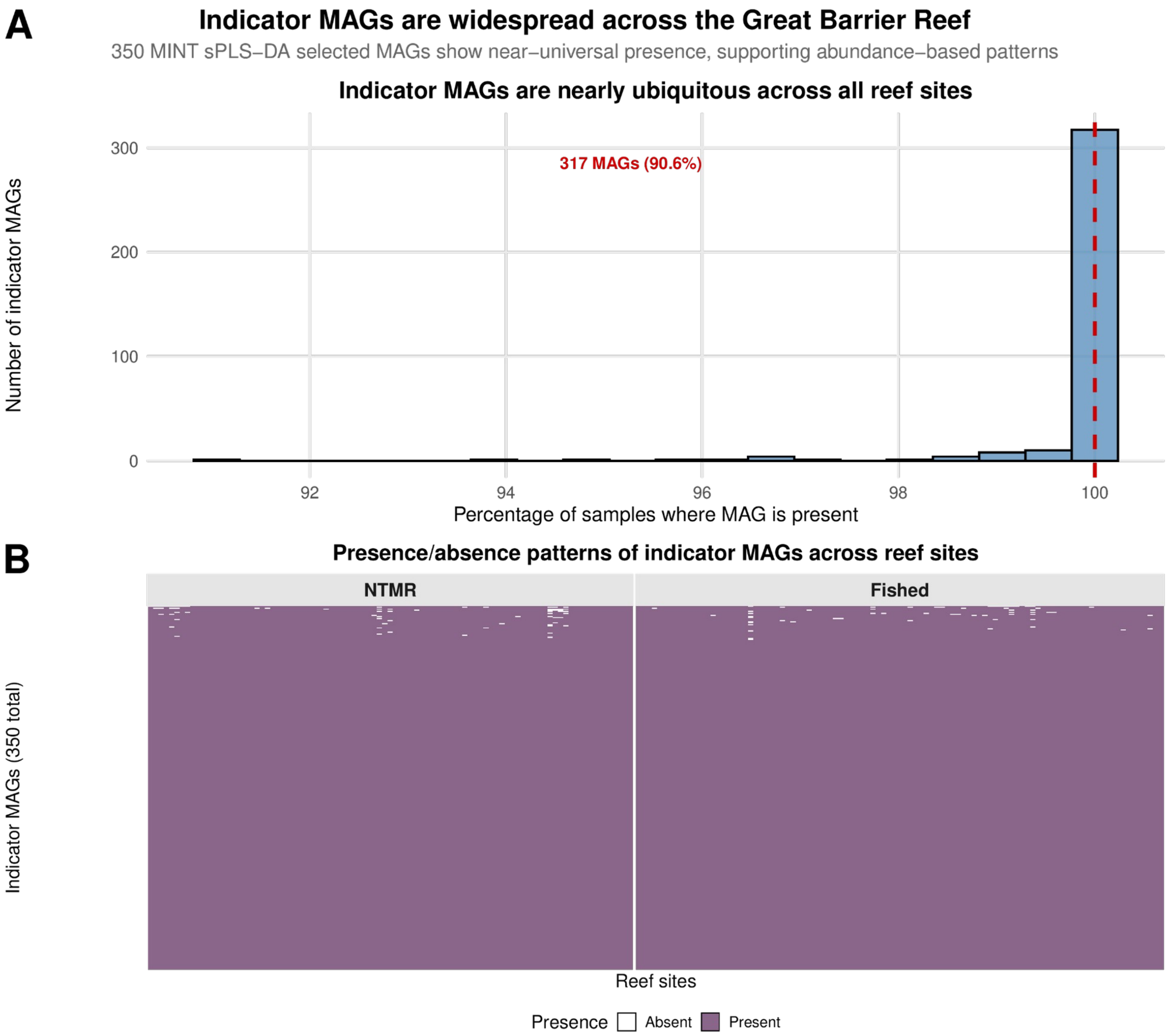


**Figure S21. Near-universal presence of MINT sPLS-DA indicator pMAGs across the Great Barrier Reef validates abundance-based ecological signals**. (**A**) Distribution of presence percentages for the 350 MINT sPLS-DA-selected indicator pMAGs across 190 seawater samples. The dashed red line indicates complete presence (100% of samples), with 317 pMAGs (90.6%) detected in every sample. (**B**) Presence/absence heatmap showing all 350 indicator pMAGs (y-axis, ordered by presence frequency) across all reef sites (x-axis), separated into NTMR (closed) and fished (open) zones. White tiles indicate absence, purple tiles indicate presence.

### Read-based metagenomic validation independent of pMAGs

To validate our MAG-based findings using an orthogonal approach that is free from biases introduced by MAG binning, completeness, or genome size estimation, we performed MINT sPLS-DA on the read-based taxonomic dataset (621 prokaryotic taxa) generated from the same metagenomic reads, previously published in Terzin et al. (2025)[21]. Briefly, quality-filtered reads were aligned against the NCBI nr database using DIAMOND[22] (v2.0.9) with an e-value threshold of $< 1 \times 10^{-5}$. Resulting alignments were imported into MEGAN[23] (v6.23.0), where the Lowest Common Ancestor (LCA) algorithm was applied to assign each read to the most specific taxonomic node shared by all reference sequences. This conservative approach avoids over-classification when reads match multiple organisms. Taxonomic abundance counts were exported at the genus level and imported into R using the phyloseq[24] package. Pre-filtering steps included removal of: (1) non-annotated reads; (2) reads annotated as eukaryotic or viral; (3) prokaryotic reads annotated only to the domain level (Bacteria or Archaea); and (4) rare/spurious reads with relative abundance < 0.0001%. This filtering resulted in a final dataset of 621 prokaryotic taxa (collapsed at genus level or above). Microbial abundance data were then center log-ratio (CLR) transformed using the microbiome[25] R package to account for sparsity and compositional nature of metagenomic sequencing data, with pseudocounts introduced prior to CLR transformation as log 0 is undefined.

MINT sPLS-DA[9,12,26] was applied with reef zoning status as the categorical outcome and GBR sector as the grouping factor to account for spatial structure. Model tuning identified one component as optimal, with 10 features selected per component. Sample plots revealed separation between NTMRs and fished reefs along component 1, with no residual clustering by sampling trip, confirming that spatiotemporal batch effects were effectively accounted for (**Fig. S22A**). The Clustered Image Map (CIM) for components 1 and 2 showed distinct enrichment patterns, with a subset of taxa enriched in NTMRs and a separate subset enriched in fished reefs. Taxa enriched in NTMRs were dominated by oligotrophs like Pelagibacteraceae (SAR11) and *Prochlorococcus*, while taxa enriched in fished reefs included members of Flavobacteriaceae and Rhodobacteraceae (**Fig. S22B**). These taxonomic patterns are largely consistent with those identified in our primary MAG-based MINT sPLS-DA analysis, though classification accuracy was higher with MAGs (71% vs. 66%), confirming the robustness of our findings across independent analytical frameworks.

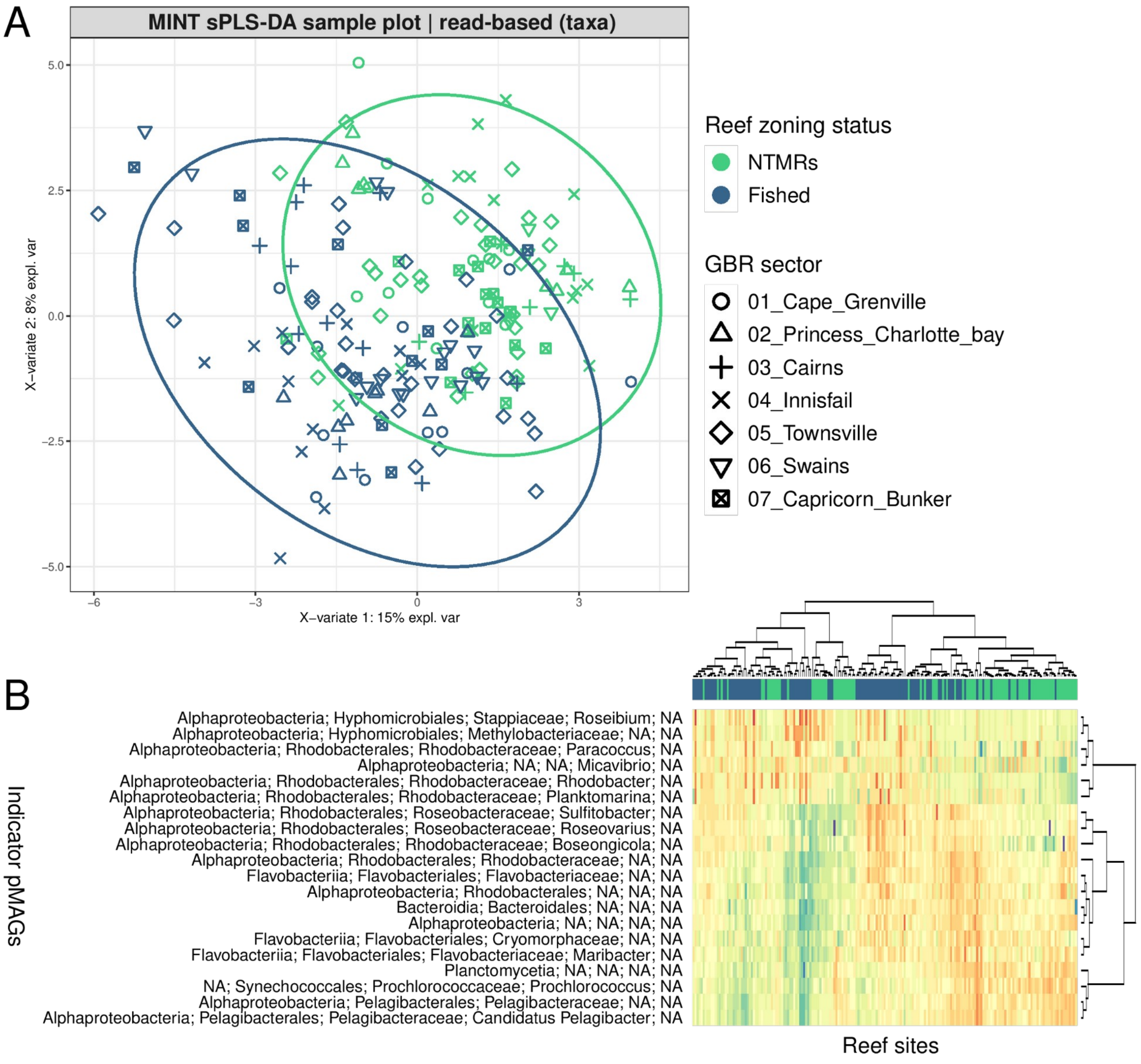


**Figure S22. Community-wide taxonomic validation of reef zoning signals using read-based analysis (621 microbial taxonomic features collapsed at genus level or above derived from DIAMOND → MEGAN read-centric profiling)**. (A) Global MINT sPLS-DA sample plot showing separation of NTMRs (green) and fished reefs (blue), with 95% confidence ellipses. (B) Clustered Image Map (CIM) of microbial taxa selected by MINT sPLS-DA on components 1–2, with reef sites (rows) coloured by protection status. This figure complements patterns identified in the MAG-based analysis (**Fig. 2; main text**).

### **M**ultivariate **INT**egration (**MINT**) **sPLS** | Correlating microbial and environmental data (**sPLS**), while accounting for sector-specific effects (**MINT**)

**Table S7. Mean ± standard deviation (SD) of MINT sPLS partial correlation scores between microbial indicators of zoning and environmental variables**. This table complements the heatmap in the main text (**Fig 3B; main text**), with positive values (red) indicating positive associations, and negative values (blue) indicate negative associations. NTMRs = No-Take Marine Reserves.

| **Reef Variable** | **Fished Reef Indicators (n = 114)** | **NTMR Indicators (n = 236)** |
|---|---|---|
| POC_µM | 0.08 ± 0.13 | -0.12 ± 0.17 |
| SEAWATER_TEMPERATURE_2.5m_RV | 0.06 ± 0.13 | -0.1 ± 0.16 |
| Foliose_non_Acropora | 0.08 ± 0.1 | -0.12 ± 0.13 |
| Carnivore | 0.01 ± 0.11 | -0.04 ± 0.14 |
| TDN_µM | 0.01 ± 0.11 | -0.03 ± 0.13 |
| Sand | 0.01 ± 0.13 | -0.03 ± 0.15 |
| Turf_algae | 0.17 ± 0.12 | -0.24 ± 0.15 |
| PP_µM | 0.14 ± 0.11 | -0.2 ± 0.14 |
| SALINITY_2.5m_RV | 0.2 ± 0.13 | -0.28 ± 0.17 |
| Phaeophytin_A_µg_L | 0.16 ± 0.08 | -0.22 ± 0.1 |

| | | |
|---|---|---|
| Massive_non_Acropora | -0.11 ± 0.09 | 0.12 ± 0.1 |
| Piscivore | -0.11 ± 0.07 | 0.13 ± 0.07 |
| FLUORESCENCE_2.5m_RV | -0.06 ± 0.07 | 0.06 ± 0.08 |
| NO2_µM | 0.14 ± 0.06 | -0.18 ± 0.05 |
| Lobate_Soft_Coral | 0.02 ± 0.07 | -0.01 ± 0.08 |
| PO4_µM | -0.01 ± 0.11 | 0.04 ± 0.13 |
| Coralline_algae | -0.14 ± 0.17 | 0.22 ± 0.22 |
| Tabulate_Acropora | -0.13 ± 0.14 | 0.19 ± 0.18 |
| Digitate_Acropora | -0.12 ± 0.16 | 0.18 ± 0.2 |
| Herbivore | -0.15 ± 0.08 | 0.21 ± 0.09 |
| Detritivore | -0.15 ± 0.12 | 0.21 ± 0.15 |
| TDP_µM | -0.17 ± 0.14 | 0.24 ± 0.17 |
| Submassive_non_Acropora | -0.24 ± 0.13 | 0.33 ± 0.16 |
| Encrusting_Acropora | -0.21 ± 0.1 | 0.28 ± 0.12 |
| Millepora | -0.2 ± 0.11 | 0.27 ± 0.14 |

### Generalised Mixed Models (GLMMs) - do NTMRs and fished reefs differ in benthic cover (hard coral, algae), fish biomass, and densities of herbivorous fish?

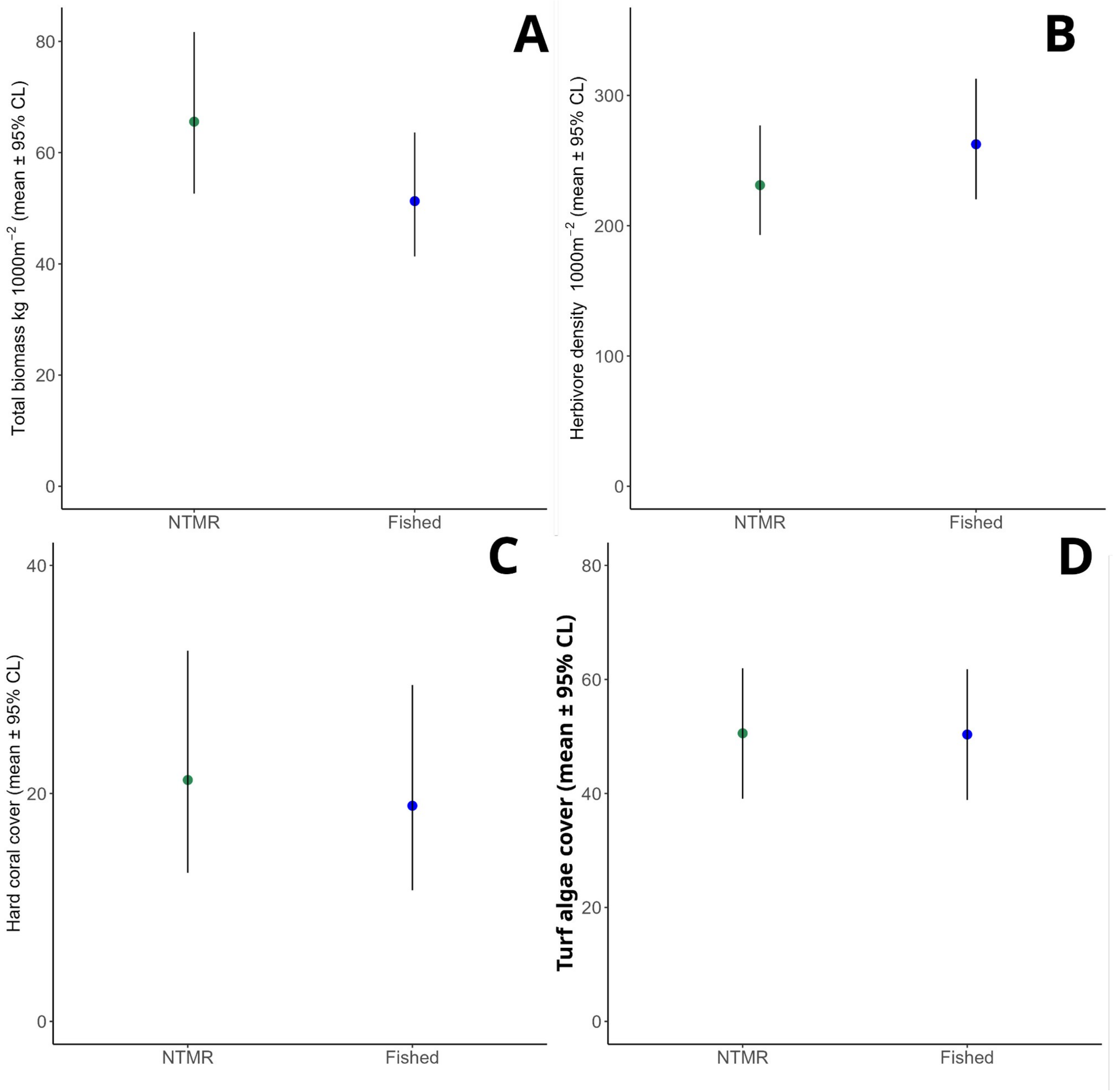


**Figure S23. Comparison of (A) total fish biomass, (B) herbivore density, (C) hard coral, and (D) turf algae cover between No-Take Marine Reserves (NTMRs) and fished reef zones, using Generalised Mixed Models (GLMMs)**. Bars show estimated marginal means (± SE) from a Gamma GLMM, with closed zones (NTMRs) supporting 24.6% higher biomass than open zones (O) ($p = 0.014$). Differences in other metrics (B-D) were insignificant. Random effects accounted for spatial hierarchy (sector, shelf, reef/site/transect); error distributions used Gamma-log link.

### Total fish biomass

#### Raw data

Table S8. Summary statistics (descriptive) for overall fish biomass.

| Status | Mean (g) | Median (g) | SD | n | SE |
|---|---|---|---|---|---|
| Closed | 100,915 | 61,616 | 129,789 | 345 | 6,988 |
| Open | 71,652 | 51,186 | 116,017 | 360 | 6,115 |

#### GLMM Results (Gamma Distribution)

```
### Running the model

biom.tmb <- glmmTMB(

biomass ~ OPENORCLOSED_AFTER2004+(1|A_SECTOR)+(1|SHELF)+(1|REEF_NAME/SITE_NO/
TRANSECT_NO),

data = tot.biom,

family = Gamma(link = "log")

)

summary(biom.tmb)
```

Model Specification

**Family**: Gamma (log link)
**Formula**:
biomass ~ OPENORCLOSED_AFTER2004 + (1 | A_SECTOR) + (1 | SHELF) + (1 | REEF_NAME/SITE_NO/TRANSECT_NO)
**Dataset**: total fish biomass
**Observations**: 705

Model Fit Statistics

(Note: AIC/BIC not available)

Table S9. Random Effects Variance.

| Group | Variance | Std. Deviation |
|---|---|---|
| A_SECTOR | 0.0497 | 0.223 |
| SHELF | $2.07 \times 10^{-7}$ | ~0 |
| TRANSECT_NO:SITE_NO:REEF_NAME | 0.324 | 0.569 |
| SITE_NO:REEF_NAME | 0.154 | 0.392 |
| REEF_NAME | 0.0412 | 0.203 |

Groups:

- A_SECTOR: 7 levels
- SHELF: 2 levels
- REEF_NAME: 47 levels

**Dispersion parameter ($\sigma^2$)**: $3.64 \times 10^{-8}$

Table S10. Fixed Effects

| Term | Estimate | Std. Error | z-value | p-value |
|---|---|---|---|---|
| (Intercept) | 11.091 | 0.112 | 98.87 | < 0.001 *** |
| OPENORCLOSED_AFTER2004O | -0.246 | 0.100 | -2.47 | 0.014 * |

#### Key Interpretation

- **Significant effect** of protection status:
  - Open (fished) areas had **24.6% lower biomass** (95% CI: 4.4%-44.8%) than NTMRs (*z* = -2.47, *p* = 0.014).
- Most variance explained by:
  - Transect-level effects (56.9% SD).
  - Site-level effects (39.2% SD).

### Herbivorous Fish Density

#### Raw data

Table S11. Summary statistics (descriptive) for herbivorous fish densities.

| Protection Status | Mean density | Median density | SD | n | SE |
|---|---|---|---|---|---|
| Closed (NTMR) | 259 | 244 | 141 | 345 | 7.60 |

| Open (Fished) | 294 | 266 | 157 | 360 | 8.28 |
|---|---|---|---|---|---|

#### GLMM Results (Negative Binomial)

```
### Run a glmm using library(glmm.tmb)

herb.tmb <- glmmTMB(

Density ~ OPENORCLOSED_AFTER2004+(1|A_SECTOR)+(1|SHELF)+(1|REEF_NAME/SITE_NO/
TRANSECT_NO),

data = herbs,

family = poisson(link = "log")

)

### Looking at the results now:

summary(herb.tmb)
```

Model Specification
Family: Negative Binomial (log link)
Formula:
Density ~ OPENORCLOSED_AFTER2004 + (1 | A_SECTOR) + (1 | SHELF) + (1 | REEF_NAME/SITE_NO/TRANSECT_NO)
Dataset: herbivorous fish (density)
Observations: 705

Table S12. Model Fit Statistics

| Statistic | Value |
|---|---|
| AIC | 8659.4 |
| BIC | 8695.9 |
| Log-Likelihood | -4321.7 |
| Deviance | 8643.4 |

Table S13. Random Effects Variance

| Group | Variance | Std. Deviation |
|---|---|---|
| A_SECTOR | $2.125 \times 10^{-2}$ | 0.146 |
| SHELF | $3.510 \times 10^{-14}$ | ~0 |
| TRANSECT_NO:SITE_NO:REEF_NAME | $3.192 \times 10^{-9}$ | ~0 |
| SITE_NO:REEF_NAME | $7.097 \times 10^{-2}$ | 0.266 |
| REEF_NAME | $7.796 \times 10^{-2}$ | 0.279 |

**Groups**:

- A_SECTOR: 7 levels
- SHELF: 2 levels
- REEF_NAME: 47 levels

Table S14. Fixed Effects

| Term | Estimate | Std. Error | z-value | p-value |
|---|---|---|---|---|
| (Intercept) | 5.443 | 0.092 | 58.95 | < 0.001 *** |
| OPENORCLOSED_AFTER2004 | 0.127 | 0.098 | 1.29 | 0.196 |

**Dispersion parameter**: 6.9

#### Key Interpretation

- No significant effect of protection status (OPENORCLOSED_AFTER2004O) on herbivore density (*z* = 1.29, *p* = 0.196).
- Most variance explained by reef-level random effects (REEF_NAME and SITE_NO:REEF_NAME).

### Hard Coral Cover

#### Raw data

Table S15. Summary statistics (descriptive) for hard coral cover.

| Protection Status | Mean Cover (%) | Median (%) | SD | n | SE |
|---|---|---|---|---|---|
| Closed (NTMR) | 23.7 | 22.0 | 14.6 | 345 | 0.79 |
| Open (Fished) | 21.2 | 17.2 | 15.4 | 360 | 0.81 |

#### GLMM Results (Binomial Distribution)

hc.tmb <- glmmTMB(cbind(n.points,total.points-.points)~OPENORCLOSED_AFTER2004 + (1|A_SECTOR)+(1|SHELF)+(1|REEF_NAME/SITE_NO/TRANSECT_NO),

family='binomial',

data=tot.hc)

summary(hc.tmb)

Model Specification:

**Family**: Binomial (logit link)

**Formula**:

cbind(n.points, total.points - n.points) ~ OPENORCLOSED_AFTER2004 + (1 | A_SECTOR) + (1 | SHELF) + (1 | REEF_NAME/SITE_NO/TRANSECT_NO)

**Dataset: Hard coral cover**

**Observations**: 705

Table S16. Model Fit Statistics

| Statistic | Value |
|---|---|
| AIC | 6467.1 |
| BIC | 6499.0 |
| Log-Likelihood | -3226.5 |
| Deviance | 6453.1 |
| Df.resid | 698 |

Table S17. Random Effects Variance

| Groups | Variance | Std. Dev. |
|---|---|---|
| SHELF | 0.0869 | 0.295 |
| REEF_NAME | 0.1488 | 0.386 |
| SITE_NO:REEF_NAME | 0.0739 | 0.272 |
| TRANSECT_NO:SITE_NO:REEF_NAME | 0.1584 | 0.398 |
| A_SECTOR | 0.0313 | 0.177 |

Groups:

- A_SECTOR, 7 levels;
- SHELF, 2 levels;
- TRANSECT_NO:SITE_NO:REEF_NAME, 705 levels;
- SITE_NO:REEF_NAME, 141 levels;
- REEF_NAME, 47 levels;

Table S18. Fixed Effects

| Term | Estimate | Std. Error | z-value | p-value |
|---|---|---|---|---|
| (Intercept) | 0.0222 | 0.238 | 0.093 | 0.926 |
| OPENORCLOSED_AFTER2004O | -0.0085 | 0.129 | -0.066 | 0.947 |

Estimated Marginal Means (Probability Scale)

| Status | Probability | SE | 95% CI |
|---|---|---|---|
| Closed | 0.506 | 0.060 | 0.391 - 0.620 |
| Open | 0.503 | 0.060 | 0.389 - 0.618 |

Contrasts

| Comparison | Estimate | SE | z-value | p-value |
|---|---|---|---|---|

| | | | | |
|---|---|---|---|---|
| Closed - Open | 0.00213 | 0.0323 | 0.066 | 0.947 |

##### Key Findings

- **No significant difference** in hard coral cover between NTMR and fished areas (z = -0.066, p = 0.947).
- Model explains moderate variation through reef- and site-level random effects.
- Estimated probabilities nearly identical (50.6% NTMR vs 50.3% fished).

### Turf algae

#### Raw data

Table S19. Summary statistics (descriptive) for turf algae cover.

| Protection Status | Mean Cover (%) | Median (%) | SD | n | SE |
|---|---|---|---|---|---|
| Closed (NTMR) | 50.8 | 51.8 | 17.7 | 345 | 0.95 |
| Open (Fished) | 52.8 | 53.0 | 14.2 | 360 | 0.75 |

#### GLMM Results (Binomial Distribution)

hc.tmb <- glmmTMB(cbind(n.points,total.points-.points)~OPENORCLOSED_AFTER2004 + (1|A_SECTOR) +(1|SHELF)+(1|REEF_NAME/SITE_NO/TRANSECT_NO),

family='binomial',

data=ta)

summary(hc.tmb)

##### Model specification

**Family**: Binomial (logit link)

**Formula**: cbind(n.points, total.points - n.points) ~ OPENORCLOSED_AFTER2004 + (1 | A_SECTOR) + (1 | SHELF) + (1 | REEF_NAME/SITE_NO/TRANSECT_NO)
**Dataset**: turf algae
**Observations**: 705

**Table S20. Model Fit Statistics**

| Statistic | Value |
|---|---|
| AIC: | 6467.1 |
| BIC: | 6499.0 |
| Log-Likelihood: | -3226.5 |

**Table S21. Random Effect Variance**

| Groups | Variance | Std. dev. |
|---|---|---|
| SHELF | 0.0869 | 0.295 |
| REEF_NAME | 0.1488 | 0.386 |
| SITE_NO:REEF_NAME | 0.0739 | 0.272 |
| TRANSECT_NO:SITE_NO:REEF_NAME | 0.1584 | 0.398 |
| A_SECTOR | 0.0313 | 0.177 |

**Table S22. Fixed Effects**

| Term | Estimate | Std. Error | z-value | p-value |
|---|---|---|---|---|
| (Intercept) | 0.0222 | 0.238 | 0.093 | 0.926 |
| OPENORCLOSED_AFTER2004O | -0.0085 | 0.129 | -0.066 | 0.947 |

Estimated Marginal Means (Probability Scale)

| Status | Probability | SE | 95% CI |
|---|---|---|---|
| Closed | 0.506 | 0.060 | 0.391 - 0.620 |
| Open | 0.503 | 0.060 | 0.389 - 0.618 |

Contrasts

| Comparison | Estimate | SE | z-value | p-value |
|---|---|---|---|---|
| Closed - Open | 0.00213 | 0.0323 | 0.066 | 0.947 |

#### Key Findings

No significant difference in turf algae cover between protection zones (z = -0.066, p = 0.947).

### Redfield ratio | Is the origin of nutrients different between NTMRs and fished reefs?

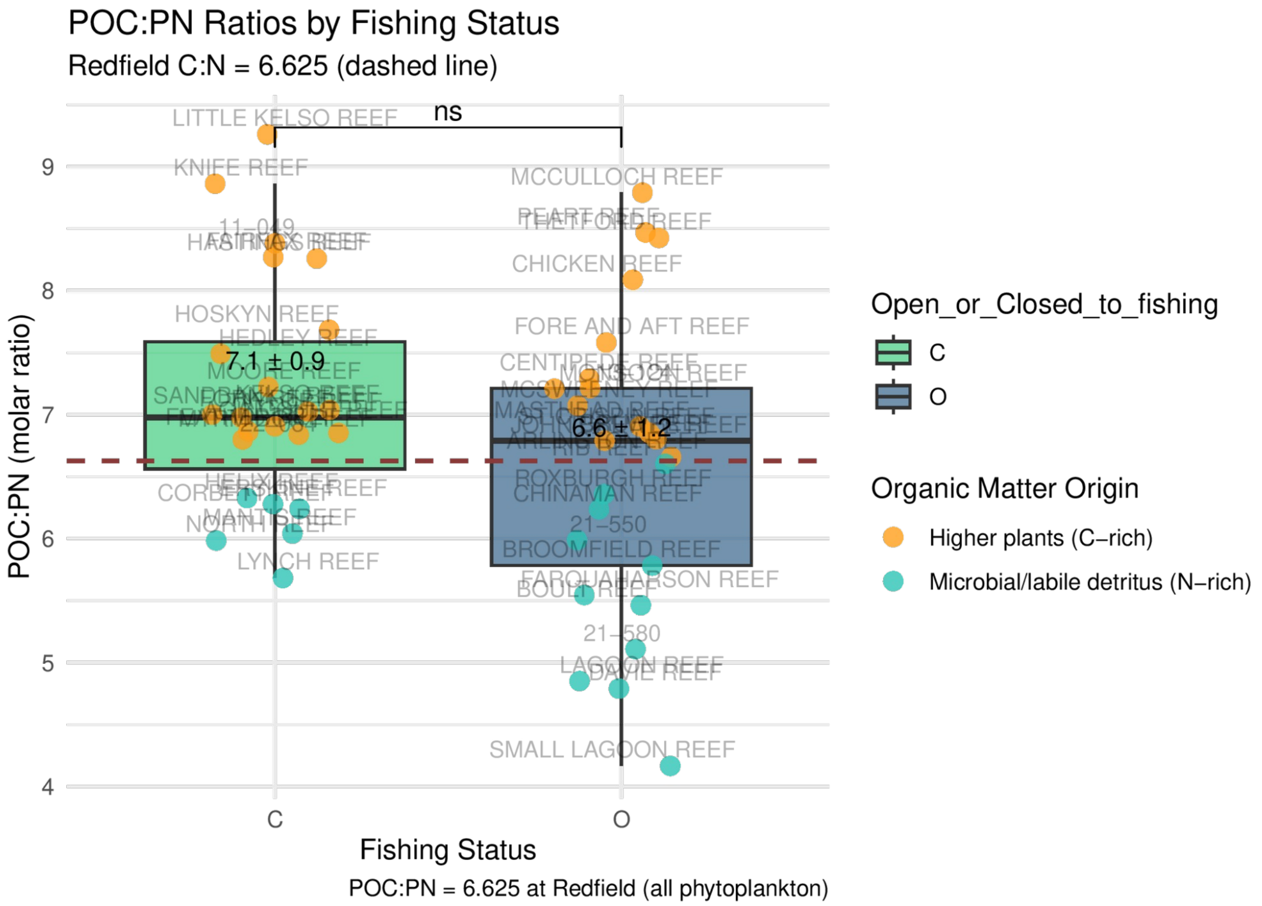


**Figure S24. The POC:PN ratio** between NTMRs and fished reefs showing which sites were C- or N-enriched. POM with more carbon indicates origins primarily of plant material (macroalgae, seagrass) whereas N-enriched POM is more likely of bacterial origin, and contains more labile detritus. The POC:PN ratio of 6.625 at Redfield would be totally phytoplanktonic in origin, and is marked with a red dashed line. Group-level comparisons were tested with a Wilcoxon rank sum test, and significance levels are indicated as: *p < 0.05; **p < 0.01; ***p < 0.001; ****p < 0.0001; "ns" = not significant.

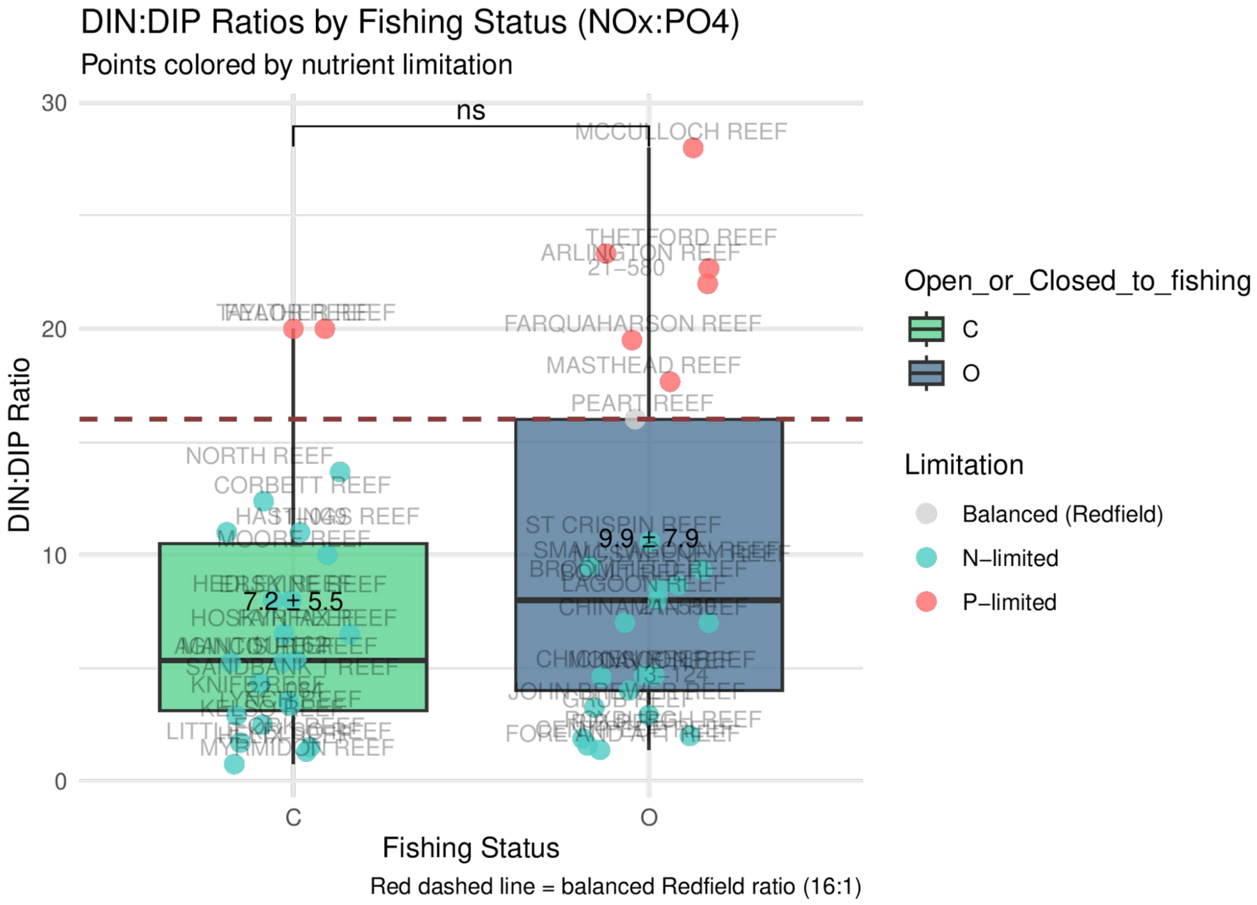


**Figure S25. The DIN:DIP ratio** between NTMRs and fished reefs showing which sites were N- or P-limited compared to the balanced redfield ratio of 16:1 (marked with a red dashed line). Group-level comparisons were tested with a Wilcoxon rank sum test, and significance levels are indicated as: *$p < 0.05$; **$p < 0.01$; ***$p < 0.001$; ****$p < 0.0001$; "ns" = not significant.

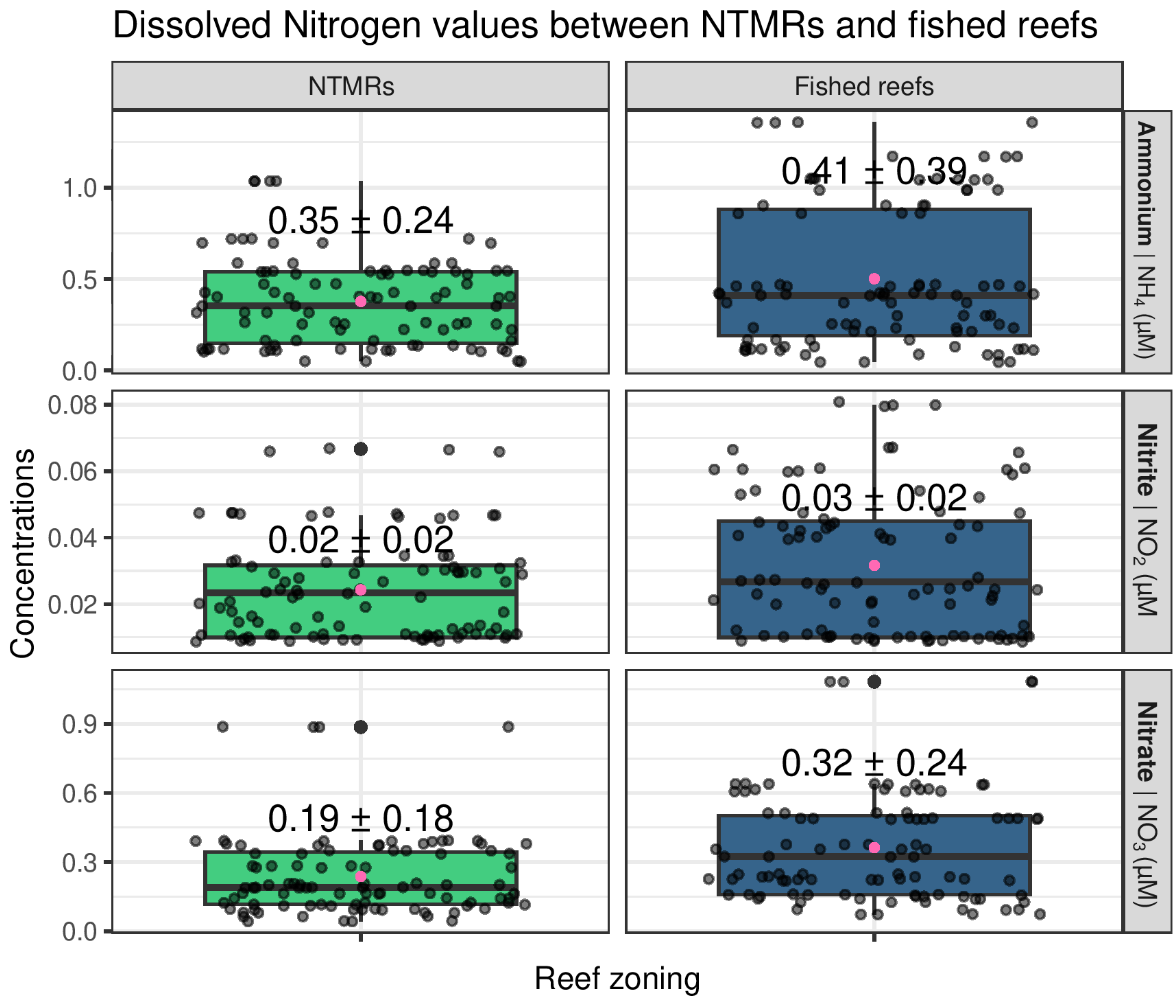


**Figure S26. Dissolved Nitrogen data**. Median ± SD values for ammonium, nitrite, and nitrate, summarised across the reef zones (NTMRs vs fished reefs). Group-level comparisons were tested with a Wilcoxon rank sum test, presented in Table S23 (below).

**Table S23. Wilcoxon rank sum tests comparing dissolved Nitrogen (ammonium, nitrite, and nitrate) values between NTMRs and fished reefs**. Significance levels are indicated as: *p < 0.05; **p < 0.01; ***p < 0.001; ****p < 0.0001; “ns” = not significant.

| variable | group1 | group2 | n1 | n2 | statistic | p.adj.bonferroni | p.adj.signif |
|---|---|---|---|---|---|---|---|
| NH4_µM | C | O | 95 | 95 | 4041 | 0.214 | ns |
| NO2_µM | C | O | 95 | 95 | 3824 | 0.065 | ns |
| NO3_µM | C | O | 95 | 95 | 2980 | 0.0000527 | **** |

### Microbial co-occurrence networks

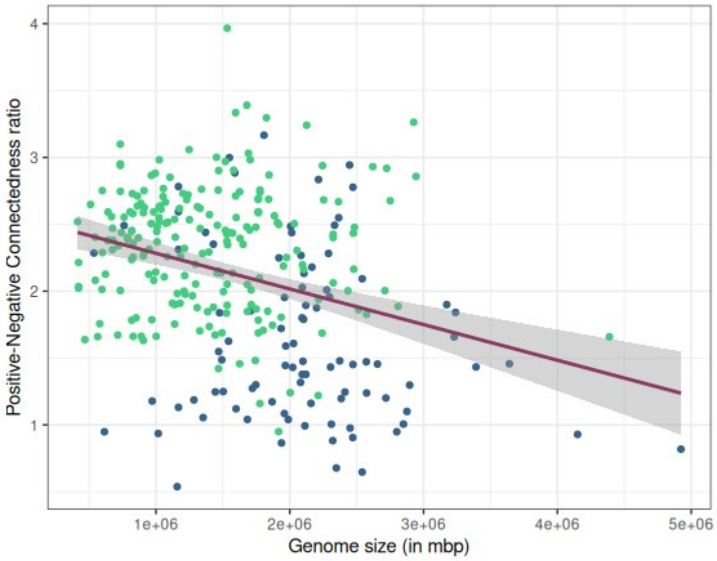


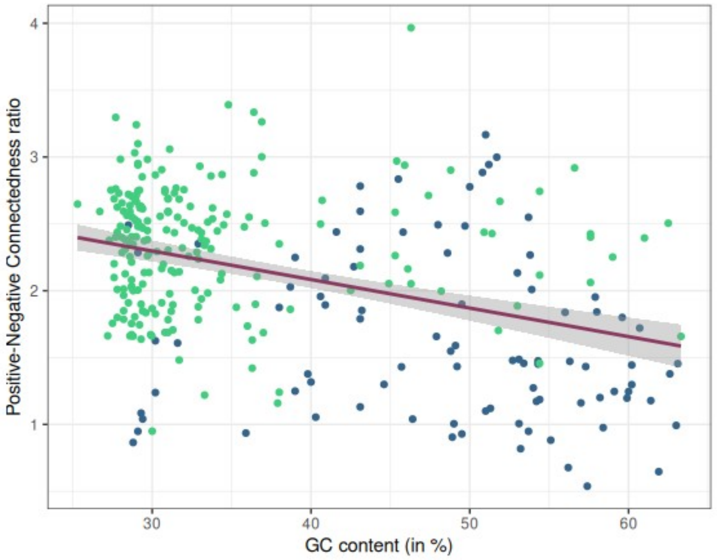


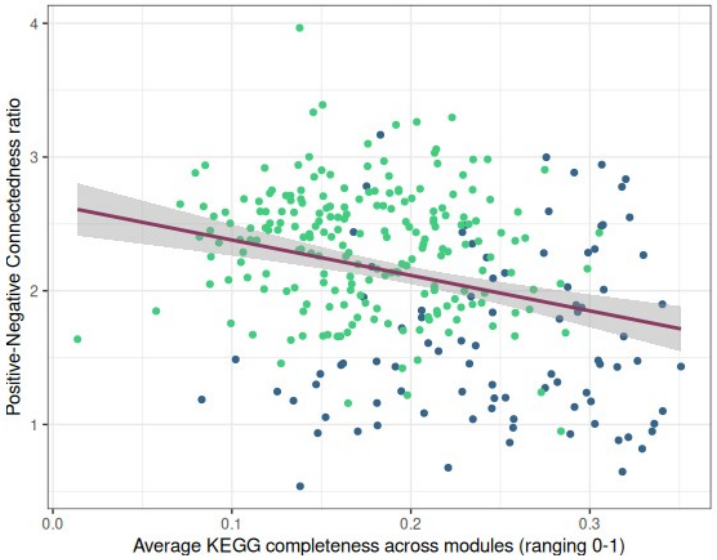

**Figure S27.** Linear regression models linking microbial genome features including genome size (**top**), GC content (**middle**), and metabolic pathway completeness (**bottom**) to network connectedness (positive:negative edge ratio) as the response variable, for metagenome-assembled genomes (MAGs) indicative of No-Take Marine Reserves (NTMRs; green) and fished reefs (blue).

The positive:negative cohesion ratio computed for each reef sample (48 reefs x 4 replicates), and visualised separately for each of the 7 GBR sectors. The boxplots show inner quartiles and median positive:negative cohesion ratio (shown as an absolute value) on the y axis, and a higher value indicates a prevalence of positive (i.e. symbiosis, metabolic co-dependency) compared to negative (i.e. competition) interactions in the microbial community. Specifically for reef zoning, this positive:negative cohesion ratio was consistently higher in NTMRs (median ± SD of positive:negative cohesion in: Cape Grenville - CG: 1.17 ± 0.10; Princess Charlotte bay - PC: 1.12 ± 0.08; Cairns - CA: 1.52 ± 0.05; Innisfail - IN: 1.20 ± 0.08; Townsville - TO: 1.31 ± 0.22; Swains - SW: 1.22 ± 0.10) compared to fished reefs (median ± SD of positive:negative cohesion ratios equalling to CG: 1.00 ± 0.16; PC: 0.97 ± 0.07; CA: 1.36 ± 0.13; IN: 1.03 ± 0.04; TO: 0.99 ± 0.19; SW: 1.13 ± 0.17) across 6 GBR sectors, apart from the Capricorn Bunker (CB) sector (with median ± SD of positive:negative cohesion being slightly higher in fished reefs: 1.08 ± 0.11, compared to NTMRs: 1.05 ± 0.09) (**Fig. S28**). Based on the results of the Wilcoxon Rank Sum tests, the positive:negative cohesion ratios were significantly higher in NTMR compared to fished zones in the following sectors: CG (W = 111, p-adj = 0.02), PC (W = 47, p-adj = 0.03), CA (W = 86, p-adj = 0.002), IN (W = 162, p-adj = 0.0002), and TO (W = 569, p-adj = 0.003). However, no significant difference was found in the southern GBR sectors SW (W = 59, p-adj = 0.427) and CB (W = 93, p-adj = 0.91).

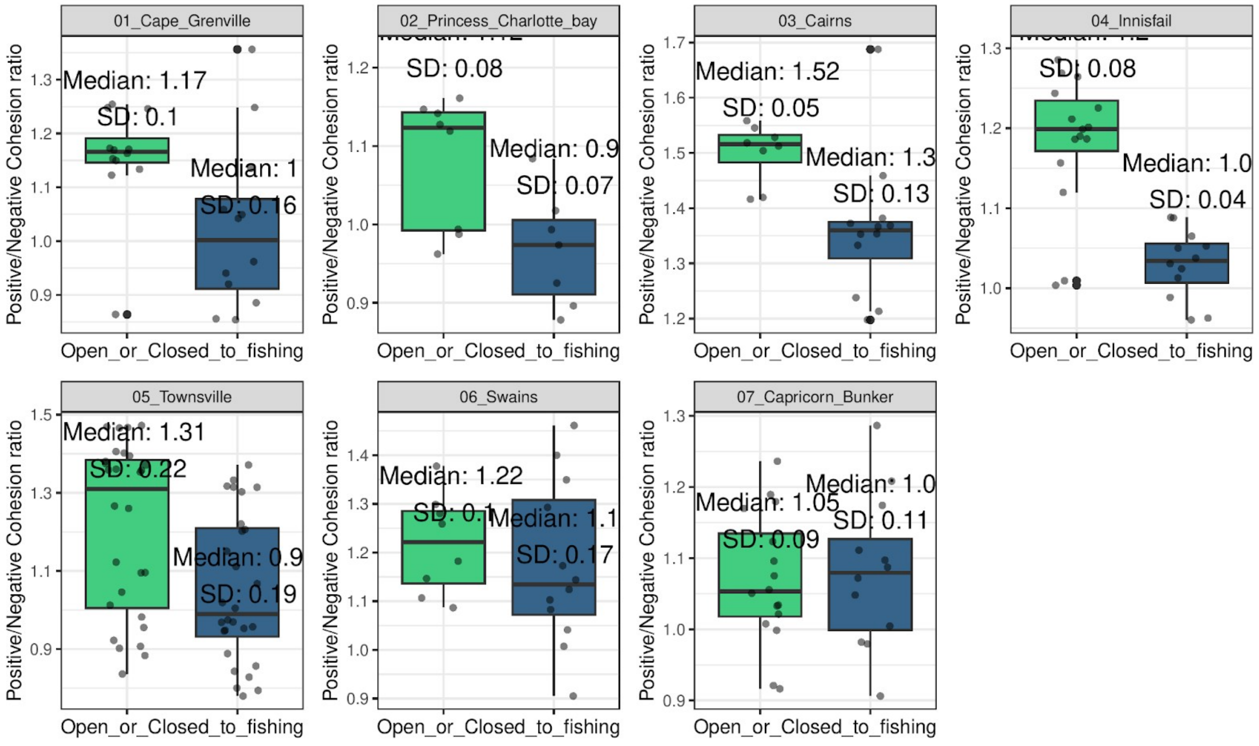


**Figure S28**. Sample-level positive:negative cohesion ratios indicate the prevalence of positive (increase in positive to negative cohesion ratio) in or negative (increase in positive to negative cohesion ratio) interactions within reef bacterioplankton between fished reefs (blue) and NTMRs (green), in each of the 7 GBR sectors we sampled.

Further, the higher positive:negative cohesion was also observed for the sites sampled in the winter trip, suggesting a prevalence of positive interactions (mutualism, co-occurrence due to metabolic exchange) in the winter (**Fig. S29**) when nutrients are depleted (**Fig. S30**). In contrast, we see a potential increase of negative/mutually exclusive interactions (predator/prey, pathogen/host, parasite/host, and etc.) in the summer trips (**Fig. S29**) when nutrients are elevated (**Fig. S30**), potentially indicating that opportunistic microbes are competing for available nutrients that are elevated in the summer transects.

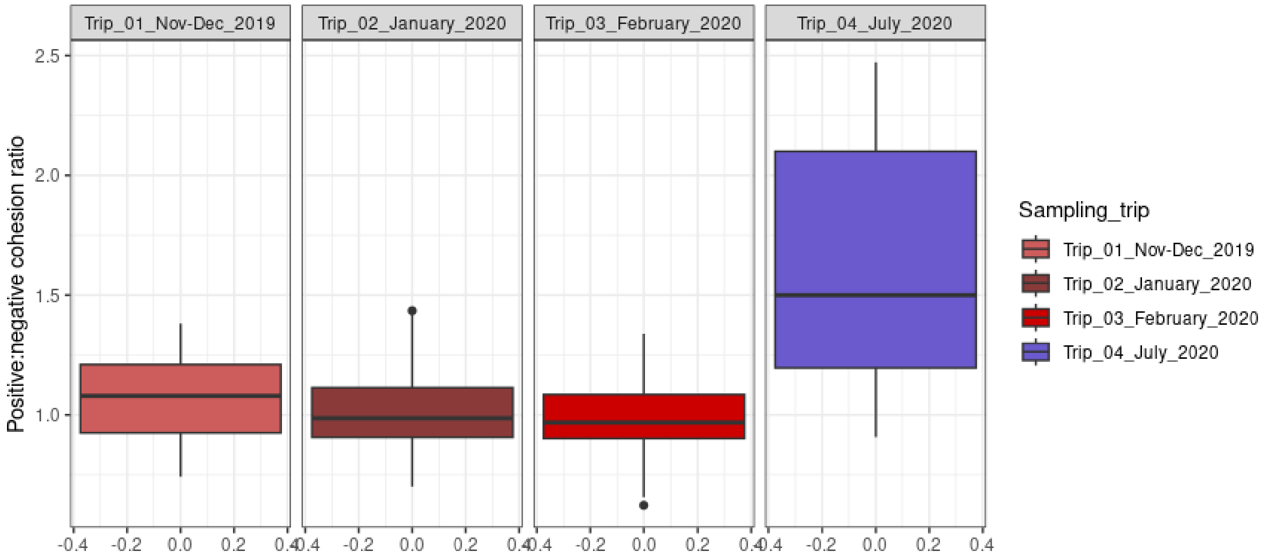


**Figure S29**. Sample-level positive:negative cohesion ratios indicate the prevalence of positive (increase in positive to negative cohesion ratio) in or negative (increase in positive to negative cohesion ratio) interactions within reef bacterioplankton between summer sampling transects (Trips 1-3, red) and the winter trip (Trip 4; blue).

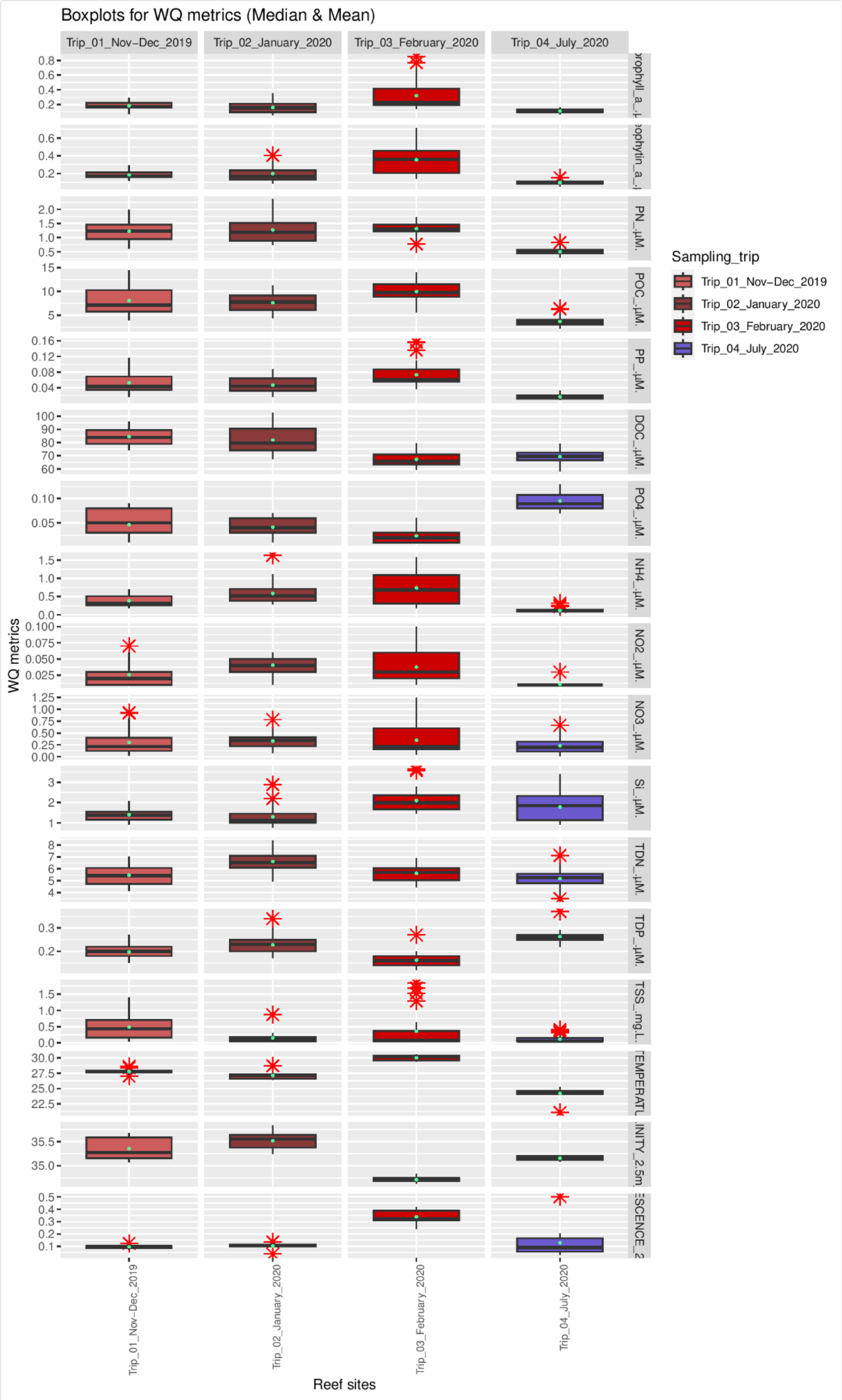


**Figure S30: Physico-chemical data**. Median ± SD values of 17 physico-chemical variables collected. Values are summarised across the four sampling trips, with the colour code corresponding to Fig. 1 in the main text. Acronyms explained: ammonia ($NH_4^+$), nitrite ($NO_2^-$), nitrate ($NO_3^-$), total dissolved nitrogen (TDN), phosphate ($PO_4^{3-}$), total dissolved phosphorus (TDP), dissolved organic carbon (DOC), silicate (Si), total suspended solids (TSS), chlorophyll a (Chl-*a*), phaeophytin a (Phaeo), particulate organic carbon (POC), particulate nitrogen (PN), and particulate phosphorus (PP).

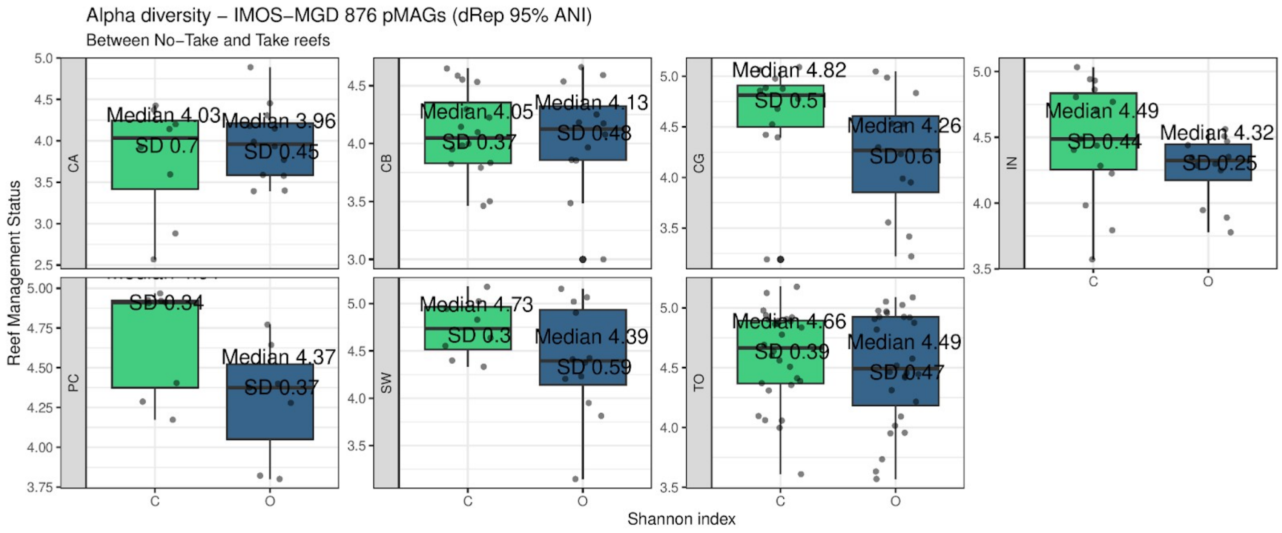


**Figure S31. Microbial diversity between the zones.** Alpha diversity (Shannon index) of seawater microbiomes between NTMRs (green) and fished reef sites (blue) across the 7 sampled GBR sectors.

### Within- versus between-module connectivity in NTMR and fished-reef networks

Network stability is thought to depend not only on the overall density of associations but on their topological arrangement: communities in which associations are concentrated within modules (tightly linked subgroups) are predicted to better contain environmentally induced perturbations, whereas communities with a higher proportion of between-module connections may allow disturbances to propagate more widely. Because the elevated positive:negative cohesion we observed in NTMRs (Fig. 4I) does not by itself indicate greater stability (since dense cooperative coupling can in fact be destabilising, while competition can be stabilising[27]), we assessed whether the two zones differed in network compartmentalisation directly.

For each of the 14 sector×zone co-occurrence networks, we used the Clauset-Newman-Moore module assignments (as computed for the modularity analysis) to classify every edge as within-module (both endpoints assigned to the same module) or between-module. We then computed, per network, the within-module edge fraction (within-module edges divided by total edges) and the within:between edge ratio, and compared each metric between NTMRs and fished reefs using Mann-Whitney U tests on the seven sector-specific values per zone.

Neither metric differed significantly between zones. The within-module edge fraction was near-identical (NTMR 0.689 ± 0.045; fished 0.696 ± 0.029; Mann-Whitney U, $p = 0.535$), as was the within:between edge ratio (NTMR 2.28 ± 0.57; fished 2.32 ± 0.32; $p = 0.535$). Across all networks, roughly two-thirds to three-quarters of edges fell within modules in both zones (within-module fraction range 0.64–0.78), and this proportion showed no systematic association with protection status (**Fig. S32**). Consistent with the absence of a significant modularity difference between zones (**Fig. 4J**), these results indicate that NTMR and fished-reef networks are similarly compartmentalised, hence our data did not identify differences in stability of seawater microbiome communities between co-located NTMR and fished sites sampled here.

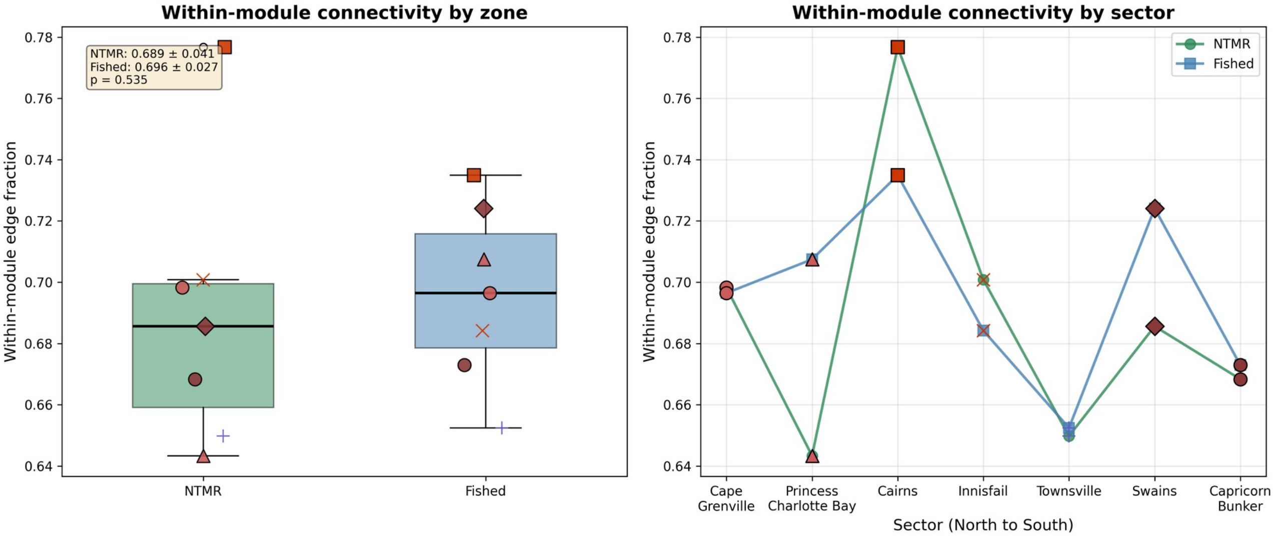


**Figure S32. Within- versus between-module connectivity in NTMR and fished-reef co-occurrence networks**. Within-module edge fraction (within-module edges / total edges) for each of the 14 sector×zone FlashWeave networks, shown by zone (left; boxplots with sector-specific markers and trip-based colours as in Fig. 4) and by GBR sector (right). Neither the within-module edge fraction nor the within:between edge ratio differed significantly between NTMRs and fished reefs (Mann-Whitney U, $p = 0.54$ for both), indicating similar network compartmentalisation across zones.

**Table S24. Within- versus between-module connectivity for the 14 sector×zone microbial co-occurrence networks**. Module assignments were obtained using the Clauset-Newman-Moore greedy algorithm (as for the modularity analysis). Each edge was classified as within-module (both endpoints in the same module) or between-module. The within-module edge fraction is within-module edges divided by total edges. NTMR = No-Take Marine Reserve.

| GBR sector | Zone | Nodes | Edges | Modules | Within | Between | Within-module fraction | Within:between ratio |
|---|---|---|---|---|---|---|---|---|
| Cape Grenville | NTMR | 873 | 1087 | 129 | 759 | 328 | 0.698 | 2.31 |
| Cape Grenville | Fished | 868 | 1074 | 146 | 748 | 326 | 0.696 | 2.29 |
| Princess Charlotte Bay | NTMR | 869 | 987 | 244 | 635 | 352 | 0.643 | 1.8 |
| Princess Charlotte Bay | Fished | 838 | 899 | 210 | 636 | 263 | 0.707 | 2.42 |
| Cairns | NTMR | 847 | 851 | 194 | 661 | 190 | 0.777 | 3.48 |
| Cairns | Fished | 872 | 1060 | 123 | 779 | 281 | 0.735 | 2.77 |
| Innisfail | NTMR | 857 | 1083 | 129 | 759 | 324 | 0.701 | 2.34 |
| Innisfail | Fished | 850 | 1007 | 179 | 689 | 318 | 0.684 | 2.17 |
| Townsville | NTMR | 875 | 1462 | 51 | 950 | 512 | 0.65 | 1.86 |
| Townsville | Fished | 872 | 1453 | 39 | 948 | 505 | 0.652 | 1.88 |
| Swains | NTMR | 479 | 299 | 274 | 205 | 94 | 0.686 | 2.18 |
| Swains | Fished | 873 | 1178 | 77 | 853 | 325 | 0.724 | 2.62 |
| Capricorn Bunker | NTMR | 873 | 1176 | 129 | 786 | 390 | 0.668 | 2.02 |
| Capricorn Bunker | Fished | 869 | 1058 | 180 | 712 | 346 | 0.673 | 2.06 |
| NTMR mean ± SD | | | | | | | 0.689 ± 0.045 | 2.28 ± 0.57 |
| Fished mean ± SD | | | | | | | 0.696 ± 0.029 | 2.32 ± 0.32 |
| Mann-Whitney U | | | | | | | p = 0.535 | p = 0.535 |

**KEGG Pathway** analysis | Which metabolic traits are enriched in microbial indicators of NTMRs vs. fished reefs?

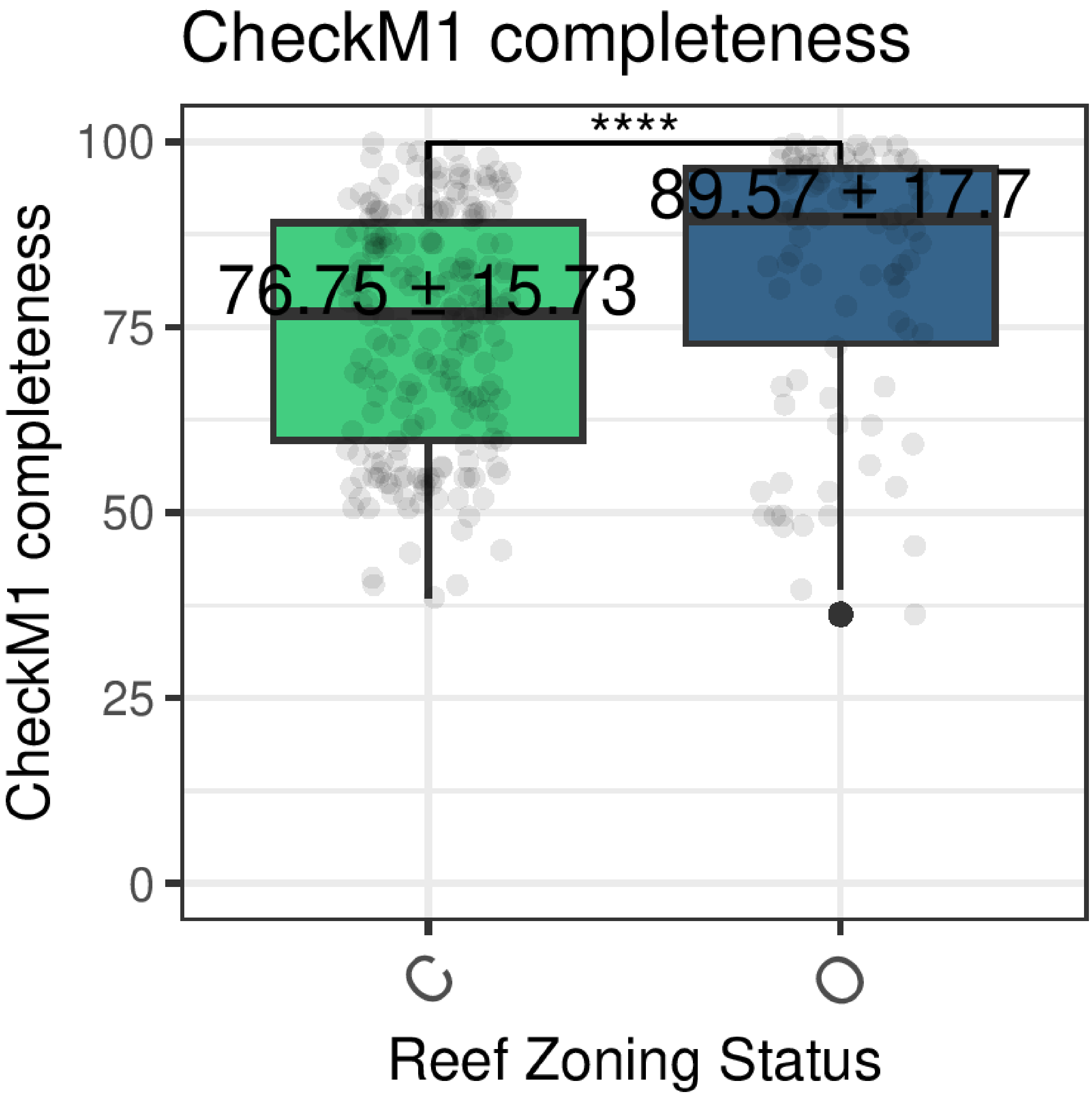


**Figure S33**. Boxplots comparing CheckM1 genome completeness score distributions between microbial indicators of NTMRs vs fished reefs. Significance levels from Wilcoxon rank sum tests are indicated as: *$p < 0.05$; **$p < 0.01$; ***$p < 0.001$; ****$p < 0.0001$; "ns" = not significant.

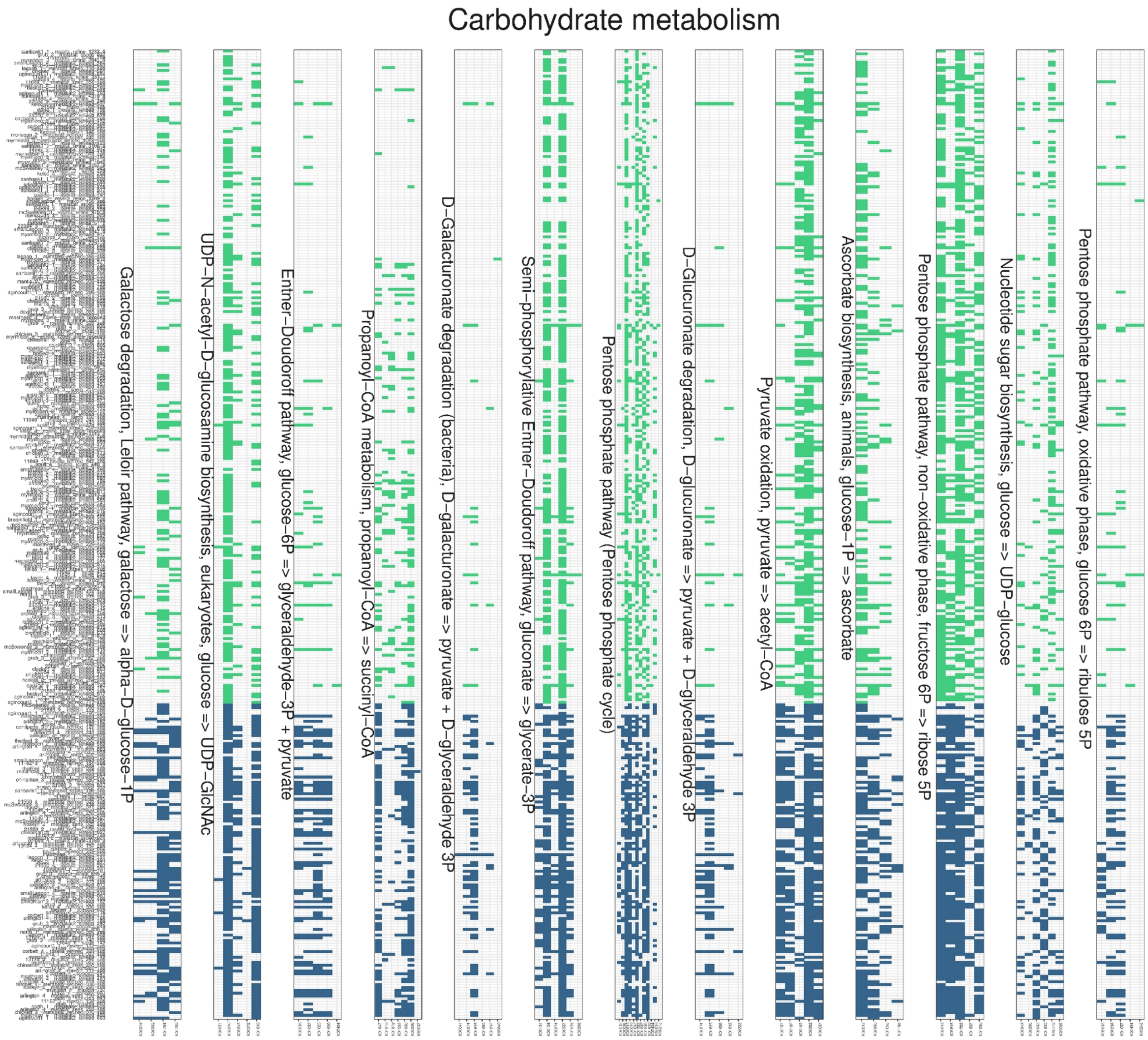


**Figure S34. Gene level module completeness (Carbohydrate metabolism).**

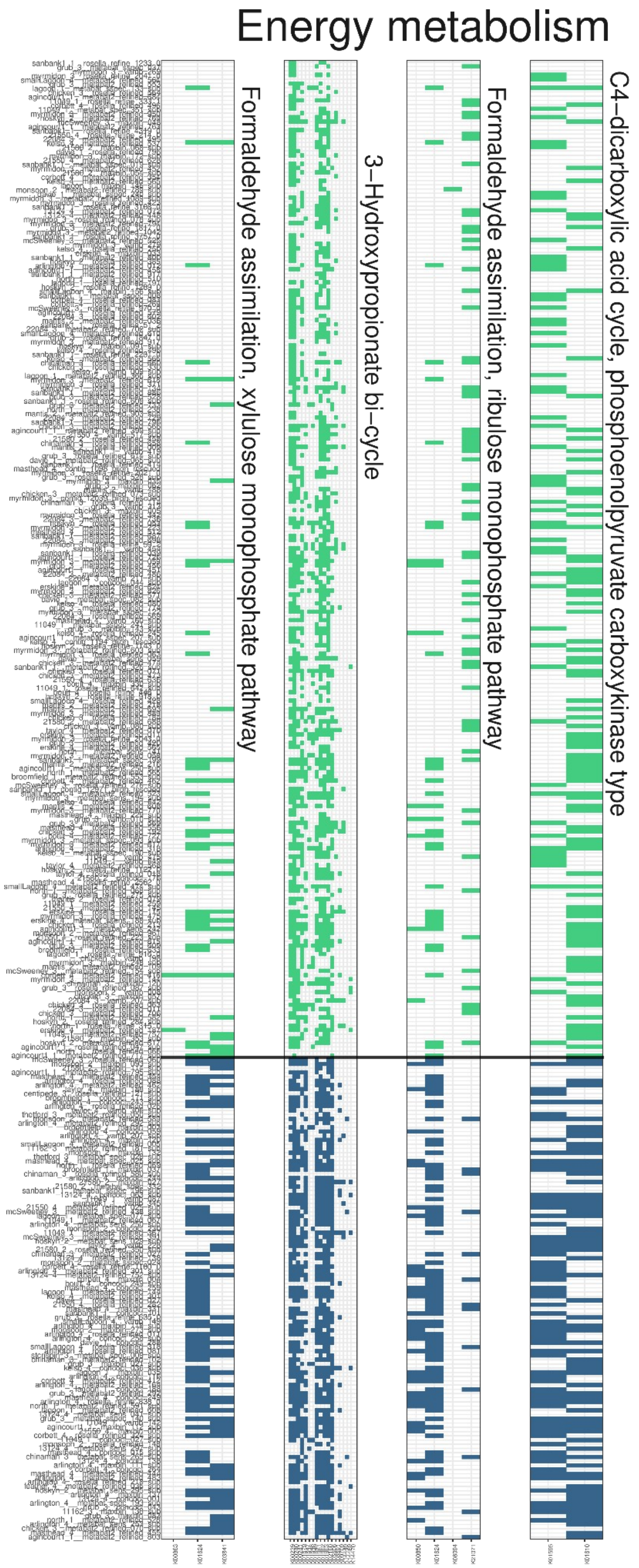


**Figure S35. Gene level module completeness (Energy metabolism).**

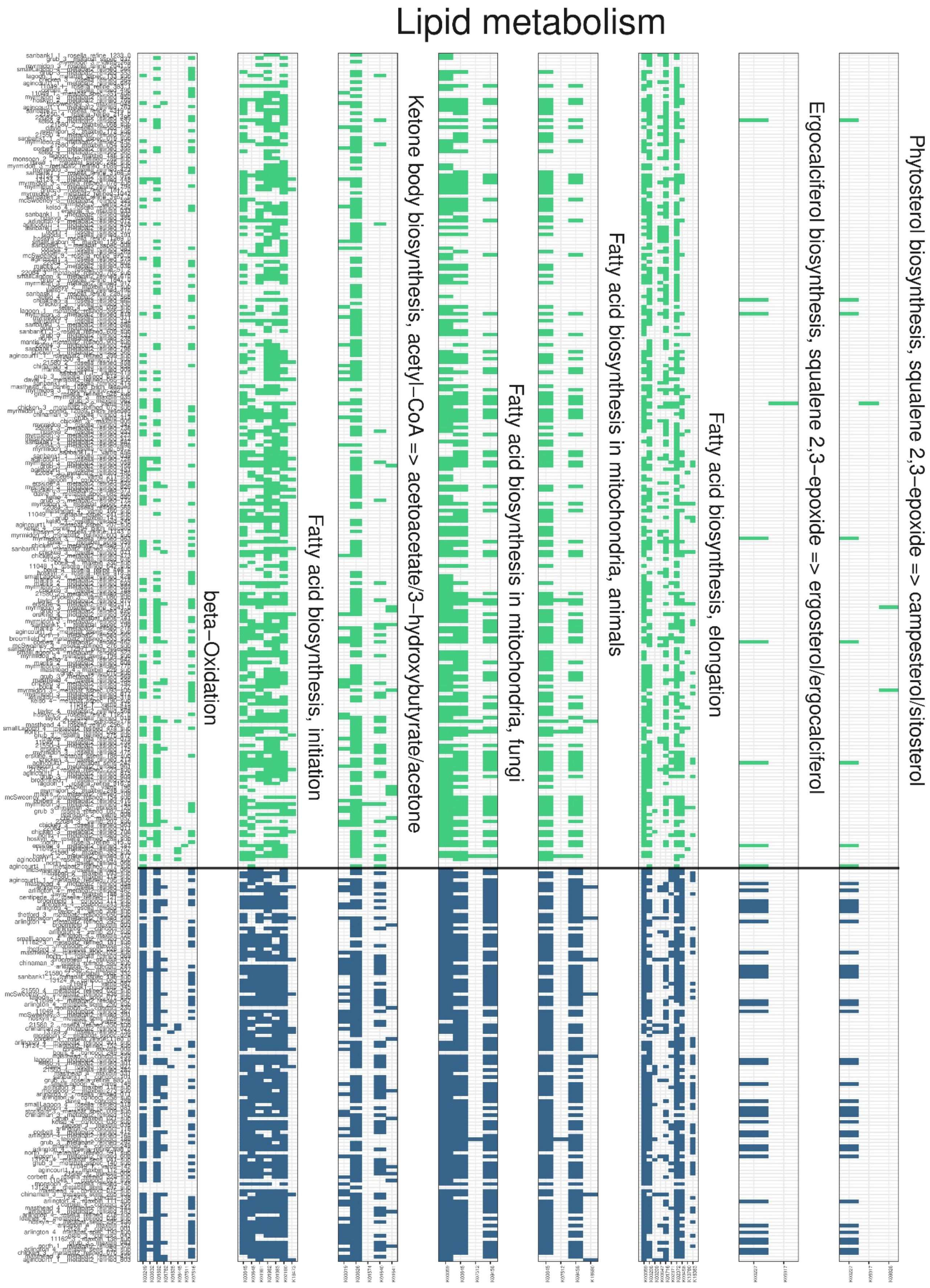


**Figure S36. Gene level module completeness (Lipid metabolism).**

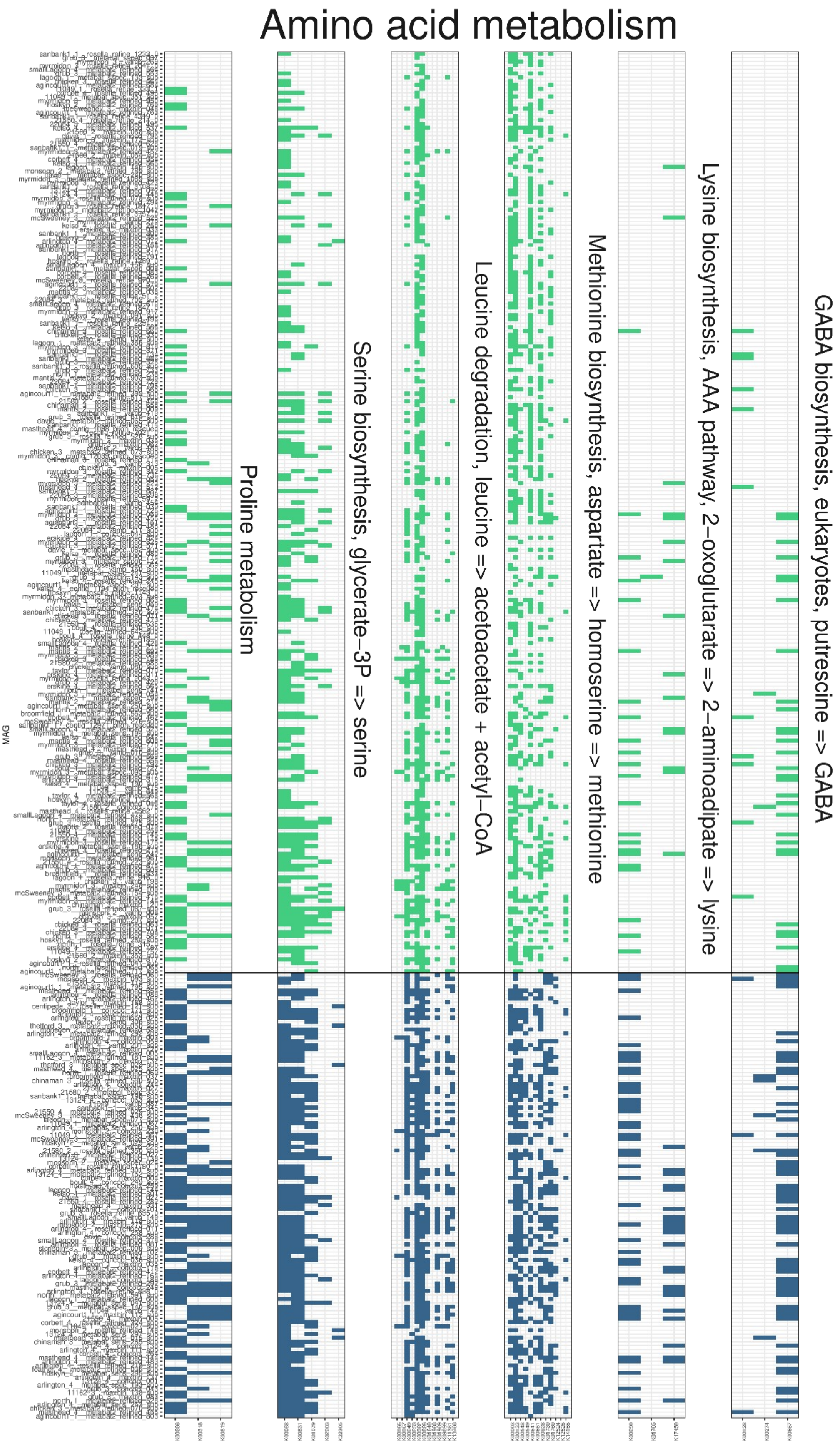


**Figure S37. Gene level module completeness (Amino acid metabolism).**

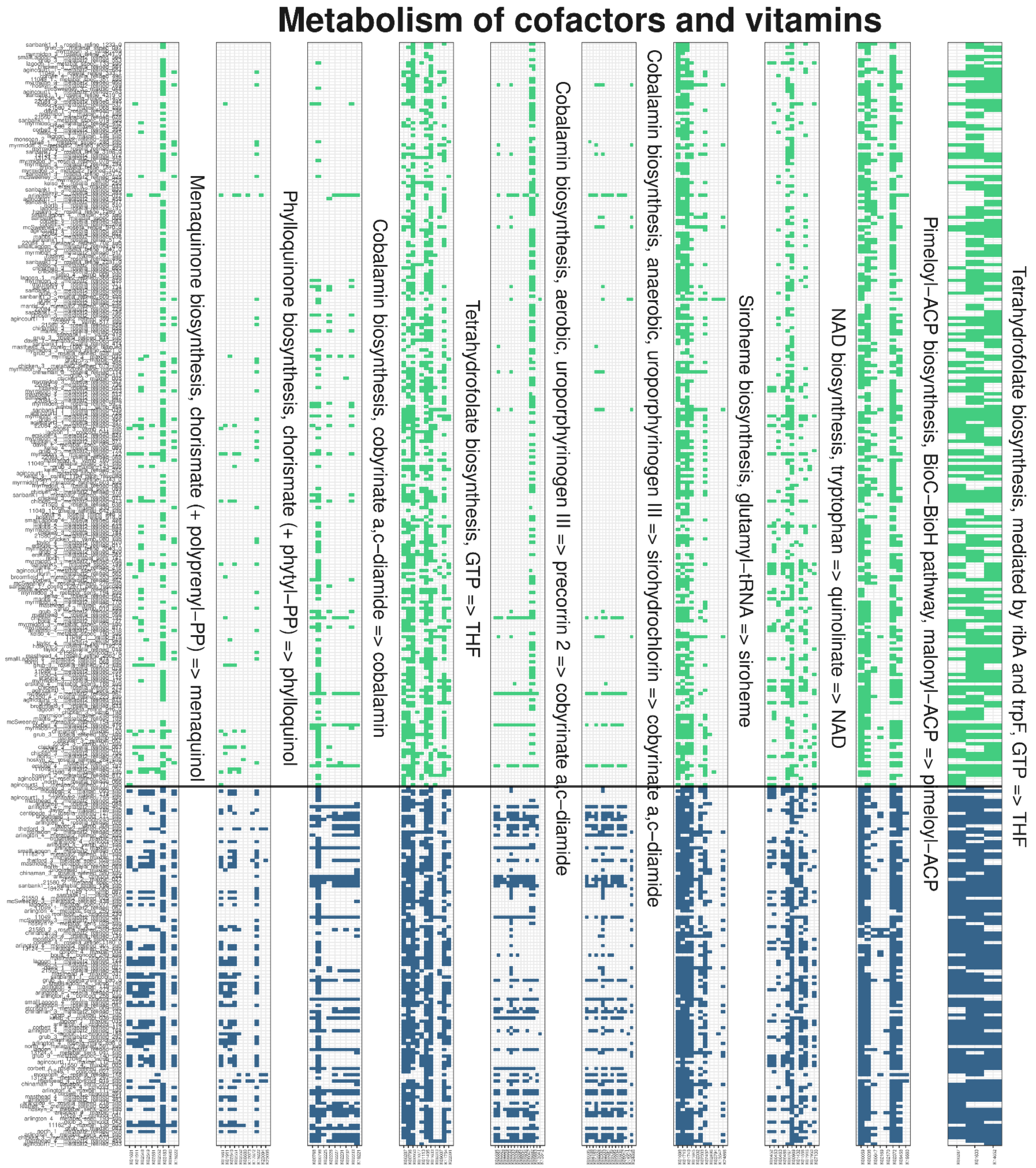


**Figure S38. Gene level module completeness (Metabolism of cofactors and vitamins).**

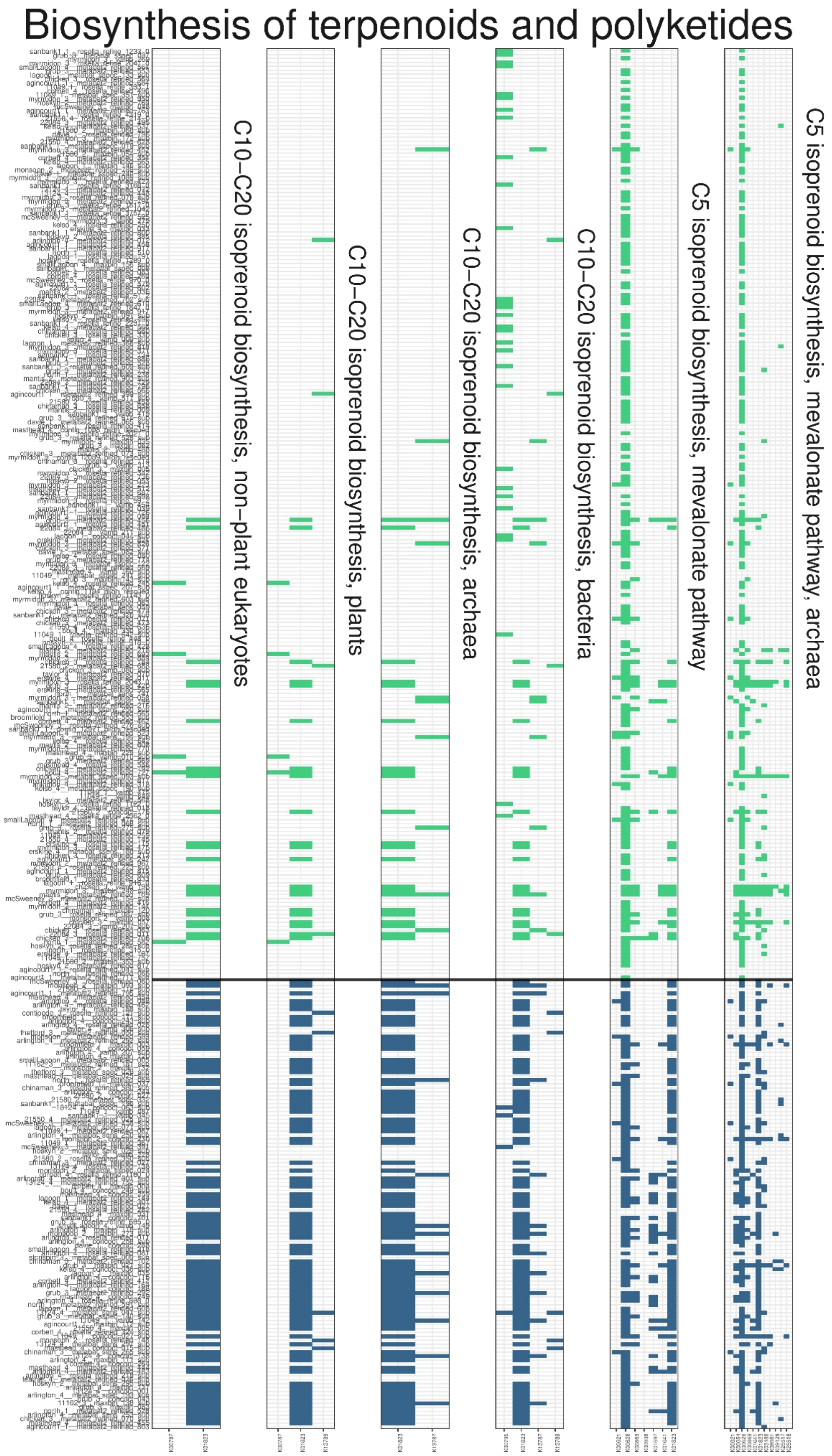


**Figure S39. Gene level module completeness (Biosynthesis of terpenoids and polyketides).**

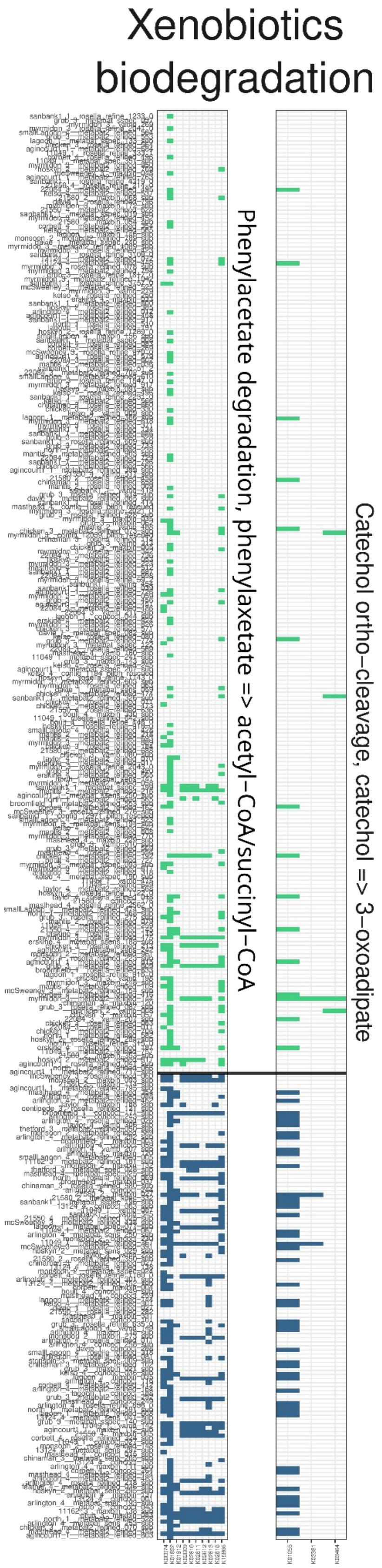


**Figure S40. Gene level module completeness (Xenobiotics biodegradation).**

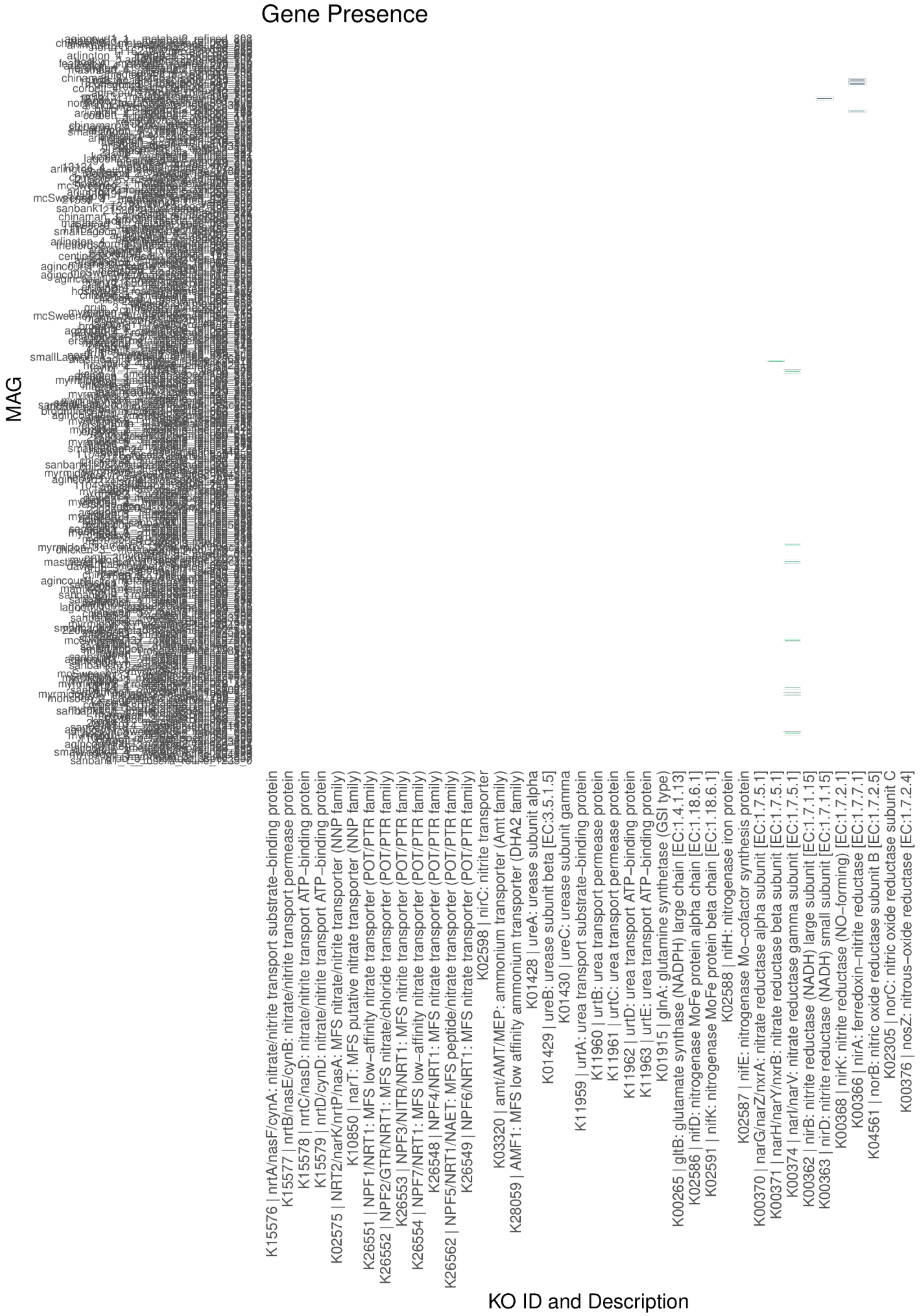


**Figure S41. Functional potential for nitrogen acquisition in zoning-indicator MAGs**. Heatmap of nitrate, nitrite, ammonium, and urea transporter and metabolic genes, which were colored if present (NTMR-enriched microbes in green; fished-reef enriched microbes in blue).

### Read-based metagenomic functional validation using GO terms

To provide a community-wide functional perspective complementary to the KEGG module completeness analysis of indicator pMAGs (**Fig. 5; main text**), we performed MINT sPLS-DA on the read-based functional dataset (4,287 GO terms at rank 5) generated from the same metagenomic reads (Illumina-only), previously published in Terzin et al. (2025)[21]. Briefly, quality-filtered reads were aligned against the NCBI nr database using DIAMOND (v2.0.9) with an e-value threshold of $< 1 \times 10^{-5}$. Resulting alignments were imported into MEGAN (v6.23.0), where GO term annotations were assigned to each read and collapsed at GO rank 5. GO term abundance counts were exported and imported into R using the phyloseq R package. Pre-filtering steps included removal of: (1) non-annotated reads; (2) reads annotated as eukaryotic or viral; and (3) rare/spurious GO terms with relative abundance < 0.0001%. This filtering resulted in a final dataset of 4,287 GO terms. Microbial functional abundance data were then center log-ratio (CLR) transformed using the microbiome R package to account for sparsity and compositional nature of metagenomic sequencing data, with pseudocounts introduced prior to CLR transformation as log 0 is undefined.

MINT sPLS-DA was applied with reef zoning status as the categorical outcome and GBR sector as the grouping factor ("study" in MINT[26]) to account for spatio-temporal batch structure, following identical procedures to those described for the read-based taxonomic validation above. Model tuning identified the optimal number of components and features per component based on the balanced error rate (BER) under leave-one-group-out cross-validation (LOGOCV), with GBR sectors as groups. Sample plots revealed separation between NTMRs and fished reefs along component 1, with no residual clustering by sampling trip, confirming that spatiotemporal batch effects were effectively accounted for (**Fig. S42A**). The Clustered Image Map (CIM) for components 1–2 showed distinct enrichment patterns between zones (**Fig. S42B**).

GO terms enriched in NTMRs included those associated with efficient nutrient acquisition, such as (1) IPR015862 (MglA-type sugar ABC transporters), which facilitate high-affinity scavenging of dissolved monosaccharides including glucose and galactose from oligotrophic seawater[28]; and (2) IPR024919 (EcfT — Energy-coupling factor transporter transmembrane protein), a class of ABC transporters found exclusively in bacteria and archaea that mediate uptake of essential vitamins and micronutrients including cobalamin (B12), thiamine (B1), folate, and riboflavin at extremely low environmental concentrations[29], and whose importance in oligotrophic marine bacteria is underscored by the demonstrated requirement of SAR11 for exogenous vitamin precursors from seawater[30]. This also aligns with our MAG-based analysis which showed that NTMR-enriched pMAG indicators had far lower completeness in cobalamin biosynthesis pathways compared to fished reef indicators (**Fig. 5, main text; Fig. S37-39**), implying NTMR microbes scavenge vitamins from the environment rather than synthesising them *de novo*. Further, NTMR-enriched functional genes (GO terms) were involved in core cellular maintenance, including translation (GO:0006412), biosynthetic processes (GO:0009058), and lyase activity (GO:0016829; see **Fig. S42**), which are well-established signatures of constitutively expressed housekeeping functions in streamlined marine oligotrophs (such as *Pelagibacter*, SAR86, and *Prochlorococcus* — indicative of NTMRs; see **Fig. 2, main text; Fig. S22**), consistent with the notion that oligotrophic marine bacteria maintain steady expression of core metabolic genes regardless of environmental conditions[31,32].

GO terms enriched in fished reefs included those associated with active energy metabolism, nitrogen assimilation, anabolic reactions (including cofactor and vitamin biosynthesis), and motility (**Fig. S42**), broadly consistent with the copiotrophic and metabolically independent functional profile identified in fished reef indicator pMAGs (**Fig. 5, main text**). Specifically, enrichment of GO:0006091 (generation of precursor metabolites and energy) and GO:0016491 (oxidoreductase activity; see **Fig. S42**) directly echoes the MAG-based finding that fished reef indicators are restructured for rapid energy harvest from organic substrates, with more complete carbohydrate utilisation pathways (pentose phosphate, Entner-Doudoroff, and galactose degradation; **Fig. 5, main text**) converging on glycolysis and fuelling the TCA cycle for ATP generation. IPR011283 (acetoacetyl-CoA reductase), the key enzyme in polyhydroxybutyrate (PHB) biosynthesis, further extends this picture: PHB production channels excess acetyl-CoA — whose synthesis is enhanced in fished reef indicator pMAGs via β-oxidation (M00087) and pyruvate oxidation (M00307; **Fig. 5, main text**) — into carbon and energy storage under nutrient-imbalanced but carbon-rich conditions[33], a hallmark of copiotrophic boom-and-bust metabolism. Enrichment of IPR027283 (bacterial glutamate synthase large subunit) and GO:0006807 (nitrogen compound metabolic processes; see **Fig. S41**) further aligns with the MAG-based observation that fished reef indicators have enhanced amino acid biosynthetic capacity (serine, methionine, proline, tryptophan; **Fig. 5, main text**) and likely drive remineralisation of particulate organic matter to produce dissolved inorganic nitrogen. Active nitrogen assimilation (in the form of ammonium) via the GS/GOGAT pathway (Glutamine Synthetase (GS) and Glutamate Synthase (GOGAT) cycle) is mechanistically consistent with our hypothesis of fished-reef enriched bacteria incorporating inorganic nitrogen into organic carbon skeletons in anabolic reactions (**Fig 5; main text**), since GS/GOGAT pathway is enriched in environments where ATP is plentiful[34] (such as the nutrient-enriched fished reefs) — and of which IPR027283 glutamate synthase (enriched in fished reefs; see **Fig. S42**) is the central enzyme.

Further, our MAG-based finding that fished-reef pMAG indicators have substantially greater completeness of cofactor and vitamin biosynthesis pathways, including cobalamin (**Fig. 5, main text; Fig. S38**) was independently corroborated with a read-based metagenomics analysis using GO terms. Specifically, GTP cyclohydrolase 1 type 2/Nif3 (IPR002678) catalyses the first committed step of *de novo* folate (B9 vitamin) biosynthesis[35], and its enrichment in fished reefs (**Fig. S42**) further creates a compelling functional contrast with the ECF transporter signal enriched in NTMRs (IPR024919): where oligotrophic NTMR-associated communities scavenge vitamins from the environment at extremely low concentrations[29,30], fished reef communities appear to synthesise them *de novo*, which is a metabolic strategy consistent with greater genomic autonomy and copiotrophic lifestyle. Finally, fished-reef enriched GO terms involved in flagellar assembly and regulation genes — including IPR005503 (flagellar basal body-associated protein FliL), IPR023597 (flagellar regulator Flk), and IPR007412 (anti-sigma-28 factor FlgM, which gates flagellar gene expression by inhibiting the flagella-specific sigma factor FliA until hook-basal body assembly is complete) (**Fig. S42**). This provides independent corroboration of copiotrophic motility capacity in fished reef communities. Flagellar motility is a well-established copiotrophic trait enabling navigation toward nutrient hotspots in heterogeneous marine environments[36]. While flagellar motility does not appear directly in our KEGG module analysis, it is consistent with the broader competitive and antagonistic microbial interaction network patterns observed in fished reefs (**Fig. 4, main text**).

It should be noted that some GO terms in both groups likely reflect database annotation artefacts inherent to read-based NCBI nr classification via DIAMOND/MEGAN, where prokaryotic reads can be assigned to the closest eukaryotic or viral homologue in the database regardless of biological plausibility (e.g. viral capsid proteins and eukaryotic cytoskeletal proteins appearing among zone-associated GO terms); this limitation underscores the value of the MAG-based KEGG analysis as the primary, mechanistically rigorous framework. Nonetheless, the broad functional contrast between zones, including (1) efficient nutrient uptake and streamlined core metabolism in NTMRs versus (2) active energy harvest, nitrogen assimilation, and motility in fished reefs — is consistent across both the read-based GO term and

MAG-based KEGG approaches, providing independent, assembly-free support for the MAG-based functional patterns (reported in **Fig. 5; main text**).

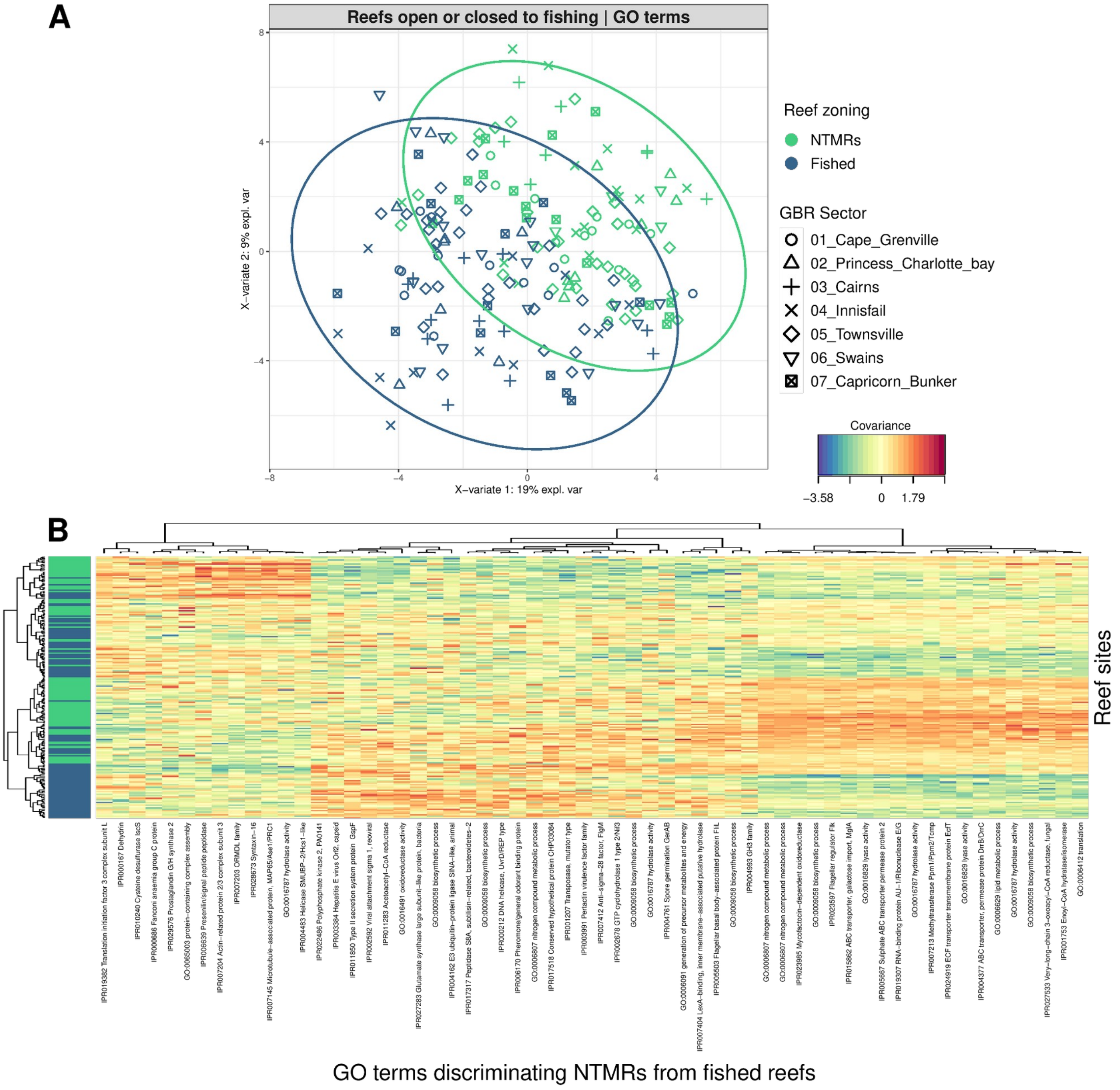


**Figure S42. Community-wide functional validation of reef zoning signals using read-based analysis (4,287 GO terms derived from DIAMOND and MEGAN read-centric profiling)**. (A) Global MINT sPLS-DA sample plot showing separation of NTMRs (green) and fished reefs (blue), with 95% confidence ellipses. (B) Clustered Image Map (CIM) of GO terms selected by MINT sPLS-DA on components 1–2, with reef sites (rows) coloured by protection status. GO terms enriched in NTMRs reflect efficient nutrient acquisition and core cellular maintenance, while those enriched in fished reefs are associated with active energy metabolism, nitrogen assimilation, and motility, broadly consistent with the functional patterns identified in the MAG-based KEGG analysis (**Fig. 5; main text**).

### Robust Optimum (RO) method | Microbial niche modelling

For each microbial predictor, we computed the microbial niche tolerance ranges using the robust optimum method (Cristóbal et al., 2014)[37], as explained in the main text. This allowed us to define the lower and upper niche limits—the environmental conditions below and above which the $pMAGs_{95\%ANI}$ cannot survive due to unfavorable conditions—as well as the ecological niche optimum, which corresponds to the environmental value at which that microbe is found at its highest relative abundances, i.e. optimal conditions for the existence, development, growth, and proliferation of that microbe (Ter Braak & van Dam, 1989). A comparative niche analysis focused on the 350 indicator $pMAGs_{95\%ANI}$ that distinguish between NTMRs and fished reefs, particularly in relation to dissolved nitrogen variables ($NH_4$, $NO_2$, and $NO_3$) which were the explanatory drivers on fished reefs. Pairwise comparisons of niche preferences for these dissolved nitrogen variables between NTMRs and fished reefs were visualised with boxplots in ggplot2 3.5.1 [38].

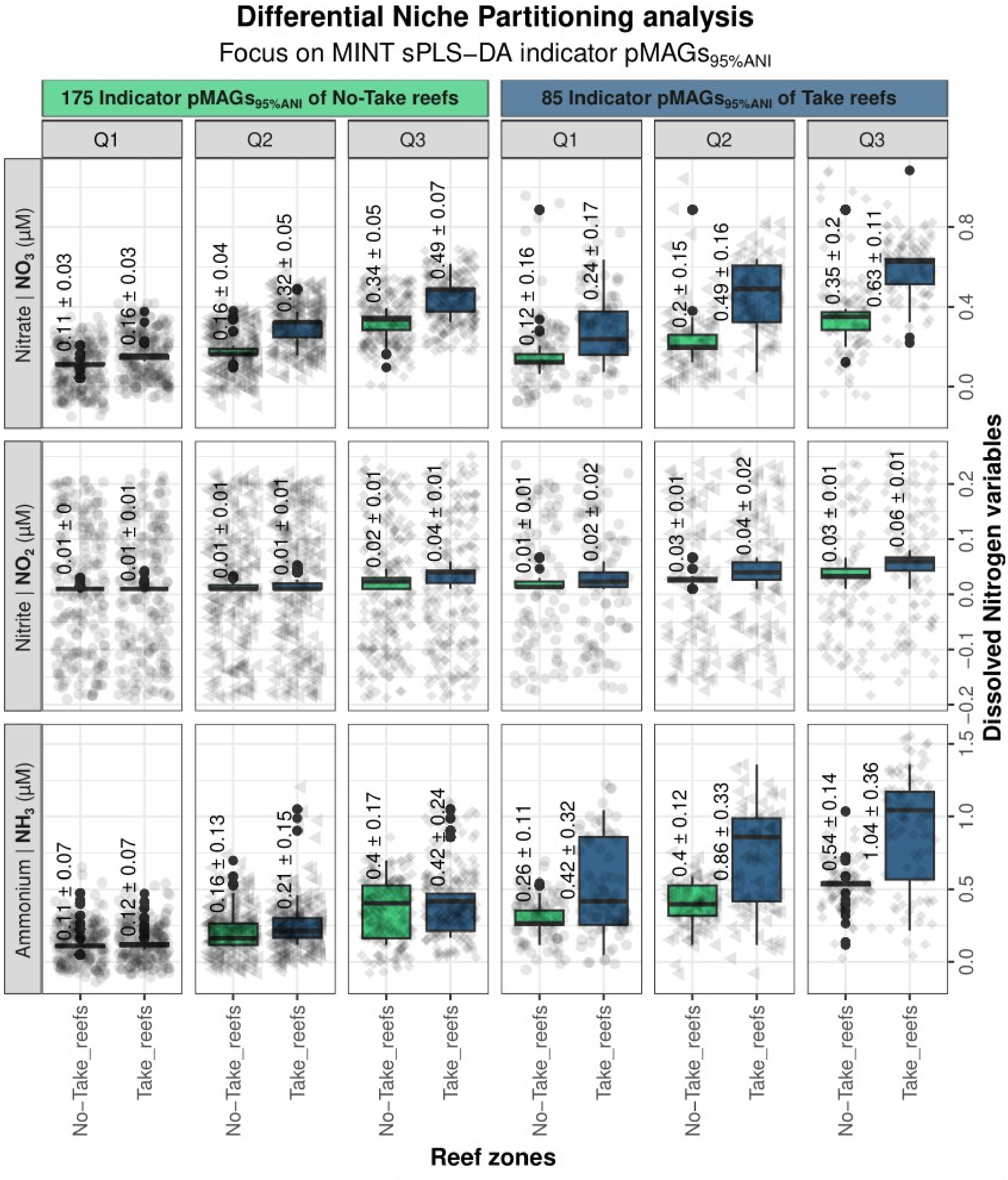


**Figure S43**. Comparative niche analysis for dissolved nitrogen variables between microbial indicators of fished reefs vs NTMRs. Boxplots show niche tolerance ranges (Q1: lower bound, Q2: optimum, Q3: upper bound) for the 236 microbial indicators of NTMRs (left) and 114 indicators enriched in fished reefs (right) —for ammonium (bottom), nitrite (middle), and nitrate (top). Niche bound values are visualised using distinct point shapes. This plot shows that microbial indicators of fished reefs have a preference towards higher values of all dissolved Nitrogen variables in fished reefs.

### **R**andom **f**orest (**RF**) models | Predictions of continuous environmental variables from microbial abundances

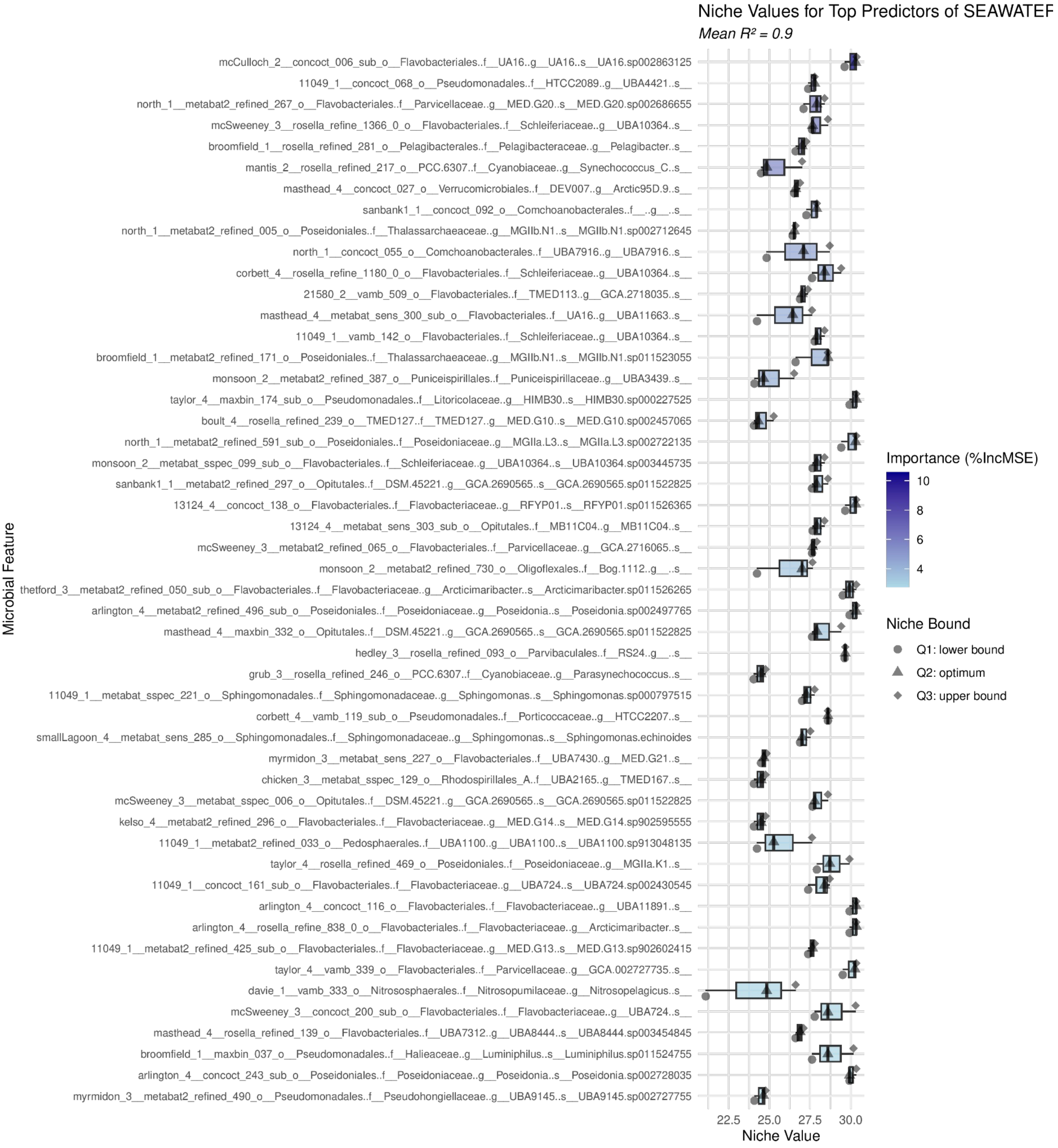


**Figure S44. Microbial predictors of surface seawater temperature (SST)**. Boxplots show niche tolerance ranges (Q1: lower bound, Q2: optimum, Q3: upper bound) for the top 50 microbial predictors per pMAG for SST. Niche bound values are visualised using distinct point shapes: circles (Q1), triangles (Q2), and squares (Q3). In addition, microbial predictors are additionally colored by random forest importance (%IncMSE), using a light-to-dark blue gradient (light blue = low importance, dark blue = high importance).

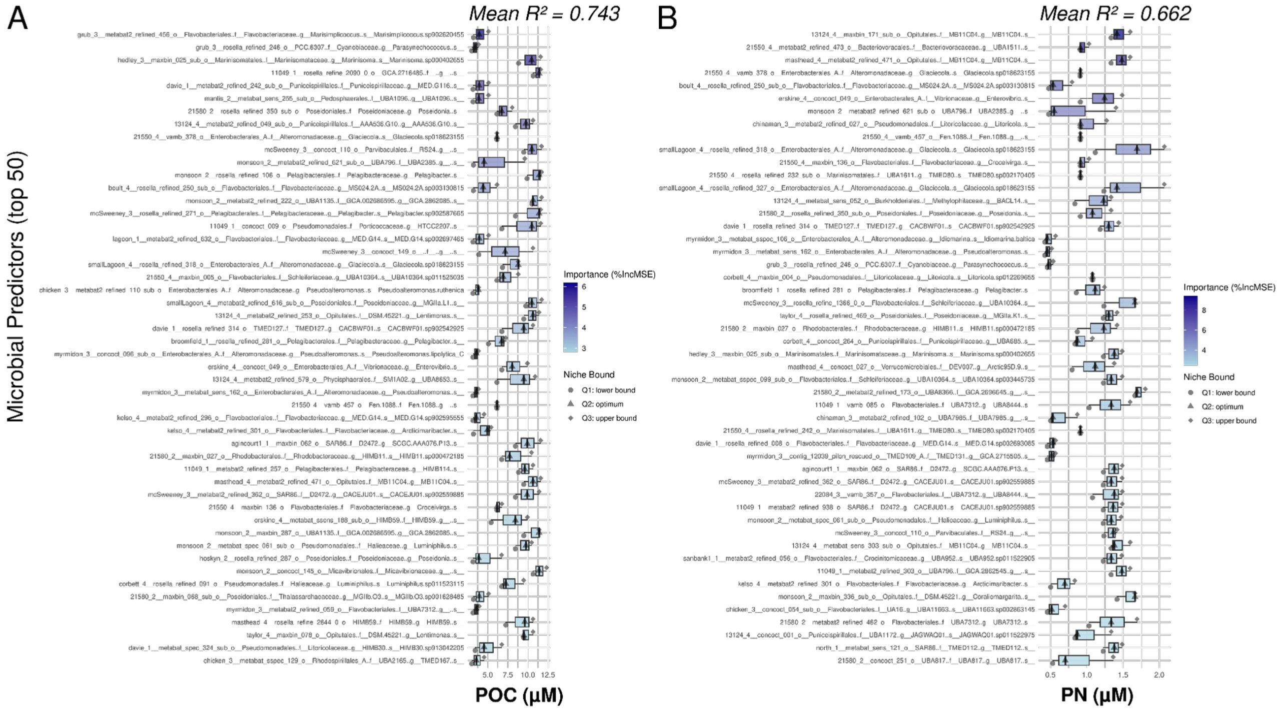


**Figure S45. Microbial predictors of particulate nutrients**. Boxplots show niche tolerance ranges (Q1: lower bound, Q2: optimum, Q3: upper bound) for the top 50 microbial predictors per pMAG for (**A**) Particulate organic carbon (POC) and (**B**) particulate nitrogen (PN). Niche bound values are visualised using distinct point shapes: circles (Q1), triangles (Q2), and squares (Q3). In addition, microbial predictors are additionally colored by random forest importance (%IncMSE), using a light-to-dark blue gradient (light blue = low importance, dark blue = high importance).

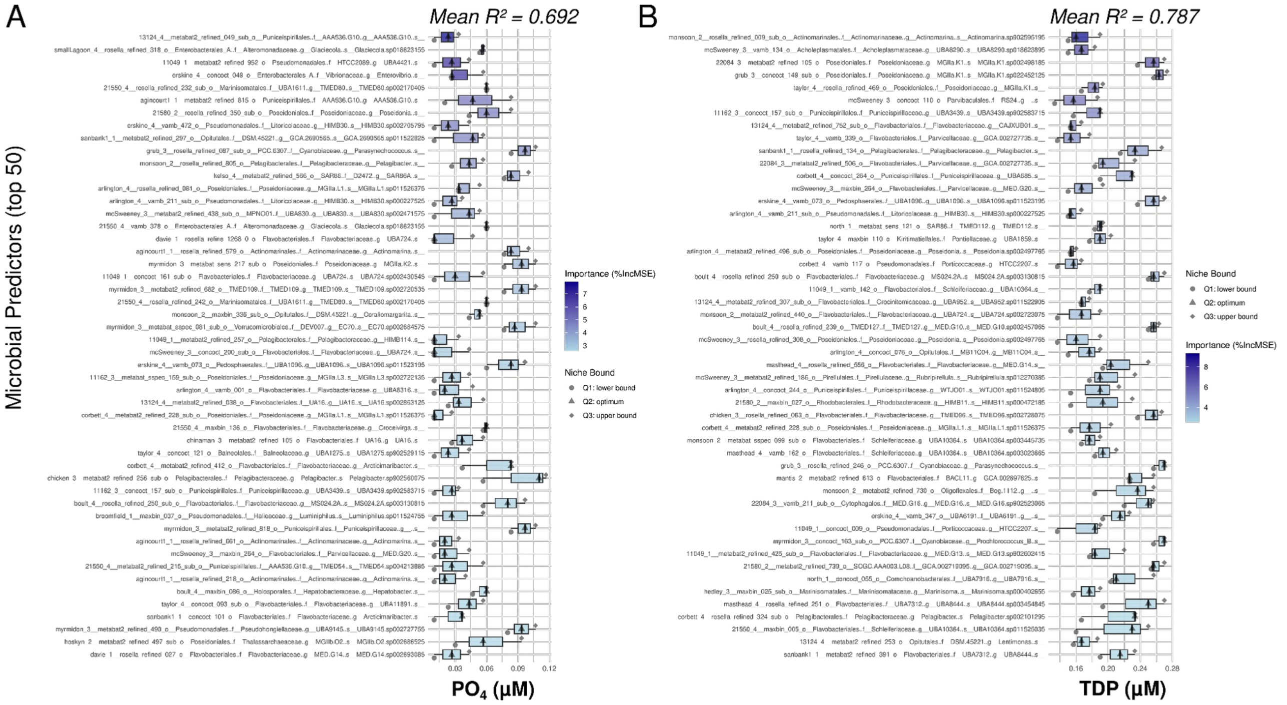


**Figure S46. Microbial predictors of dissolved phosphorus**. Boxplots show niche tolerance ranges (Q1: lower bound, Q2: optimum, Q3: upper bound) for the top 50 microbial predictors per pMAG for (A) phosphate ($PO_4^{3-}$) and (B) total dissolved phosphorus (TDP). Niche bound values are visualised using distinct point shapes: circles (Q1), triangles (Q2), and squares (Q3). In addition, microbial predictors are additionally colored by random forest importance (%IncMSE), using a light-to-dark blue gradient (light blue = low importance, dark blue = high importance).
